# Large-scale discovery and annotation of hidden substructure patterns in mass spectrometry profiles

**DOI:** 10.1101/2025.06.19.659491

**Authors:** Laura Rosina Torres Ortega, Jonas Dietrich, Joe Wandy, Hans Mol, Justin J. J. van der Hooft

**Affiliations:** Bioinformatics Group, Wageningen University & Research, Wageningen, the Netherlands; Wageningen Food Safety Research – part of Wageningen University & Research, Wageningen, the Netherlands; College of Medical, Veterinary & Life Sciences, University of Glasgow, Glasgow, United Kingdom; Department of Biochemistry, University of Johannesburg, Johannesburg 2006, South Africa

## Abstract

Untargeted mass spectrometry measures many spectra of unknown molecules. To annotate them, tandem mass spectrometry (MS/MS) generates fragmentation patterns representing common substructures. MS2LDA discovers these patterns via unsupervised topic modelling as Mass2Motifs. However, MS/MS-based substructure identification is limited by computational efficiency and interpretability. Here, we report an up to 14x speed improvement through improved algorithmic efficiency. Furthermore, the new automated Mass2Motif Annotation Guidance (MAG) aids in structurally identifying Mass2Motifs. Using three chemically diverse curated MotifDB-MotifSets for benchmarking, MAG achieved median substructure overlap scores of 0.75 up to 0.93, demonstrating robust substructural annotations. We further validated MS2LDA 2.0 in experimental data by identifying substructures of pesticides spiked into a biological matrix and demonstrated its discovery potential by annotating previously uncharacterized fungal natural products. Together with the new visualization app and MassQL-searchable MotifDB, we anticipate that MS2LDA 2.0 will boost the identification of novel chemistry and hidden patterns in mass spectrometry profiles.

## Main

Mass Spectrometry (MS) is an essential tool used across diverse fields, such as metabolomics, food safety, natural products, and drug discovery. Its high sensitivity and specificity make it essential for identifying and characterizing small molecules and biomolecules in complex samples^1^. Tandem mass spectrometry (MS/MS), a key advancement first developed in proteomics^2^, has transformed metabolomics since its introduction into the field^3^. By fragmenting detected molecular ions into smaller ions, MS/MS provides richer structural information. Furthermore, MS/MS has become critical in organizing^4^ and annotating^5^ mass spectrometry-based metabolomics profiles.

Despite advances in instrumentation and the rapid growth of public MS/MS libraries^6,7^, a substantial fraction of MS/MS spectra remains unannotated, often less than 15%, depending on the sample type^8^. To close this gap, several computational approaches have been proposed, ranging from propagating structural annotations through molecular networking^4^, via fragmentation tree-based annotation methods like SIRIUS^9^, to deep learning methods matching spectra to structures such as DiffMS^10^, MIST^11^, and MSNovelist^12^. Despite these efforts, annotating metabolite features on the basis of their MS/MS spectra remains difficult due to their complex fragmentation behavior, limited mass spectral libraries, and the diversity of chemical space^13,14^.

Especially when full structural elucidation is not feasible, identifying substructures can still provide meaningful chemical insight, such as by detecting novel bioactive scaffolds or by revealing new relationships between molecules. Substructure-based tools analyze fragmentation patterns to detect molecular building blocks^15^. One of the first of these was MS2LDA^15^, an unsupervised method based on topic modelling, which identifies recurrent patterns of fragments and losses, termed Mass2Motifs. These Mass2Motifs are manually annotated (for example, a neutral loss of 162 is associated with a hexose substitution, *e.g.*, glucose), and because such substructures recur across molecules, the same Mass2Motif can be present in multiple spectra. The unsupervised nature of MS2LDA enables the discovery of such recurring patterns without relying on predefined substructure libraries, making it particularly valuable for exploratory data analysis.

Since its conception in 2015, MS2LDA has grown into a widely adopted tool for substructure discovery in MS/MS data. Over time, it has evolved from a standalone algorithm into a more user-friendly and integrated platform. A web application was introduced to explore Mass2Motifs^16^, and a database with manually annotated and curated Mass2Motifs, named MotifDB, was built to enable researchers to reuse and share substructure annotations^17^. MS2LDA was also integrated with other metabolomics tools, such as the widely-used GNPS platform^4^, allowing researchers to find relationships between molecules based on shared substructures and not only on mass spectral similarity^18,19^. These developments have made MS2LDA a widely-used tool, for instance in natural product discovery^20^, in studying microbial diversity^21,22^, and in xenobiotic screening^23^.

Despite its advancements, several key challenges remain. First, extracting Mass2Motifs from large datasets (over 20,000 spectra) is computationally demanding, taking many hours to compute. Second, the interpretation of Mass2Motifs still depends heavily on manual annotation by human experts. While other tools such as MESSAR^24^ automate parts of this process, they rely on predefined libraries and fixed features and ignore peak intensities. Third, MS2LDA lacks integration with widely used Python-based metabolomics frameworks such as matchms^25,26^ which did not exist at the time of its initial development.

To address these limitations, MS2LDA 2.0 introduces a completely redesigned and scalable framework for substructure exploration in mass spectrometry. Alongside a more computationally efficient and performant topic modeling algorithm, it brings several key innovations: a novel automated Mass2Motif Annotation Guidance (MAG) system powered by Spec2Vec^27^, a fully rebuilt, queryable MotifDB leveraging MassQL^28^, full integration with Python-based metabolomics tools through matchms, and a completely redesigned interactive visualization app (MS2LDAViz 2.0) that can run both locally and on a server.

In this study, we validate MS2LDA 2.0 through benchmarking of manually curated MotifSets, demonstrate its application in analyzing spiked-in pesticides in food, and showcase its discovery potential in experimental datasets and large-scale MS/MS mass spectral libraries. We also highlight its integration into modern mass spectrometry analysis frameworks. To facilitate MS2LDA 2.0’s uptake by the community, an online MS2LDAViz 2.0 instance is running on https://ms2lda.org where two case study results can be directly loaded into the app to inspect and analyze the results.

## Results

The completely redesigned unsupervised substructure discovery tool MS2LDA 2.0 handles increasing dataset sizes. However, this results in larger collections of mass spectral patterns, Mass2Motifs, that need to be structurally annotated to use them to their full potential. To streamline this process, we developed the automated Mass2Motif Annotation Guidance (MAG) and the Mass2Motif-matching functionality by Spec2Vec^27^ against MotifDB. First, we evaluated the performance of MAG against manually annotated Mass2Motifs and subsequently demonstrate its capabilities using three different case studies with growing complexity: from spiked-in molecules in a complex matrix, via fungal natural product discovery, to a large set of >75,000 heterogeneous library spectra. Finally, we describe the complete MS2LDA 2.0 framework, including the new MassQL-powered MotifDB and MS2LDA 2.0 app to visualize, inspect, and annotate results.

### Automated Mass2Motif Annotation Guidance

When redesigning MS2LDA 2.0, we identified the manual annotation of Mass2Motifs as a substantial bottleneck, making automated approaches necessary. Even for an expert familiar with the data, manually annotating 30 Mass2Motifs, as performed in the original MS2LDA paper, was a laborious task requiring multiple days of work. As our current tool can easily discover hundreds to more than a thousand Mass2Motifs, manual structural annotation has become a substantial bottleneck. To address this challenge, we developed Mass2Motif Annotation Guidance (MAG), an automatic procedure which aids in Mass2Motif annotation by using spectral similarity scoring with Spec2Vec and additional refinement steps (see Methods) to suggest possible structures containing the Mass2Motif of interest. Subsequently, the user can then recognize a co-occurring substructure in these structures that the Mass2Motif represents. If relevant structures are found, the structural annotation process becomes faster and easier.

To benchmark MAG, we used Mass2Motif datasets that have been previously manually annotated and published as three positive ionization mode MotifSets in MotifDB: urine-derived^23^, GNPS library-derived^17^, and MassBank library-derived^17^ Mass2Motifs. The evaluation was based on the overlap of MACCS fingerprints between manually and automatically recommended structures using a motif fingerprint (see Methods: Motif fingerprints). To benchmark, we introduce a substructure overlap score, hereafter termed SOS (Supplementary Notes 1.3), where an SOS of 1 represents a complete overlap between the automated recommendation and the manual annotation.

In most cases, MAG’s substructure predictions aligned well with manual annotations. For example, in the urine-derived Mass2Motifs MotifSet, MS2LDA 2.0 achieved an average SOS of 0.72 and a median score of 0.75 across 101 Mass2Motifs (Fig. 1). Of these, 20 Mass2Motifs showed a complete substructure match with manual annotations. This means that if users were analyzing these recommendations, they would have very relevant support from MAG for over 20 out of these 101 Mass2Motifs, and useful support for another 30 Mass2Motifs. Fig. 1 shows examples of Mass2Motifs with their respective substructure-overlap scores.

**Fig 1:**
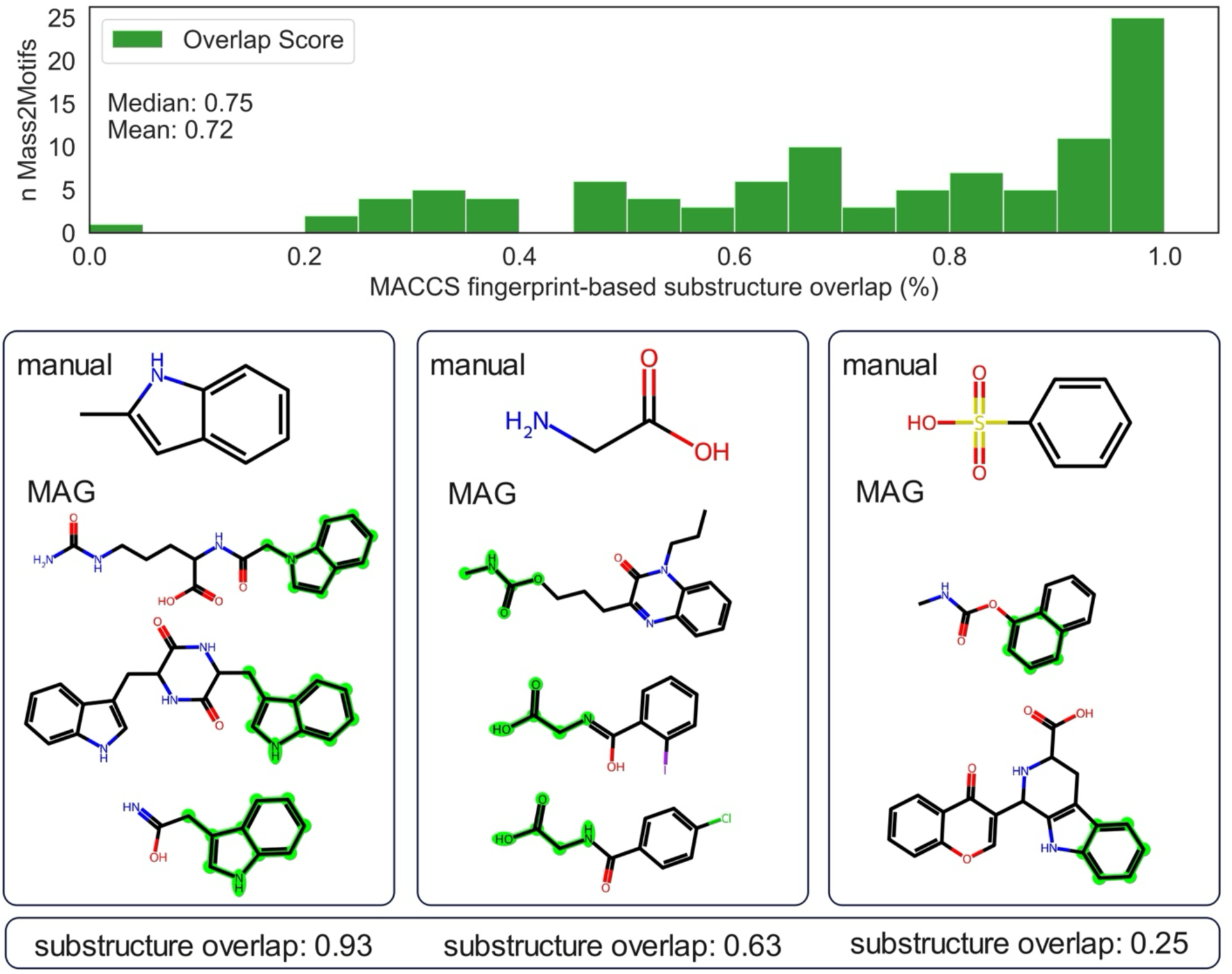
**Benchmarking Mass2Motif Annotation Guidance (MAG) on urine-derived Mass2Motifs.** Motif Annotation Guidance (MAG) was tested on an experimentally derived urine MotifSet with 101 manually annotated Mass2Motifs. The evaluation was based on MACCS fingerprint overlaps of the MAG and the manual annotated substructures. At the top a histogram shows the distribution of the alignment between manual and automated structural annotations and at the bottom three examples with perfect, good and bad substructure overlap scores (from left to right) are shown. The highlighted substructure (green) shows the structural overlap between manual and automated annotations.

While the urine-derived Mass2Motif MotifSet is based on experimental data, the GNPS and Massbank MotifSets are acquired from spectral library data. For these MotifSets, MAG achieved significantly higher substructure overlap scores, with an average of 0.85 (median 0.93) for the GNPS-based Mass2Motifs and an average score of 0.79 (median 0.93) for the Massbank-based Mass2Motifs. The discrepancy between experimental and library-based substructure overlap scores is likely caused by two main reasons. First, MAG makes use of the mass spectral libraries that have overlap with the two library-based MotifSets, thus likely containing more relevant spectral-structure combinations than for the urine experimental data. Second, the inclusion of more (unrelated) or noisy mass spectra in MS2LDA modeling of the experimental dataset compared to the selected and sometimes more cleaned spectra in libraries can lead to noisier Mass2Motifs that make automated structure annotation more difficult.

Overall, our results demonstrate that MAG provides reliable automatic guided annotations for positive ionization mode-generated Mass2Motifs, showing comparable results to manual annotations from MotifDB while making the process much faster and more efficient than the laborious completely manual process. By automating the annotation process, MAG enables efficient handling of the increasing number of Mass2Motifs generated by MS2LDA 2.0, facilitating the interpretation of large-scale experiments.

### Case Study on Pesticides

In our first case study, we consider spiked-in pesticides in a complex biological matrix derived from food. Identifying structures in food samples is a challenging process due to their complexity, which stems from the large number of different compounds present ^29,30^. To assess whether MS2LDA 2.0 can assist in rapidly identifying Mass2Motifs in a complex sample matrix, we spiked a tomato sample with a standard mixture of 211 pesticides (Supplementary Notes 2.3). This mixture contained various pesticide classes, including organophosphates, carbamates, triazole fungicides, sulfonylureas, and benzoylureas.

MS2LDA 2.0 successfully detected Mass2Motifs that correspond to recurrent substructures of the pesticides present in the spiked mixture (Fig. 2). We focus here on three Mass2Motifs with coherent MAGs that could be linked to distinct pesticide classes: organophosphates, benzoylureas and sulfonylureas. Two additional pesticide-related Mass2Motifs are presented in Supplementary Notes 2.1. To link the spiked compounds to the Mass2Motifs, we used the probability calculated by LDA during the modelling process, where 0 represents no likely association and 1 a very likely association between a spiked compound and a Mass2Motif.

**Fig. 2:**
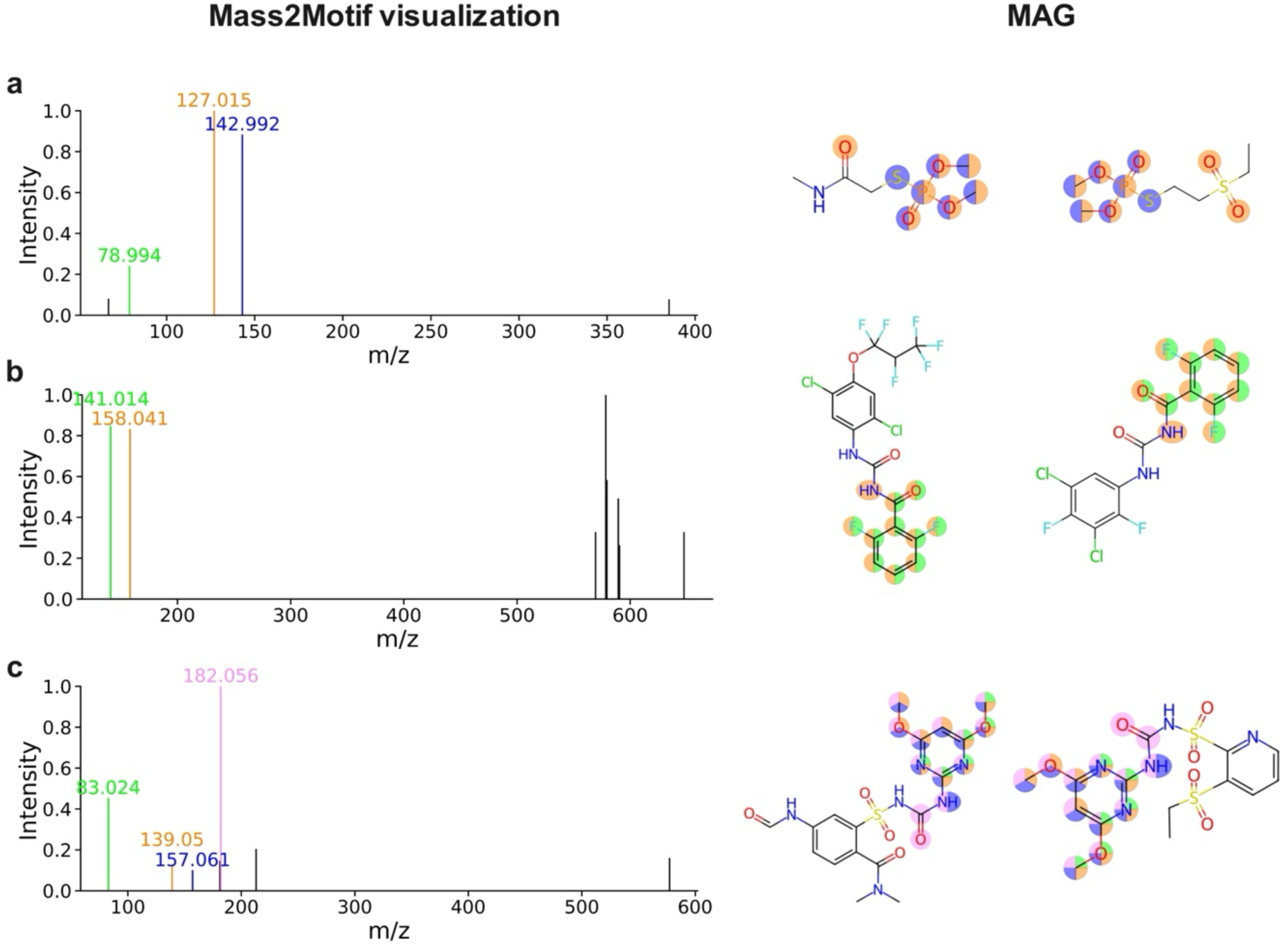
**Identified Mass2Motifs related to Spiked-in Pesticides.** a) Mass2Motif 6 shows three fragments related to the MAG of organophosphates. The color of the fragments in the Mass2Motif visualization relates to the highlighted atoms in the MAG and visualizes possible fragmentation pathways of the molecule. b) Two highlighted fragments from Mass2Motif 90 represent a benzoylurea substructure as shown by MAG annotations. All other fragments above m/z 500 are not directly related to the benzoylurea substructure represented by the Mass2Motif. c) Mass2Motif 236 shows four common fragments in the Mass2Motif visualization. The MAG suggests a sulfonylurea substructure. While the fragments do not include the sulfonyl group, they are linked to different fragmentation pathways in the 4,6-dimethoxypyrimidine substructure.

Organophosphates (OPs) are a widely used pesticide class primarily in crop protection^31^ and act as potent cholinesterase inhibitors^32^. Mass2Motif 6 (Fig. 2a) is defined by the mass fragments *m/z* 127.015, 142.992 and 78.994, representing a dimethylated (mono- or di-thio)-phosphate substructure. This substructure was prevalent in 9 out of 17 spiked OPs: 4 display a strong association with the Mass2Motif (probability > 0.7), two a moderate association (> 0.3) and two a weak association (< 0.1) (Supplementary Notes 2.2). The latter two spiked compounds still share two of the three key fragments, indicating that complementary fragment-based queries (e.g., through MassQL) could retrieve them. OPs lacking the dimethylated phosphate group follow alternative fragmentation pathways and are therefore not captured by this Mass2Motif.

Benzoylureas, another major pesticide class, function as insect growth regulators by inhibiting chitin synthases^33^. Mass2Motif 90 (Fig. 2b), characterized by fragments at *m*/*z* 158.041 and 141.015, shows a fragmentation pathway typical of benzoylureas, including the loss of an isocyanate group followed by subsequent NH₃ loss^34^. In the spiked mixture, three benzoylureas, along with one oxazole and one isoxazole sharing the typical fragmentation pathways, were present. As we used a complex biological matrix, MS/MS spectra were acquired for three of them. Of these, two benzoylureas showed sufficient association with Mass2Motif 90 (probability > 0.3) to be identified. The third compound did not associate with the Mass2Motif, likely due to the presence of additional unrelated fragments (e.g., *m/z* 578.41 and 579.41) in the Mass2Motif. As these fragments are only represented in one of the spiked compounds, the generalization of the Mass2Motif decreases. This issue can be circumvented by using the optimized fragments of the Mass2Motif (Supplementary Notes 1.2).

Sulfonylureas, which interfere with the biosynthesis of specific amino acids in plants^35^, were also identified in the dataset. The core structure of dimethoxylated sulfonylureas was associated with Mass2Motif 236 (Fig. 2c), which contained four key fragments: *m/z* 83.024, 139.05, 157.061, and 182.056. Overall, 15 compounds belonging to the sulfonylurea class were spiked. Out of the 15 compound 6 compounds included a dimethoxylated pyrimidine core of which 5 could be identified with a moderate to high certainty (probability > 0.5). Only one compound showed a low probability of 0.1, but the corresponding spectrum contained all four highlighted fragments, showcasing the specificity of Mass2Motifs. The low probability can be explained by the spiked compound having other more abundant fragments not part of the Mass2Motif. Finally, also in this case, a MassQL-like search for the identified fragments in the Mass2Motif would be able to identify all spiked compounds with the dimethoxylated sulfonylureas core structure.

Our findings demonstrate that MS2LDA 2.0 effectively identifies substructures related to pesticides of different classes within complex food matrices, thereby further demonstrating the effectiveness of the combination of MS2LDA 2.0 and MAG to discover and structurally annotate spiked-in bioactive scaffolds. We note that the ability of MS2LDA 2.0 to identify Mass2Motifs depends on the frequency and specificity of a fragmentation pattern and the collection of MS/MS spectra of sufficient signal-to-noise. In our case study, MS2LDA 2.0 was effective at experimental annotations of frequent and specific pesticide related Mass2Motifs. This reinforces its potential application in nontargeted screening of pesticides and their metabolites as well as transformation products.

### Case Study on Natural Products

The second case study explores the fungal specialized metabolome. Natural product extracts, such as those obtained from edible mushrooms, are complex metabolite mixtures that can contain bioactive compounds with potential therapeutic or toxicological effects. In their study, Khatib et al. characterized three molecular families in metabolomics profiles of two edible fungal extracts: erinacerins, hericenones, and ergostanes^36^. However, these metabolite families could only be annotated using SNAP-MS^37^ and DEREPLICATOR+^38^, approaches like GNPS library matching or SIRIUS did not provide annotations for these molecules. Here, we assess MS2LDA 2.0’s ability to confirm and extend these annotations through substructure-level analysis.

Fist, we mapped Mass2Motifs to the previously identified molecular families by Khatib et al.^36^ using MolNetEnhancer^19^ to connect the molecules by their Mass2Motifs. Second, we queried the identified Mass2Motifs in MotifDB using MassQL and conducted Mass2Motifmatching against MotifDB. MS2LDA 2.0 inferred the 200 Mass2Motifs in just 1.2 minutes, using an 8 GB RAM machine, rapidly identifying recurring substructural Mass2Motifs across the dataset of 2714 features.

The erinacerin molecular family contains seven Mass2Motifs, with the three most frequently shared ones highlighted in Fig. 3a. Mass2Motif 137 is shared among 19 of 23 nodes in this family, which was identified by MAG as a 4-[4-(2-chlorophenyl)piperazin-1yl]-4-oxobut-2-enoic acid (Fig. 3a & b), containing the substructure-pyridine ring present in erinacerin N. Another frequent motif was Mass2Motif 22, which represents another substructure present in this molecular family identified as 4-methylpentanoic acid (Fig. 3b, Supplementary Figure S22 in Section 3.3 in detail). Both substructures are characteristic of erinacerin N (C19H28N2O6) related structural features. For Mass2Motif 137, we could find a useful hit as verapamil related-structure in MotifDB via MassQL (S.I. Section 3.3 Table S7). For Mass2Motif 22 no results were retrieved from MotifDB. Another member of the erinacerin family, erinacerin F (C24H31NO7), contains Mass2Motif 108, which corresponds to a 4,6-dihydroxy-2,3-dihydro-1H-isoindol-1-one structure (Fig. 3b). Related indole-like structures were retrieved from MotifDB via MassQL queries, supporting these substructure assignments (S.I. Section 3.3, Table S8).

**Fig. 3:**
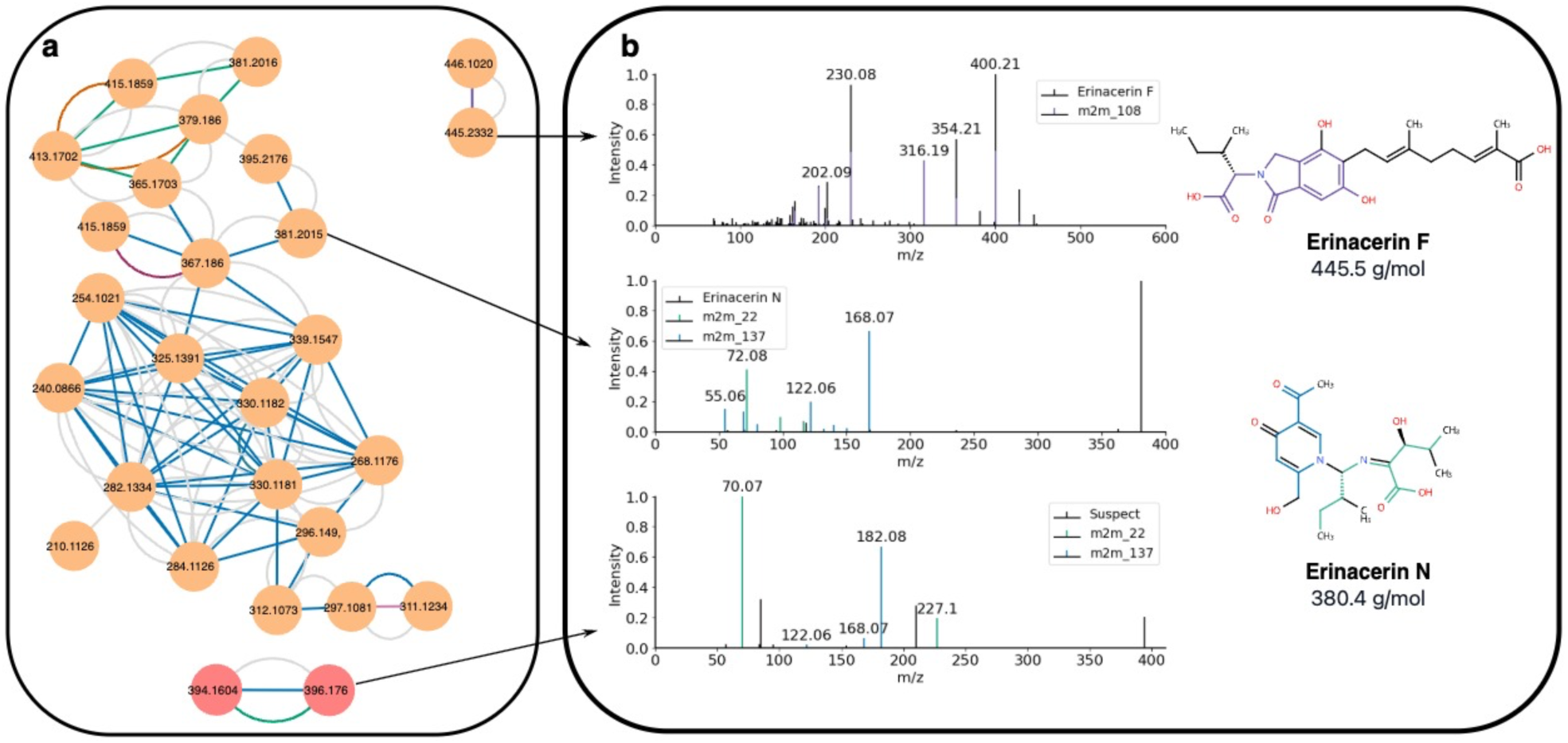
**MS2LDA 2.0 results from the fungal natural product case study.** a) Molecular familes of erinacerins, the erinacerin N molecular family being the largest (23 orange nodes, top left), erinacerin F (2 orange nodes, top right) and suspect erinacerin molecular family (2 pink nodes, bottom) reported by Khatib et al.^36^. The grey edges represent cosine similarity, colored edges represent Mass2Motifs, generated using MolNetEnhancer^19^. b) Spectra of the molecules manually annotated as erinacerin F and N (top) and the suspect molecule (bottom). Mass2Motifs related fragments colored on top. Structures of the complete erinacerin molecules with the inferred Mass2Motifs by MAG colored (right).

Beyond confirming substructures in reported erinacerins by Khatib et al., MS2LDA 2.0 uncovered two previously unannotated spectra with precursor *m*/*z* of 396.176 and 394.160, respectively, which are not clustered in this molecular family (Fig. 3a, pink nodes). The nodes shared key Mass2Motifs (137 and 22) and their mass differences relative to erinacerin N suggest possible derivatives, *e.g.*, addition of NH (15 Da), with a proposed molecular formula of C_19_H_29_N_3_O_6._ In the suspect molecular family (Fig. 3a, pink nodes), the node of *m/z* 396.176, contains Mass2Motif 137 and 22 with probabilities of 0.105 and 0.208, and for the precursor of *m/z* 394.160, the same motifs show 0.451 and 0.38 probabilities, respectively. These findings mean that Mass2Motif 137 and 22 are main topics contributing to the probabilities in both nodes, especially for the node with a *m/z* of 394.160. These compounds were not annotated by any of the previously used tools, illustrating MS2LDA 2.0’s capacity to complement existing tools in natural product dereplication.

In the hericenone molecular family, MS2LDA 2.0 identified three key Mass2Motifs across 20 nodes. Mass2Motif 163 and 186 (present in 15 and 5 nodes, respectively) were annotated both as benzodihydropyran substructures by MAG (S.I. Section 3.1 Figs S12 and S13), while Mass2Motif 90 (found in 12 nodes) was annotated as a 2-(2-methylbut2-enamido)acetic moiety (S.I. Section 3.1 Fig. S14). Via MassQL querying, we did not obtain any hits for Mass2Motif 163. Mass2Motif 186 retrieved terpenoid-like Mass2Motifs for its annotation (S.I. Section 3.1 Table S3). For Mass2Motif 90, MassQL retrieved terpene, sterone and related structures; a contrasting result compared to its MAG annotation (S.I. Section 3.1 Table S4 and Fig S14). These results could be explained by the fact that Mass2Motif 90 shows only one top fragment, when querying against MotifDB via MassQL, there is a high probability of retrieving unrelated molecules sharing the same fragment.

For the ergostane molecular family, MS2LDA 2.0 uncovered six recurring Mass2Motifs (S.I. Section 3.2 Fig. S15a). The dominant motifs included Mass2Motif 173, representing a steroid backbone and a carbon side chain, and Mass2Motif 193, both containing a steroid scaffold by MAG recommendation (S.I. Section 3.2 Figs S16 and S19), which correlates with the molecules manually annotated by Khatib et al. (S.I. Section 3.2 Fig. S15b). Mass2Motif 21 is also frequent for this family, annotated as 2,15-dihydroxy-7methyl-6-oxabicyclo[11.3.0]hexadeca-3,11-dien-5-one, which is structurally not very similar to the original structure proposed but they share the five-carbon fused ring (S.I. Section 3.2 Fig. S17). The MassQL results for the Mass2Motifs of the nodes previously reported did not yield additional structural insights for this molecular family, only Mass2Motif 0 results retrieved a diverse class of motif-related structures such as depsidones, quinones and paraconic acids, which is not aligning with MAG nor the manual annotation (S.I. Section 3.2 Fig. S18).

Finally, we computed the Mass2Motif-matching functionality between our experimental data and the following MotifDB-MotifSets: GNPS, MassBank and LDB (Lichen DataBase) MotifDB POS (Supplementary Notes 3.4). Several top-ranking, with a threshold over 0.7, supported the substructure proposed by MAG (S.I. Section 3.4 Table S10). For instance, Mass2Motif 10 matched Mass2Motif 13 and 11 with scores of 0.77 from Massbank and GNPS MotifSets respectively, both annotated as “Alkyl aromatic substructure indicative for aromatic ring”, which correlates with the MAG results shown in S.I. Section 3.4 Fig. S23, this highlights the value of Mass2Motif similarity screening as complementary strategy for substructure annotation. Another two positive examples with other structural moieties are Mass2Motif 126 and Mass2Motif 61 (S.I. Section 3.4 Figs S24 and S25).

MAG and Mass2Motif-matching annotations do not align for all the cases, for example for Mass2Motif 156 (S.I. Section 3.4 Fig. S26), the MAG annotation is 3-propanoyloxy-4(trimethylazaniumyl)butanoate, which does not align with the manual annotation as “Fragments indicative of a glycosylation”, however, when we inspect the motif (S.I. Section 3.4 Fig. S27), it reveals a loss of 161.1 *m/z*, as an indicative of a glycosylation. For motifs with high similarity, manual annotated motifs can offer a sufficient annotation, however, most of the spectra do not show high similarity with the MotifSet manually annotated Mass2Motifs. For instance, spectra similarity above 0.7 retrieved only 10 results with manually annotated motifs of the total 2714 spectra. Moreover, all these results are duplicated motifs between GNPS and MassBank. These two MotifSets share many Mass2Motifs, giving us only 5 unique results for this dataset. This reveals the importance of other motif annotation strategies developed in this study such as MAG and MassQL approaches.

Altogether, this case study demonstrates how MS2LDA 2.0 is a redesigned framework with multiple strategies that allows rapid, informative substructural annotation of experimental MS/MS data, enabling the identification of substructures related to molecular families beyond traditional spectral clustering methods.

### Case Study Suspect List

The third and final case study deals with a large and heterogeneous collection of mass spectra. We applied MS2LDA 2.0 on 77,218 MS/MS spectra representing the positive mode subset of the recently published suspect spectral list containing mass spectra retrieved from publicly available sources annotated through molecular networking^39^. The annotation of the suspects consists of an InChI of a known compound plus or minus the mass difference between the known compound and the suspect. Hence, all annotations are based on structural analogues and do not represent full structural annotations. Given its size and composition, the suspect library provides a valuable use case to test MS2LDA 2.0 to explore its large-scale analysis capabilities on a set of heterogenous mass spectra of various structures generated on different instruments and with various collision energies by different contributors, all deposited to public GNPS mass spectral libraries.

Overall, MS2LDA 2.0 identified meaningful Mass2Motifs within the suspect spectral library, as inferred from comparing suspect annotations with MAG suggestions. The 1,500 generated Mass2Motifs were manually analyzed by examining their correspondence to each suspect. Here, we selected four examples where MS2LDA 2.0 shows a good alignment with the proposed suspect. To further verify the structural annotations, we used the mass spectral prediction tool CFM-ID 4.0^40^ (details Supplementary Notes 4.1-4.4) to assess alignments between the suspect and the Mass2Motifs. To highlight the presence of the Mass2Motif across the suspect library, we retrieved the number of compounds for which this Mass2Motif had the highest association based on LDA calculated probabilities. In Fig. 4a and 4b, both suspects are complementarily explained by two Mass2Motifs, while in Fig. 4c and 4d one Mass2Motifs explains a major part of the substructure of the suspects.

**Fig. 4:**
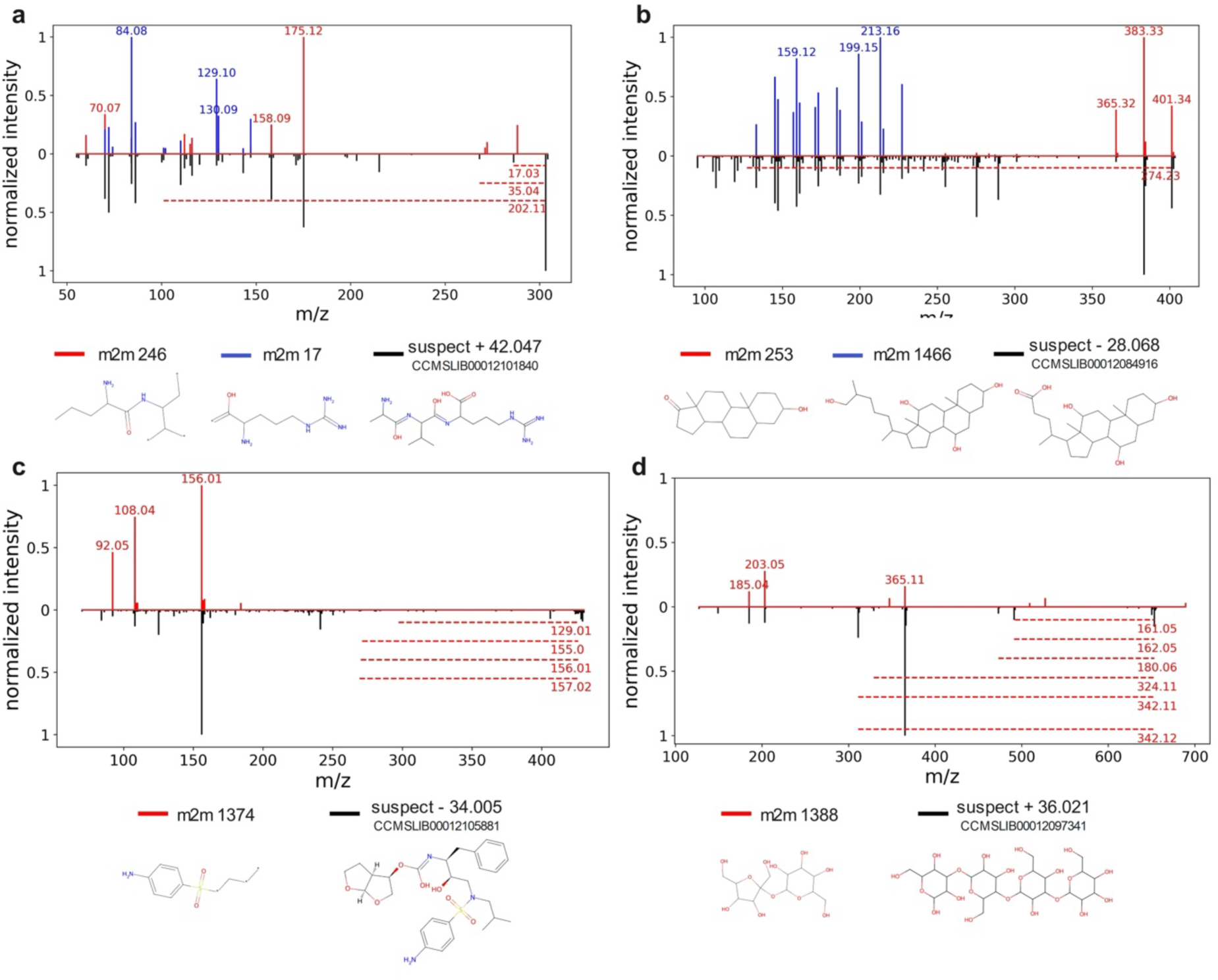
**Large scale Mass2Motif exploration with the GNPS spectral suspect list.** Four examples (a-d) show Mass2Motifs that represent substructures of identified suspects. The blue and red highlighted fragments represent the corresponding Mass2Motifs, while mirrored spectra in black represent the related suspect spectra. selected. The dotted horizontal lines represent losses belonging to the same colored Mass2Motifs. Below each mirror-plot, the MAG based recommendation (left, under the Mass2Motif (m2m) color identifier) as well as manual suspect reported by Bittremieux et. al. (right, under the suspect color indicator) are shown.

In Fig. 4a, a suspect that is structurally similar to alanyl-valyl-arginine could be associated with two Mass2Motifs that represent two different building blocks. Mass2Motif 246 is associated to the tris-amino substructure through the fragments *m*/*z* 60.06 and 175.12. Mass2Motif 246 and Mass2Motif 17 overlap with *m*/*z* 70.07 and 72.08 describing an alkylamine substructure. Furthermore, a loss of 17.03 in Mass2Motif 246 indicates the presence of a free amino group as it is present in the tris-amino substructure. Mass2Motif 17 mainly focuses on the amide substructure of arginine through fragment *m*/*z* 143.12. Mass2Motif 246 was the most dominant feature in 126 spectra, and Mass2Motif 17 in 311 spectra.

Fig. 4b shows two Mass2Motifs that together describe a bile-acid related suspect. Mass2Motif 1466 focuses on the decomposition of the core steroid structure following the fragmentation patterns of androstane or metenolone alike structures^41^. Mass2Motif 253 is mainly defined by three fragments with *m*/*z* values above 350 (365.32, 383.33, 401.34), which have a *m*/*z* difference of 18.01, with 365.32 being annotated as a bile-acid core structure by CFM-ID 4.0. Therefore, these fragments probably describe a fragmentation pathway within the bile-acid conjugation side or at the core steroid structure losing two water molecules. Mass2Motif 253 was dominant in 63 spectra and Mass2Motif 1466 in 481.

In Fig. 4c, Mass2Motif 1374 describes a sulfonamide substructure, which is a building block of a suspect associated with Darunavir. The main fragments describe the fragmentation of the sulfonamide substructure from the suspect (*m*/*z* 156.01) and the further decomposition of the sulfonamide (*m*/*z* 108.04 and *m*/*z* 92.05). Mass2Motif 1374 was dominant in 96 spectra.

Fig. 4d shows a suspect associated with 3-alpha,4-beta,3-alpha-galactotetraose, supported by at least two fragments with relatively high abundance. The difference between *m*/*z* 185.04 and *m*/*z* 203.05 indicates a loss of water, while additionally, several losses including 162.05 (hexose) further highlight how this Mass2Motif is related to the presence of conjugated sugar moieties including hexoses like glucose or galactose. Mass2Motif 1388 was dominant in 90 spectra.

These examples show how Mass2Motifs can describe a diverse range of molecules such as peptides (arginine-related, Fig. 4a), steroids (bile-acid-related, Fig. 4b), sulfonamides (Fig. 4c), and glycosides (Fig. 4d). This highlights the capability of MS2LDA 2.0 to handle large datasets and to find meaningful substructures across different structural classes. Without an exhaustive evaluation of all Mass2Motifs, we expect that many more novel identified Mass2Motifs patterns can be found by mining public mass spectrometry data.

Hence, there is a large potential for use of MS2LDA 2.0 on other large-scale MS/MS datasets, even with mass spectra from various sources such as the suspect library as we demonstrate here.

## Integration with other computational mass spectrometry frameworks and computationally e7icient topic modelling

In the re-design of MS2LDA 2.0, seamless integration with widely used Python-based frameworks for computational mass spectrometry was another main goal (Fig. 5a). The original version relied on custom Python code to process mass spectra and handle Mass2Motifs, limiting interoperability with community-standard tools. Instead, MS2LDA 2.0 has adopted standardized frameworks such as matchms^26^ to process mass spectra, including filtering, similarity calculations and data import/export, through well-maintained, modular components. This improves reproducibility, maintainability, and compatibility with other tools, especially those also built on the matchms framework.

**Fig 5:**
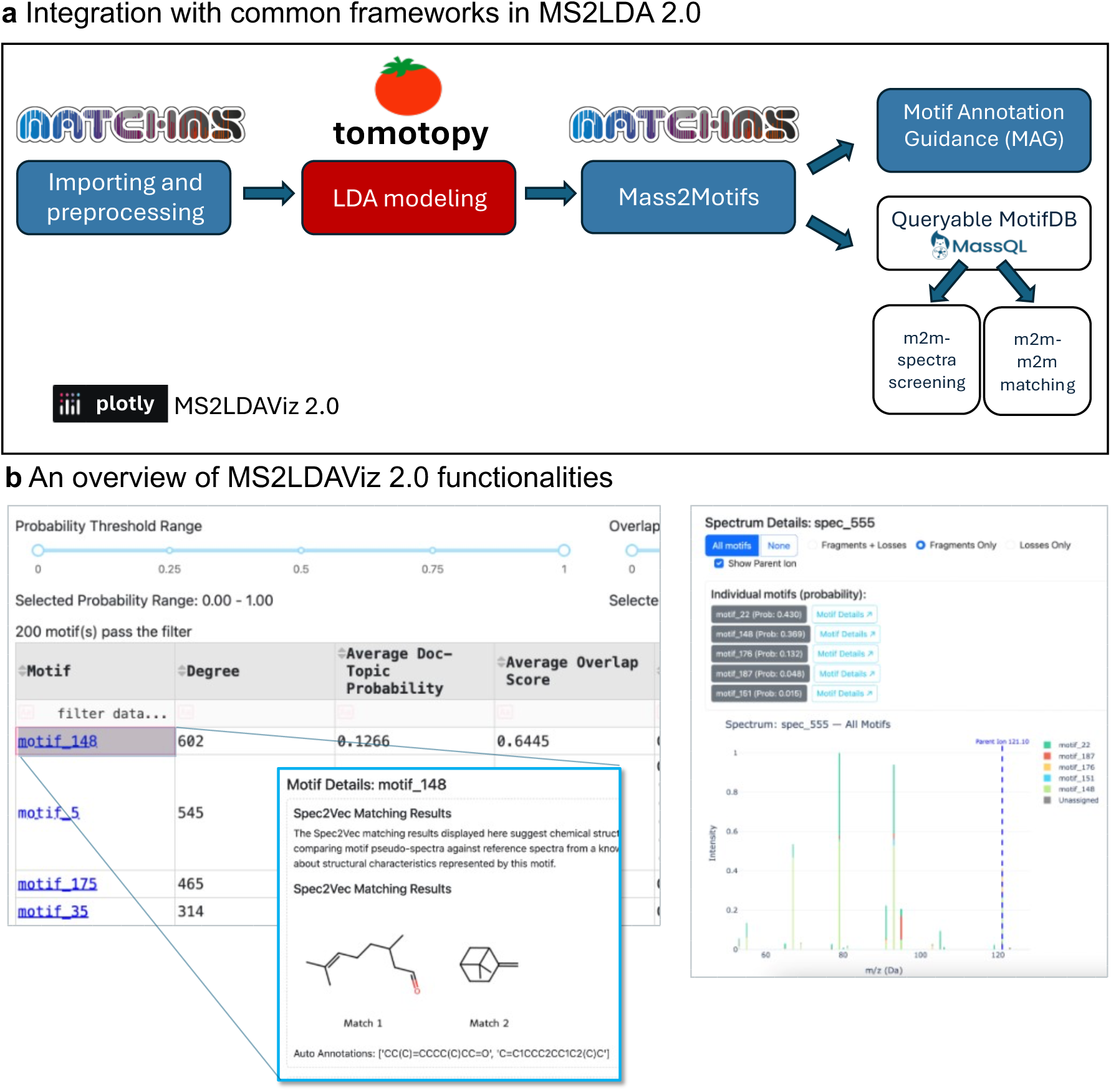
**Workflow and Tool Integration in MS2LDA 2.0.** b) Integration of MS2LDA 2.0 with common computational mass spectrometry frameworks. Mass spectrum importing and preprocessing are performed using matchms, while topic modeling is handled by the optimized LDA implementation provided by Tomotopy. The resulting Mass2Motif objects (Supplementary Notes 5.2) are now fully interoperable with multiple downstream tools, allowing seamless integration with automated Motif Annotation Guidance (MAG) and motif querying of MotifDB via MassQL. MotifDB can also be used for Mass2Motif-Mass2Motif (m2m-m2m) matching and Mass2Motif-spectra (m2m-spectra) screening. All these functionalities are now incorporated in an interactive MS2LDAViz 2.0 application. b) Overview of the MS2LDAViz 2.0 application functionalities. The left panel shows a ranked table of inferred Mass2Motifs along with automated structural annotations provided by MAG. Users can interactively inspect detailed annotations, including suggested chemical structures, by selecting individual motifs. The right panel illustrates how a single MS/MS spectrum can be decomposed into multiple contributing Mass2Motifs, a unique feature of the MS2LDA 2.0 approach that facilitates rapid interpretation of complex spectra.

MS2LDA 2.0 now utilizes Tomotopy for LDA inference^42^, a library written in C++ that supports parallelized inference using collapsed Gibbs sampling. This shift enables substantial speed improvements for the topic modelling: on an AMD EPYC 7532 32-Core Processor using 4 cores, MS2LDA 2.0 was 3x to 14x as fast as the previous version, which used a single-threaded batch variational Bayes implementation in NumPy. For example, discovering 400 Mass2Motifs in the GNPS Suspect-List library (87,899 spectra in total) required 3.46 hours with MS2LDA 2.0 on this hardware setup, compared to 20.51 hours with its predecessor (S.I. Section 3.2 Fig. S34). When reducing the number of cores and using a single core as the original MS2LDA version, the two versions showed similar speed (Fig. S33). This shows the importance of parallelized computing inference by Tomotopy, making large-scale analysis now feasible.

After MS2LDA 2.0 topic modelling, the resulting Mass2Motifs are now well integrated with matchms (Supplementary Notes 5.2), allowing integration with specialized downstream tasks. First, MAG utilizes Mass2Motifs as input for Spec2Vec^27^ to aid in the structural annotation of substructures. Based on the resulting annotations, structural fingerprints such as MACCS or Daylight can be generated, creating new annotation capabilities that are not possible with previously available workflows. Second, Mass2Motifs generated by MS2LDA 2.0 are now stored in a new standard file format based on the MassQL framework^28^. This enables flexible querying and retrieval of Mass2Motifs based on characteristic fragments and losses within MotifSets. Furthermore, the current setup allows researchers to easily contribute their own MotifSets generated with MS2LDA 2.0 to MotifDB, with the ultimate goal of making it an expanding community-driven motif annotation effort. Selected MotifDB-MotifSets generated with the initial MS2LDA version have been transferred into the new format to make them compatible with version 2.0.

The newly developed MS2LDAViz 2.0 app makes visualization of several aspects of Mass2Motifs easy and interactive (Fig. 5b). Previously, visualization was implemented using a Django-based web application, which requires considerable web development expertise, making it challenging for researchers without extensive software development backgrounds to install, modify, or extend upon. In contrast, the newly developed app is written in Dash, which offers a simpler Python-based approach. This makes it possible for data scientists and researchers to more easily visualize annotation output and to customize and extend the views themselves (Fig. 5b). We refer to the Methods and section 6 of the Supporting Information for more details on MS2LDAViz 2.0.

As a demonstration of seamless integration into existing analysis environments, we visualized molecular networks in Cytoscape using MolNetEnhancer^19^ in the *Case Study Natural Products*. This enabled the annotation of previously annotated as well as novel fungal metabolites from a fungal dataset, which was further supported by querying already discovered Mass2Motifs from MotifDB. **a** Integration with common frameworks in MS2LDA 2.0 **b** An overview of MS2LDAViz 2.0 functionalities

## Discussion

Untargeted metabolomics continues to face major challenges with structural elucidation, particularly when full structural elucidation is infeasible. However, in such cases, substructure-level analysis can provide meaningful chemical insights and drive hypothesis generation. The original MS2LDA introduced topic modelling for cofragmentation patterns (Mass2Motifs) from MS/MS data. While influential, it was limited by its high computational complexity, heavy reliance on full manual annotation, and fixed codes limiting integration with contemporary data-analysis frameworks.

Here we present MS2LDA 2.0, which offers a substantial improvement in performance over its predecessor. Due to its integration with other Python-based workflows, MS2LDA 2.0 offers multiple downstream functionalities once the Mass2Motifs are inferred. A central innovative functionality in MS2LDA 2.0 is the automated Motif Annotation Guidance (MAG), which uses Spec2Vec to match Mass2Motifs with known structures from mass spectral libraries. MAG provides an automated route to interpretation that complements human expert knowledge and allows researchers to prioritize motifs for further analysis leveraging public mass spectral library information.

In our benchmark, MAG achieved a strong performance across the three largest manually curated MotifSets from MotifDB, with median substructure overlap scores of 0.75, 0.93, and 0.93. Moreover, in experimental datasets, including complex pesticide-spiked tomato samples, MAG was able to retrieve various expected pesticide classes with high fidelity. Nonetheless, MAG’s performance depends on the availability and coverage of mass spectral reference libraries. For instance, in the fungal natural case study, some Mass2Motifs, such as Mass2Motif 90 associated to the hericenone family, provided a poor annotation (Fig. S15). We observed that annotation accuracy tends to increase with the number of peaks shared between the experimental Mass2Motif and the reference structure, as seen with Mass2Motif 193 and 108. MAG is not intended to replace human expert curation, but rather to complement it to accelerate interpretation, highlight likely candidates, and reduce the overall annotation burden.

Following up on the Mass2Motif annotation strategies, one important feature of the original MS2LDA was the ability to fix known topics from existing MotifSets directly during inference. This functionality helped the user to match their Mass2Motifs to the relevant MotifSets available in MotifDB. Although computationally less efficient, the original variational Bayes implementation was flexible enough to support this custom modification. In MS2LDA 2.0, this direct fixing of motifs during inference is no longer possible because it relies on the optimized but less customizable Tomotopy library. As an alternative, a similar outcome can now be achieved *post hoc* through spectral similarity comparisons. Because Mass2Motifs can now be processed as standardized spectrum-like objects via matchms, users can directly compare them to reference motifs after inference. This allows a more targeted analysis to reinforce structural hypotheses generated by MAG and manual review. This Mass2Motif-matching functionality for annotation was successfully demonstrated, for example, in the fungal natural product case study with known substructures such as lysine and phenylalanine (Supplementary Notes 3.4). However, similarity comparisons are not without limitations. Some Mass2Motifs may reflect overly general or noisy fragmentation patterns, leading to ambiguous matches. Further refinement of spectral similarity metrics and threshold parameters in Mass2Motifmatching remains an area for improvement and could enhance confidence in annotation, particularly when combined with other evidence sources. Nevertheless, our current implementation provides a flexible framework to match Mass2Motif patterns.

Another new key strategy to annotate Mass2Motiffs is by querying MotifDB via MassQL. While MS2LDA 2.0 can discover key mass spectral patterns in the data, MassQL provides a framework to accurately describe them and search them at scale in a reproducible manner. Here, we connected these two tools to provide an alternative annotation route to MAG. On the one hand, MassQL allows users to query the fragments and losses present in Mass2Motifs against curated entries in MotifDB. On the other hand, MS2LDA 2.0 can be used to infer representative patterns of a sample that later can be used in the overall MassQL ecosystem. Although not fully explored in this study, we encourage the community to use these tools in a complementary manner. In this work, we demonstrated the compatibility of MS2LDA 2.0 and MassQL in the fungal natural product dataset: annotations retrieved through MassQL-based screening were consistent with those suggested by MAG, highlighting the complementary potential of using these two approaches together in combination (Supplementary Notes 3).

To visually interact with all the previously existing and newly developed MS2LDA functionalities, including inspection of inferred motifs, MAG recommendation, Mass2Motif-matching, and MassQL querying, all these features are all brought together in the new MS2LDAViz 2.0 application. This Dash-based app enables user-friendly inspection and downstream analysis, allowing researchers to browse Mass2Motifs in the context of molecular networks, investigate MAG annotations, perform targeted motif searches, and generate hypotheses directly from MS/MS datasets without the need to write any code and with the benefit of visualizations to support decision-making.

Despite these advances, some limitations remain. As with the original MS2LDA, users must still predefine the number of Mass2Motifs (topics) prior to inference, and the model remains sensitive to hyperparameters (e.g., alpha and beta), which can affect motif quality. Besides performing a good estimation of the biochemical diversity and thus chemical scaffolds, substructures, and functional groups in the dataset, one strategy to mitigate this is to assess consistency across multiple models runs using different seeds or topic counts. Alternatively, nonparametric Bayesian models such as the Hierarchical Dirichlet Process (HDP)^43^ could infer topic numbers directly from the data. However, HDP is computationally intensive and can be difficult to tune for large-scale datasets, so it remains an avenue for future work to investigate its use for substructure discovery in metabolomics. In addition, some experimental Mass2Motifs, particularly those involving novel chemistry not represented in mass spectral libraries, still require full manual interpretation. Furthermore, MAG is currently working for positive ionization mode data; future community-based efforts to create a larger compiled negative ionization mode mass spectral library will also allow MAG to function for Mass2Motifs generated in negative ionization mode. While MS2LDA 2.0 facilitates much of the annotation process, human expert interpretation remains essential in these advanced cases. Nevertheless, when such Mass2Motifs are annotated with new substructures or scaffolds, the new MotifDB will facilitate their addition, and the MassQL integration supports searching for their presence across the entire body of public mass spectrometry data.

In summary, MS2LDA 2.0 integrates large-scale discovery and semi-automated annotation of Mass2Motifs within the broader metabolomics ecosystem. Its modular design and interoperability with community-driven resources like MotifDB and MassQL offer a practical path forward for substructure discovery in untargeted MS/MS workflows. As the field continues to expand and MotifDB grows, we anticipate that Mass2Motif annotation will become an increasingly routine part of high-throughput untargeted metabolomics analysis.

## Methods, Datasets, and Case Studies

We note that for all case studies and benchmarks MS2LDA 2.0 parameters need to be selected. Additionally, when running MS2LDA 2.0 a convergence curve of the LDA modelling process is generated to assess if the model was run for sufficient time. All selected parameters and generated convergence curves can be found in Supplementary Notes 7 and 8.

### MotifSets for MAG Benchmarking

The selection of suitable MotifSets for the Motif Annotation Guidance benchmarking was based on the number of annotated Mass2Motifs and their annotation confidence. We selected three MotifSets from MotifDB: urine-derived Mass2Motifs (101 Mass2Motifs)^23^, GNPS library-derived Mass2Motifs (73 Mass2Motifs)^15,17^, and MassBank library-derived Mass2Motifs (42 Mass2Motifs)^15,17^. These three MotifSets were chosen as they represent relatively large Mass2Motif collections whose annotations were reviewed with confidence. To convert the descriptive manner of Mass2Motif annotations into computer readable SMILES, every description was manually inspected and the SMILES for the described compound was retrieved from PubChem^44^.

### Case Study on Pesticides

We experimentally prepared a tomato sample spiked with 211 pesticides and performed a HPLC-MS/MS analysis. The names of all spiked compounds can be found in a separate file in the Supplementary Notes.

Sample preparation Case Study on Pesticides

First, 10 grams of blank tomato sample were transferred into a 50 ml Greiner tube. 10 ml of acetonitrile/1% formic acid were added and extracted in an overhead shaker for 30 min. Then, 4 grams of MgSO_4_ and 1 gram sodium acetate were added and the mixture was vortexed for 1 min to induce phase separation. Afterwards, the sample was centrifuged for 5 minutes at 3500 rpm. 250 µl of the upper layer of the extract was transferred into an LC vial, 25 µl of a 1000 ng/ml pesticide mixture containing 211 compounds and 225 µl of water were added. After mixing using a vortex, this vial was used for the LC-HRMS/MS analysis.

### LC and MS settings Case Study on Pesticides

Liquid chromatography was carried out on a Vanquish UHPLC system (Thermo Fischer Scientific). The LC eluents A and B were water and methanol both containing 2 mM ammonium formate and 0.1% formic acid. The gradient started with 100% phase A with a linear increase of phase B, which reached 100% at 15 min. For 6 min the eluent stayed at 100% of phase B and then increased to 100% phase A with a linear gradient within 1 min. The run stopped after 25 min. The flow rate was set to 0.3 mL/min and the injection volume was set to 5 µL. The column was a Waters UPLC BEH C18 1.7 um 2.1 mm x 100 mm with a column oven temperature of 50°C.

Mass spectrometric detection was carried out on a Thermo IQX Orbitrap instrument (Thermo Fisher Scientific) operated in positive ESI mode. Mass spectrometry was performed with a sheath gas flow rate of 48, auxiliary gas flow rate 11, and sweep gas flow rate 2. The spray voltage was set to 3400 V while the ion transfer tube temperature and the vaporizer temperature were set to 320°C and 350°C, respectively.

For MS1 the resolving power (R) of 120K was used, with a maximum injection time of 246 ms, an automated gain control (AGC) target of 400000, and RF lens of 70%. All data were acquired in centroid mode from a mass range from 100-1000 m/z. A dynamic exclusion filter was applied where compounds were excluded for 3s after the first appearance with a 10-ppm mass tolerance, also excluding isotopes.

An inclusion and exclusion list were used (see all compounds in S.I. Section 2.2 Table S2). The inclusion list contained 36 of the 211 spiked compounds and the exclusion list was automatically created based on the procedure blank run.

MS/MS data were acquired using the HCD cell for fragmentation with a normalized and stepped collision energy of 30/80%. The maximum injection time was 22 ms with an AGC target of 50000 and the Orbitrap resolution was set to 15K.

### Case Study on Fungal Natural Products

The natural product dataset was originally obtained from Khalib et. al.^36^. This dataset is publicly available and contains HPLC-MS/MS data measured in positive mode and data dependent acquisition mode for two fungal species: the mushrooms *Hericium erinaceus* and *Pleurotus eryngii.* We applied MS2LDA 2.0 on the dataset, thereby focusing on finding the correlations between Mass2Motfis and structures reported in the paper of three biochemically relevant metabolite groups that could not be annotated with GNPS library matching or SIRIUS in-silico annotation.

### Case Study using the Suspect List

The suspect list comprises a large collection of 87,899 mass spectra derived from published datasets in the form of suspects that are related to known library compounds^39^. The complete dataset was used to showcase MS2LDA 2.0 computational efficiency. As input for the case study, the 77,218 suspects measured in positive ionization mode were used to evaluate its automated annotation capabilities.

### Datasets used for Scalability Evaluation

To test the computational efficiency of MS2LDA 2.0, we downloaded the following libraries from the GNPS public libraries^4^: GNPS-NIH-NATURALPRODUCTSLIBRARY (1,267 spectra), GNPS-NIST14-MATCHES (5,185 spectra), GNPS-NIH-NATURALPRODUCTSLIBRARY_ROUND2_POSITIVE (7,377 spectra), BERKELEY-LAB (18,352 spectra), and the Suspect Library (87,899 spectra). All libraries were downloaded from the GNPS webpage. We calculated the time that the LDA inference took for the original MS2LDA implementation and version 2.0, thereby varying the number of discovered Mass2Motifs (100, 200, 300, 400). Since the algorithm of the new version is known to converge more slowly, the iterations were set to 1000 for MS2LDA 2.0 and 100 for the original MS2LDA. We note that the setting of 100 iterations for the initial version does not always lead to completely converged models, especially for larger datasets.

### Retraining Spec2Vec and Reference Database

A dataset combining publicly available positive ionization mode mass spectra from MoNA, MassBank, GNPS and Brungs et. al. library containing 502,685 spectra (dated 15/02/2025) was downloaded, preprocessed, and used to retrain Spec2Vec (details at https://github.com/vdhooftcompmet/MS2LDA/tree/main/notebooks/Spec2Vec), and it served as the reference library for the Motif Annotation Guidance. The mass spectral library dataset is publicly available from Zenodo (https://zenodo.org/records/12543129).

## Computational Methods

### MS2LDA 2.0 Framework and Implementation

MS2LDA 2.0 is a framework designed to identify substructures in tandem mass spectrometry (MS/MS) datasets. The workflow consists of five main steps: (a) spectrum loading and preprocessing, (b) topic modeling using Latent Dirichlet Allocation (LDA), (c) Mass2Motif annotation, (d) visualization, and (e) Mass2Motif storage and querying via MotifDB. MS2LDA 2.0 supports standard input formats such as .mgf, .mzML, and .msp. Mass spectrum processing, including ion peak filtering and metadata handling, is performed using the matchms library (v0.27.0). Mass fragment peaks and neutral losses are rounded to two or three significant digits (default: two) and converted to string representations (e.g., frag@70.04) to use for topic modeling. Peak intensities are discretized by scaling normalized intensity values (0-1) to integers (0-100), with each fragment or loss represented as repeated ‘words’ proportional to intensity. This ensures that more intense peaks have greater influence in the topic modeling, analogous to frequent words in text-based LDA.

LDA is performed using the tomotopy library (v0.13), producing Mass2Motifs as distributions over fragment and loss features. These features are then converted back to their original numerical values (e.g., frag@70.04 → 70.04), with topic-word probabilities (β values) used as normalized intensities in the final Mass2Motif representations. For further details on the underlying LDA method and its application to metabolomics data, we refer to the original MS2LDA paper by Van der Hooft *et al*.^15^.

A new web-based application, MS2LDAViz 2.0, has been developed using the Dash framework (v2.18). This app allows users to interactively explore Mass2Motif spectra, annotations, and Mass2Motif comparisons. Additionally, a command-line interface and Jupyter notebooks are provided for users who prefer scripted analyses or customizable interactive data exploration.

Mass2Motifs can be stored and queried using the MassQL library (v0.0.15). Queries are expressed using the MassQueryLanguage4Mass2Motifs syntax, an extension of MassQL specifically designed for searching Mass2Motifs (details in Supplementary Section 5b). These queries allow users to search against MotifDB, a community-driven repository of previously annotated MotifSets (see MotifDB Update and Integration), allowing efficient retrieval and comparison of Mass2Motifs across different datasets.

### Convergence Criteria

In the original MS2LDA, users were required to manually set the number of iterations for training the LDA model. In the updated version, MS2LDA 2.0 introduces early stopping, allowing the training to terminate automatically based on user-set convergence criteria. Users can specify a convergence threshold evaluated against one of four available metrics: perplexity, maximum likelihood, document-topic entropy, or document-word entropy. At defined intervals (default step size = 50 iterations), the relative change in the selected metric is calculated over recent checkpoints (default window size = 500 iterations). If this relative change falls below the specified threshold, training stops early. Otherwise, training continues until the maximum number of iterations is reached. Examples of convergence plots illustrating this process are provided in the Supplementary Notes 7.

### Motif Annotation Guidance (MAG)

A detailed example of how MAG identifies structures for annotation guidance can be found in the Supplementary Notes 1.1 and 1.2. Here, we provide a brief overview of its main innovations leading to the automated MAG. First, Spec2Vec^27^ is used to compute embeddings for each Mass2Motif as well as the embeddings for compounds in the reference database. Then, cosine similarity between embeddings is efficiently calculated using the FAISS library^45^. By default, the top five database hits are retrieved as candidate matches. However, depending on the database content, fewer relevant hits may be returned.

To filter out irrelevant database hits that may represent different substructures, we apply a masking strategy. Masking involves systematically removing one fragment or loss at a time from the Mass2Motif and recalculating its embedding using Spec2Vec. For example, given a Mass2Motif containing fragments 70.04, 91.03, and 125.69, each fragment is masked in turn, resulting in three modified motifs: one without 70.04, one without 91.03, and one without 125.69. A new embedding is generated for each masked variant, and its cosine similarity is recalculated against the initially retrieved database hits.

The idea behind masking is that database hits corresponding to the same substructure will exhibit a similar change in similarity scores when key fragments or losses are masked. For instance, if four of the five hits share a substructure associated with fragments 70.04 and 125.69, their similarity scores are expected to decrease when those fragments are masked. In contrast, a fifth hit representing a different substructure (primarily associated with fragment 91.03) will be affected differently.

This behavior is identified by applying agglomerative clustering (scikit-learn version 1.5.1) to the resulting cosine similarity matrix from all masked variants. Ideally, clustering isolates the hits that represent the same substructure. The cluster that includes the best original database hit is then selected to guide MAG.

### Mass2Motif Fingerprints

MS2LDA 2.0 creates fingerprints to represent possible substructures in molecules. These fingerprints are based on results from the Motif Annotation Guidance (MAG). MAG works by identifying a set of candidate chemical structures that are likely to share a common substructure and belong to the same Mass2Motif.

To build a fingerprint that captures just this shared substructure, MS2LDA 2.0 first generates a fingerprint for each candidate structure. Then, it compares them and keeps only the bits (chemical features) that appear in at least 80% of the fingerprints (default threshold). This way, the final fingerprint highlights the features most likely to belong to the shared substructure.

### MotifDB Update and Integration

The updated version of MotifDB enables storing, querying, and retrieving annotated Mass2Motifs. MotifDB stores Mass2Motifs in an extended MassQL format (version 0.15.0), adapted to handle differences between spectra and Mass2Motifs. One substantial difference is the explicit storage of losses, as these cannot be derived directly *from the “spectrum-like object”* for Mass2Motifs due to the absence of precursor ions. Additionally, since Mass2Motifs lacks MS1-level data and retention times, certain MassQL functionalities, such as MS1-level or retention-time-based queries, are not supported in MotifDB. Additionally, we extended the current format by supporting *the* stor*age of* additional metadata which can be found in Supplementary Notes 5.3). *This additional metadata was inspired by the original MotifDB and was found helpful to select relevant MotifSets to match Mass2Motifs to, for example based on shared origin.* The metadata can also be queried using the new METAFILTER:metadata=value syntax from the new visualization interface. All Mass2Motifs generated by MS2LDA 2.0 are automatically stored in this MotifDB-compatible format. Furthermore, the current MotifDB is stored within the MS2LDA 2.0 GitHub repository (https://github.com/vdhooftcompmet/MS2LDA/tree/main/MS2LDA/MotifDB), making it easier to version and trace changes over time, and it is downloaded directly when installing MS2LDA 2.0.

### Visualization

A new web-based application has been developed to allow users to visualize discovered Mass2Motifs and inspect screening results. This new MS2LDAViz 2.0 application replaces the previously deployed application at ms2lda.org^16^, which is no longer actively maintained due to computational efficiency issues and its customized code structure. By contrast, the new visualization leverages Dash^46^, offering a simpler development syntax and a more straightforward deployment on remote servers or local machines. Compared to Django, Dash has a lower barrier to entry and is more widely familiar to data scientists, making it easier for the metabolomics community to contribute and expand visualization features. To facilitate the uptake of MS2LDA by the community and allow for quick inspection of MS2LDA models by visualizing them, we have launched a demo version on ms2lda.org. Users can also load in the pesticide and natural products case study results to browse through the results presented in this study.

With the new MS2LDAViz 2.0, users now also have the option to install the application locally, increasing accessibility and flexibility across different research scenarios. In the local application, there is also a graphical user interface (GUI) available to run MS2LDA, guiding users through the parameter settings. A high-level overview of the functionalities in the new application is shown in Fig. 6b with a more detailed overview being available in Supplementary Notes 6.1-6.3. Ultimately, we envision MS2LDAViz 2.0 to become a community-driven resource enabling users to browse, annotate, and contribute annotated Mass2Motifs back to MotifDB, fostering collaboration in substructure discovery.

### Mass2Motif-Mass2Motif Matching

One functionality from the initial version that is no longer available in MS2LDA 2.0 is the use of a MotifDB-MotifSet as fixed topics during the LDA model learning. Previously, fixed topics allowed predefined Mass2Motifs to be directly identified within new datasets by combining the discovery of “free” (unsupervised) and “fixed” (supervised) topics, the latter with manually curated structural annotations, allowing for the knowledge from previously annotated analyses to be transferred to a new analysis. This functionality is omitted in the current release due to a limitation in Tomotopy, which does not support exact fixing of previously discovered topics. As an alternative, we allow users to screen for previously characterized motifs by computing cosine similarities between Spec2Vec embeddings of new Mass2Motifs and existing MotifSets. These MotifSets may originate from prior experiments or from MotifDB. This screening approach differs from the Motif Annotation Guidance (MAG) described above, and is complementary, as its primary purpose is to rediscover previously annotated motifs through Spec2Vec matching, without explicitly verifying the importance of individual fragments (Supplementary Notes 6.4).

## Data availability

The experimental fungal dataset used in this study is publicly available from sources mentioned in the original publication by Khatib et al.^36^ The pesticide dataset is available from GNPS-MassIVE (MSV000097937). Results including all Mass2Motifs and their annotations, which were generated for both datasets are available from Zenodo (DOI: https://zenodo.org/records/15857387), and as notebooks in the GitHub repository (https://github.com/vdhooftcompmet/MS2LDA/tree/main/notebooks/Paper_results).

## Code availability

The MS2LDA 2.0 framework, including all modules and scripts for reproducing the results presented in this paper, is available at https://github.com/vdhooftcompmet/MS2LDA and every release is archived on Zenodo (DOI for version 2.0: https://doi.org/10.5281/zenodo.15858124; a persistent DOI to the latest version: https://zenodo.org/records/15858124) under the MIT License. The software can be easily installed via PyPI using ‘pip install ms2ldà -see for other installation option the GitHub Readme. Files to retrain Spec2Vec models are archived and versioned on Zenodo (DOI: https://zenodo.org/records/12543129). Comprehensive installation instructions and usage guides for MS2LDA 2.0 and its visualization interface MS2LDAViz 2.0 are provided in the repository’s README file.

## Acknowledgements

The authors thank Niek de Jonge, Esteban Charria Girón, and Dick de Ridder for proofreading the manuscript. Furthermore, we thank Paul Zomer for the assistance with measuring and setting up the pesticide case study.

## Author Contributions

RTO, JD, JW, and JJJvdH conceptualized the project idea. RTO and JD contributed equally to software development, data analysis, and writing the manuscript. JW contributed to software development and writing the manuscript. HM contributed to the pesticides case study. JJJvdH supervised the project, contributed to data analysis, and writing the manuscript. All authors edited and approved the final manuscript.

