## Supplementary Information for "Large-scale discovery and annotation of hidden substructure patterns in mass spectrometry profiles"

### 1. Automated Mass2Motif Annotation Guidance and Benchmarking

#### 1.1 The three steps of Mass2Motif Annotation Guidance

One of the core innovations of MS2LDA 2.0 is its ability to automatically guide the annotation of Mass2Motifs by suggesting structures that are likely to contain the detected substructure of interest.

The following paragraph describes the three steps that lead to a Mass2Motif Annotation Guidance (MAG): a library search with Spec2Vec[1,2], masking of fragments and losses, and finally a clustering of structural candidates based on the preceding steps. To illustrate MAG we use a Mass2Motif from the case study on pesticides (Fig. S1).

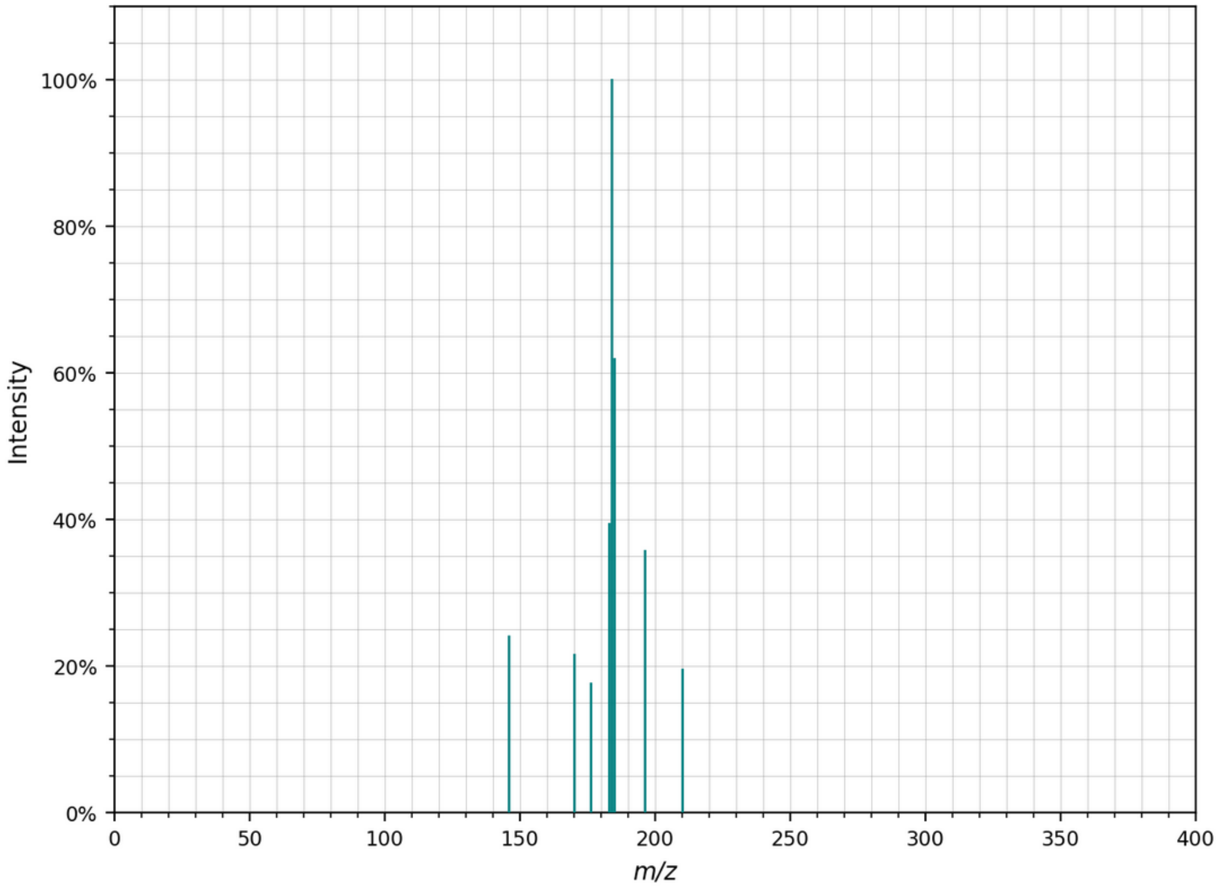

**Fig. S1.** *Mass2Motif 219 from the case study on pesticides consists of nine mass fragments and one neutral loss (not shown; has no importance in this example).*

The first step of MAG uses Spec2Vec to retrieve the most similar spectra between the Mass2Motif and a reference database containing over 500,000 spectra (see Methods section for details). The number of similar library spectra retrieved can be configured, with the default set to 10. If the task were to find full structural matches, Spec2Vec similarity scores above 0.7 or higher would be considered valuable and reliable information for an annotation. However, for substructure annotation, the scores are generally lower (often between 0.3 and 0.7) and less reliable. This reduced reliability also affects the relative ranking of spectra, meaning that a score of 0.62 may indicate a better substructure match than a score of 0.64.

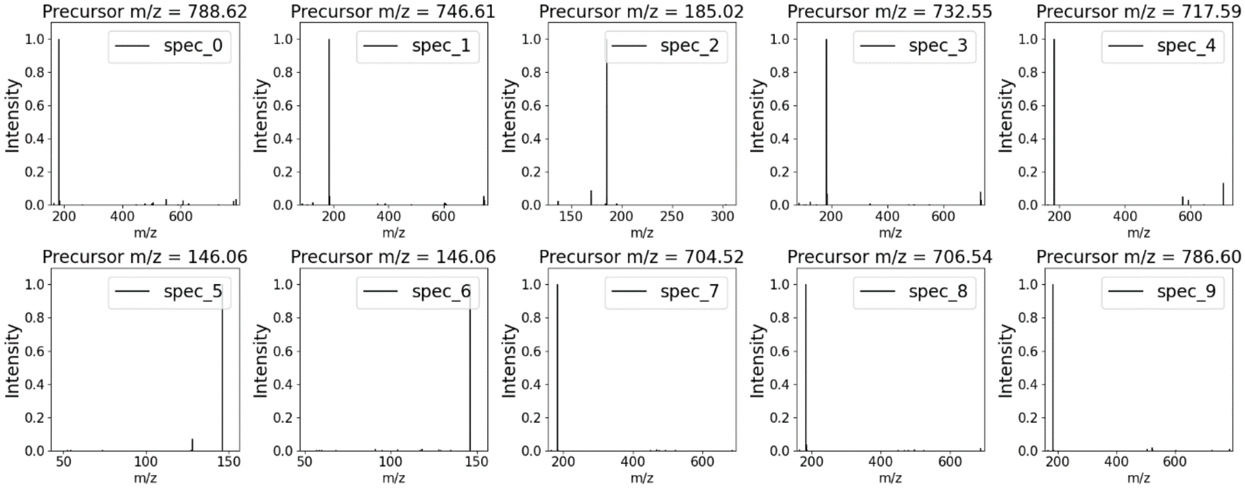

**Fig. S2.** *Ten retrieved spectra based on the highest Spec2Vec scores (0.38-0.45) between a Mass2Motif and the reference database. The similarity score is the highest in the top left corner and the lowest in the bottom right corner.*

The top ten spectra most similar to the Mass2Motif, based on Spec2Vec similarity, differ in terms of their base peaks (Fig S2). To algorithmically detect such anomalies within the top ten spectra, a masking procedure is applied.

Masking is a procedure analogous to backward elimination: each fragment or loss feature is temporarily removed (masked), and the Spec2Vec similarity is recalculated. This approach helps localize the fragments and losses that contribute most to the structure matches and allows for clustering based on these important features. In our implementation, all fragments and losses are iteratively replaced with -1, and the Spec2Vec similarity between the masked Mass2Motif and each spectrum is recalculated. This results in a pattern suitable for clustering (Fig. S3).

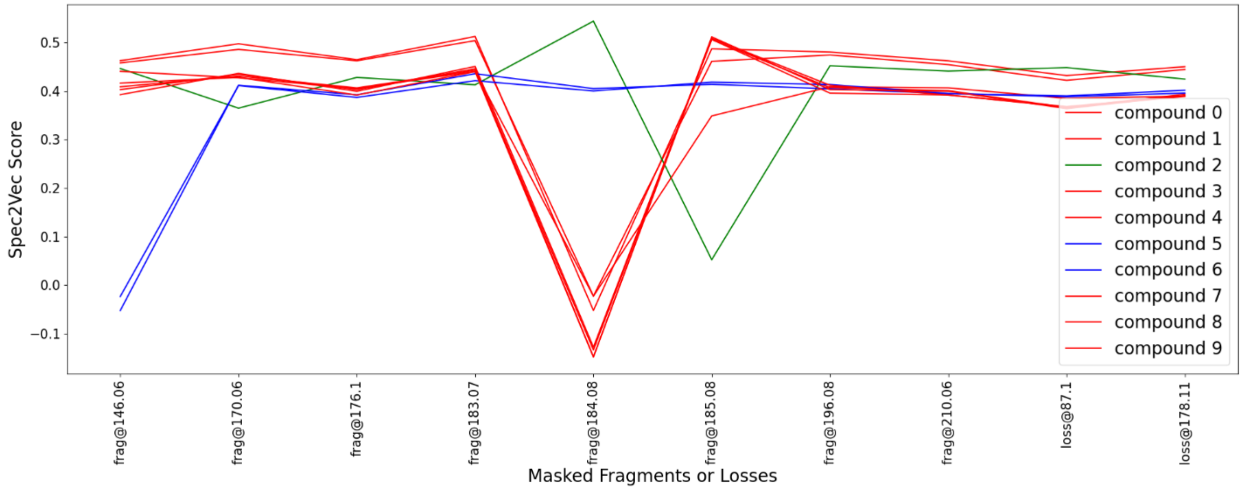

**Fig. S3.** *The visualization of the masking results shows the effect on the Spec2Vec score for the top ten compound when a fragment or loss is removed from the Mass2Motif. The Figure already includes the clustering based on the masking, which splits the ten compounds in three clusters highlighted by different colors.*

Fig. S3 shows the result of agglomerative clustering, which identified three distinct clusters. The largest cluster (red) containing 7 compounds is recommended by MAG. Upon examining the structures behind the spectra, we see that MAG correctly clustered compound 2 (cluster 2), compounds 5 and 6 (cluster 3), and compounds 0, 1, 3, 4, 7, 8, and 9 (cluster 1), according to structural similarity (Fig. S4).

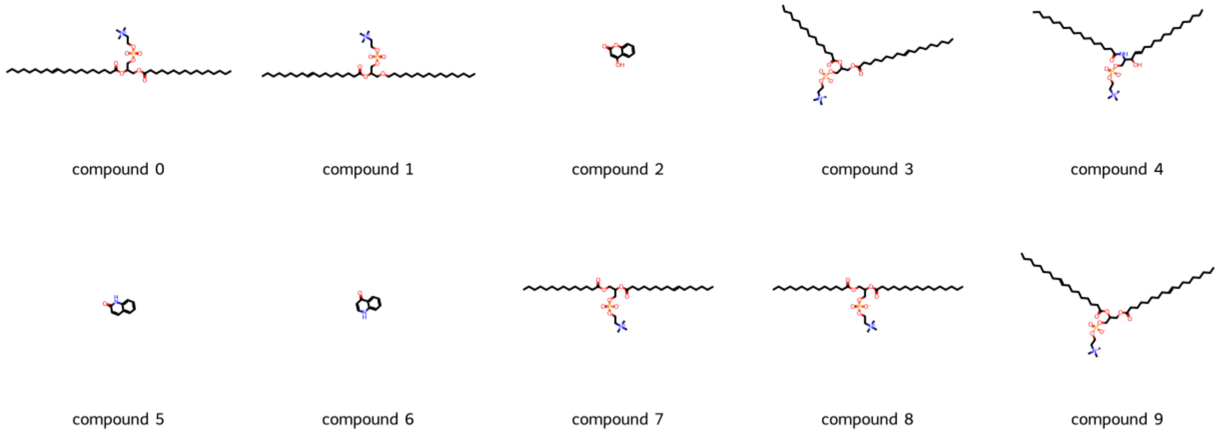

**Fig. S4**. *Structural annotation of the top ten retrieved compounds from the reference database after computing Spec2Vec.*

#### 1.2 Mass2Motif Optimization

As shown in Fig. S4, Spec2Vec can sometimes suggest a group of chemical structures that do not share meaningful substructure similarities—for example, compound 1 and compound 2 (Fig. S4). This happens because the algorithm bases its suggestions solely on observed fragments and losses, which can be misleading.

MAG attempts to address this by clustering more similar compounds together. However, even within a single Mass2Motif, unrelated fragments or losses can still be present. These may result from noise or artifacts in the dataset, or from genuinely new substructures that are not included in any reference database. If such novel substructures are expected, manual inspection and annotation of the Mass2Motif becomes necessary.

In many cases, however, the problematic fragments are not truly novel but stem from misassignments made by the LDA algorithm—for instance, due to unique side chains or dataset-specific features. To refine the results, we remove all fragments that are not consistently found across the recommended group of structures. The optimized Mass2Motif then retains only the fragments shared among the clustered compounds. As illustrated in Fig. S5, only one fragment is common to compounds 0, 1, 3, 4, 7, 8, and 9—implying that the other fragments are likely unrelated to the core substructure shared by this group.

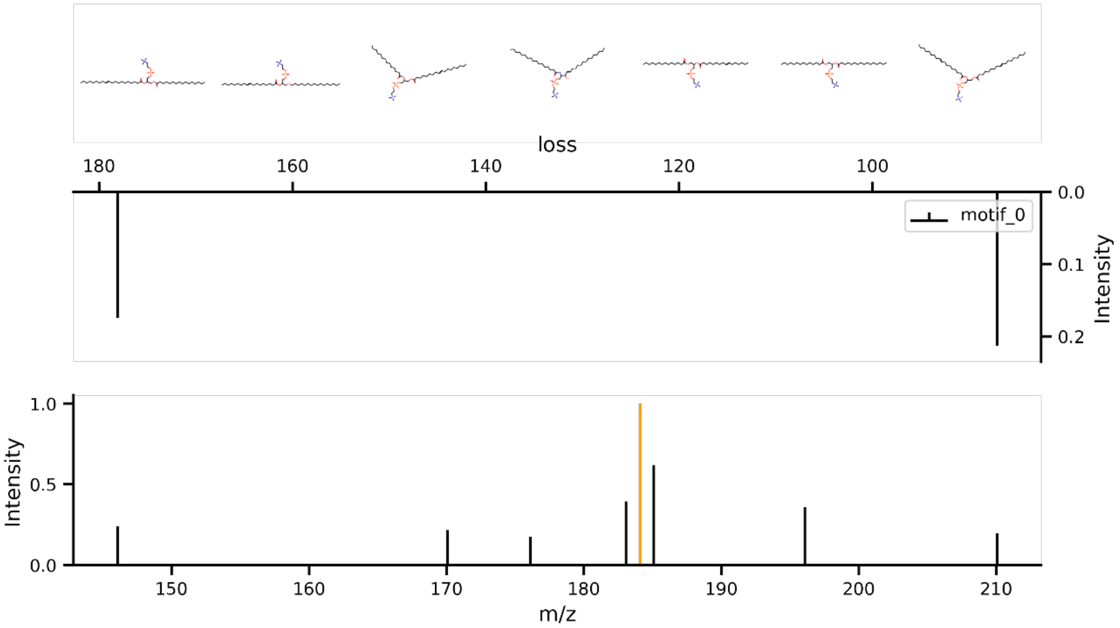

**Fig. S5.** *Example of output, top panel shows structure recommendations from MAG. The bottom panel displays the fragments associated with this Mass2Motif (black) as a standard spectrum. The orange-highlighted fragment is consistently observed across all recommended structures and represents the characteristic fragmentation of the phosphate group. Mirrored above the fragments are the neutral losses related to the Mass2Motif, arranged from right to left in increasing mass difference from the precursor ion.*

#### 1.3 Substructure Overlap Score Definition

The first step in benchmarking MAG was to define a metric that quantifies how well a predicted substructure aligns with a manual annotated structure. We defined a substructure representation for each Mass2Motif using the Mass2Motif fingerprint (see Method section) and compared it to the fingerprint derived from manual annotations. The manual annotations were derived from three MotifSets in MotifDB: Urine-derived Mass2Motifs[2], GNPS[3] library Mass2Motifs, and Massbank[3] library-derived Mass2Motifs. Since the Mass2Motif annotations in MotifDB are given as free-text descriptions, each description was manually converted to a SMILES representation. For example, the annotation “C7H7 and C5H5 fragments – indicative of methylbenzene substructure (aromatic)” was translated to the SMILES string “CC1=CC=CC=C1”. SMILES were retrieved from PubChem using the compound names, and all mappings are documented in our GitHub repository that can be found here: https://github.com/vdhooftcompmet/MS2LDA/tree/main/notebooks/Paper_results.

To compare manual and automatic annotations, we introduce the Substructure Overlap Score (SOS, formula 1). This metric quantifies how much of the smaller fingerprint is present in the larger one (with smaller, we refer to the number of 1s in a fingerprint and not the number of bits).

$$\text{SOS}=\frac{(fp_{smaller} \cap fp_{bigger})}{\sum_{b=0}^{B} fp_{smaller_b}} \qquad (1)$$

The decision to use always the smaller fingerprint instead of always using the manually annotated fingerprint as substructure was made since manual annotations in MotifDB do not consistently refer to well-defined substructures. For instance, annotations range from general substructures like "phthalate substructure" to full compound-based descriptions such as “(5-Hydroxy-2,2-dimethyl-4-oxo-3,4-dihydro-2H-chromen-7-yl)oxy substructure.”

In the main manuscript, we report SOS values based on MACCS fingerprints, as they align more closely with the information derived from mass spectrometry data. Unlike Daylight (RDKit) fingerprints, which are sensitive to subtle differences in molecular topology, MACCS fingerprints are key-based and remain consistent across variations that would not produce different fragmentation patterns. However, MACCS fingerprints may also overestimate similarity by grouping together distinct structures that would yield different fragmentation spectra. Despite this, MACCS provides a pragmatic and arguably more realistic estimate of annotation accuracy than RDKit fingerprints alone. As shown in Fig. S6–S8, the differences between RDKit- and MACCS-based SOS scores are relatively minor.

#### 1.4 Extended Benchmarks

This section presents benchmarks for three selected MotifSets. In addition to the results obtained using MACCS fingerprints, we also include results based on RDKit (Daylight) fingerprints. To validate the use of the SOS (Substructure Overlap Score) as a performance metric for MAG, we compared the MotifSet predictions with randomly selected compounds from the Spec2Vec reference database and processed them in the same way as for the Mass2Motif fingerprints.

Overall, the SOS score benchmarks demonstrate the improved performance of MAG compared to a random baseline (Fig. S6-S8). While randomly selected compounds from the reference database may occasionally show high structural overlap, primarily due to the presence of very common substructures, such matches are less abundant. In contrast, MAG consistently identifies structurally coherent matches, as reflected by significantly higher SOS scores in terms of distribution, mean, and average values. Through the comparison of randomly selected molecules as a baseline and the MAG annotation, we validated that MAG can suggest plausible structural matches for newly discovered Mass2Motifs with shared chemistry in mass spectral libraries.

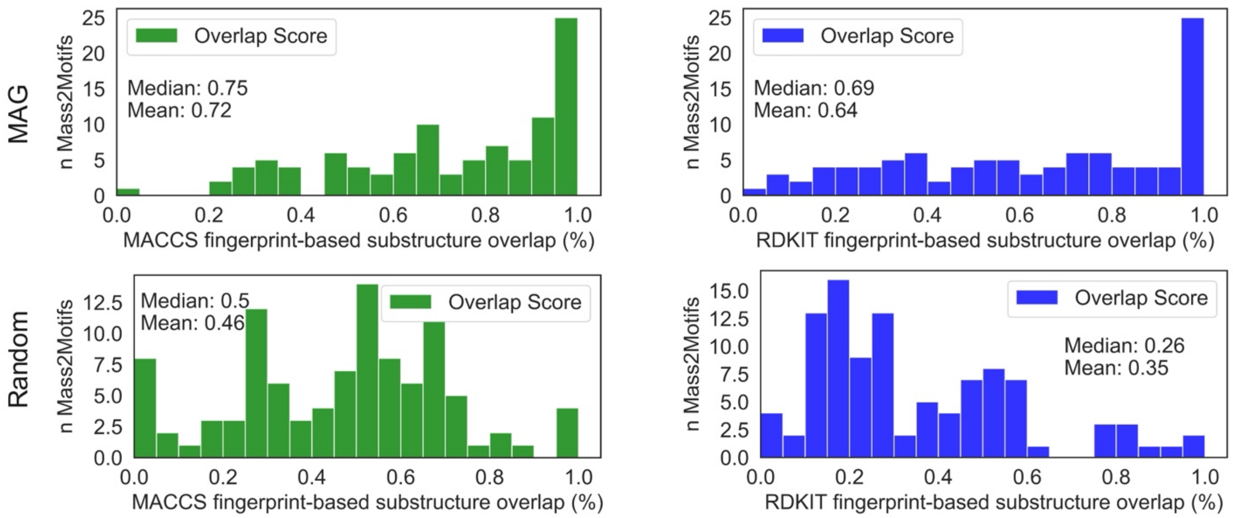

**Fig. S6.** *Substructure overlap distribution of the scores with MACCS fingerprints (green) and RDKit fingerprints (blue) for MAG compared to randomly selected molecules for the urine-derived MotifSet. This represents how MAG outperforms the baseline of randomly selected molecules by 0.25.*

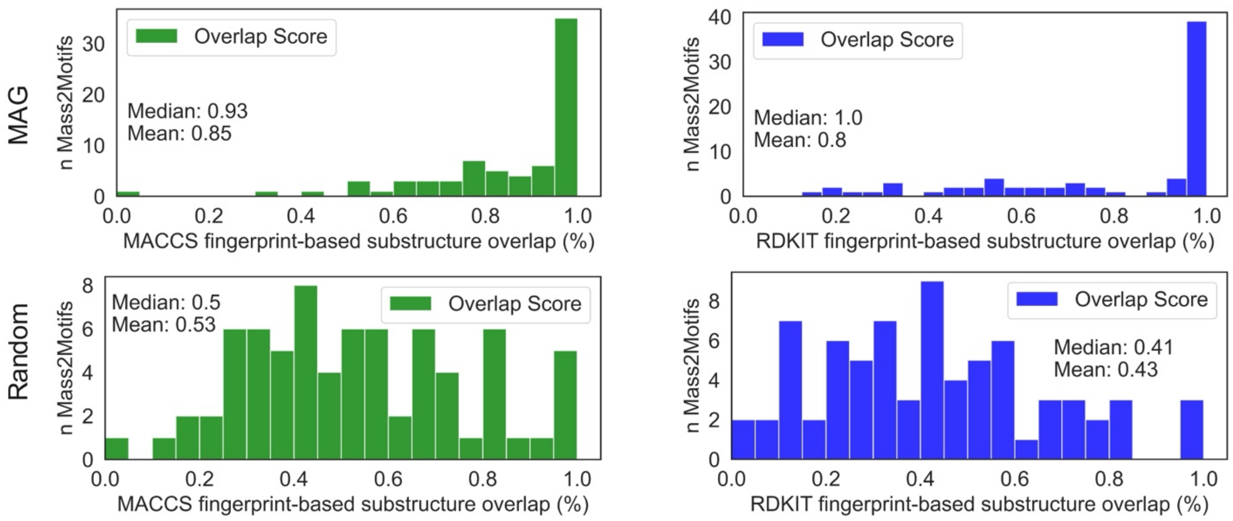

**Fig. S7***. Substructure overlap distribution of the scores with MACCS fingerprints (green) and RDKit fingerprints (blue) for MAG compared to randomly selected molecules for the GNPS MotifSet. This represents how MAG outperforms randomly selected molecules in terms of overlapping scores by 0.3.*

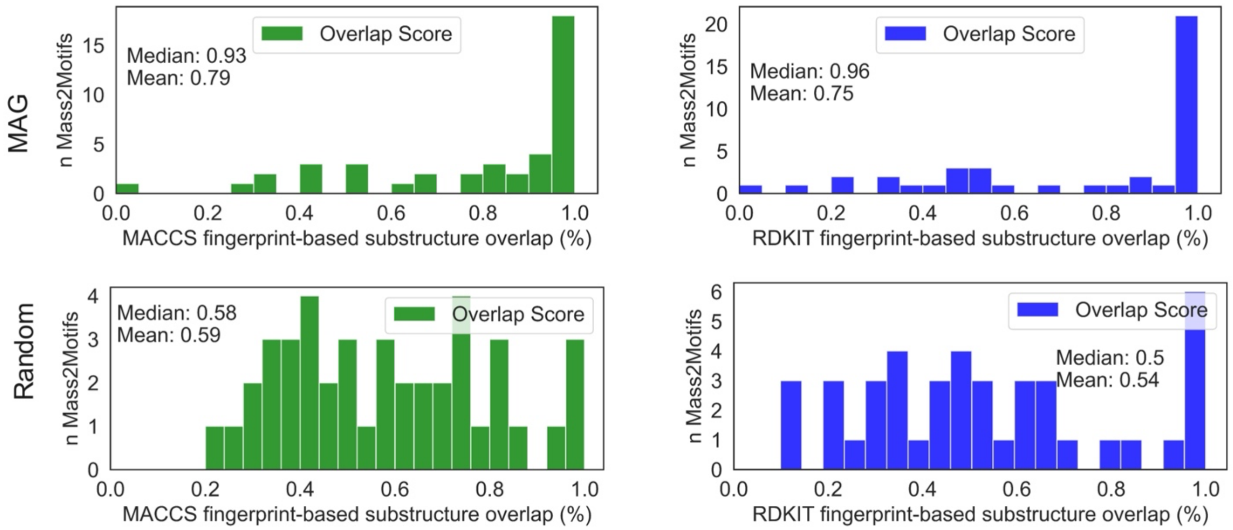

**Fig. S8.** *Substructure overlap distribution of the scores with MACCS fingerprints (green) and RDKit fingerprints (blue) for MAG compared to randomly selected molecules for the MassBank MotifSet. This represents how MAG outperforms randomly selected molecules in terms of overlapping scores by 0.2.*

### 2. Case Study Pesticides

#### 2.1 Additional Mass2Motifs related to Pesticides

Beyond the Mass2Motifs highlighted in the case study on pesticide section in the main text, other pesticide-related Mass2Motifs could also be successfully identified.
For instance, the fragment at *m/z* 72.044 serves as a distinctive marker for dimethylated carbamates, which is a common substructure in many pesticides. Although this class is often defined by a single fragment, its presence provides a strong indication for flagging relevant compounds (Fig. S9).

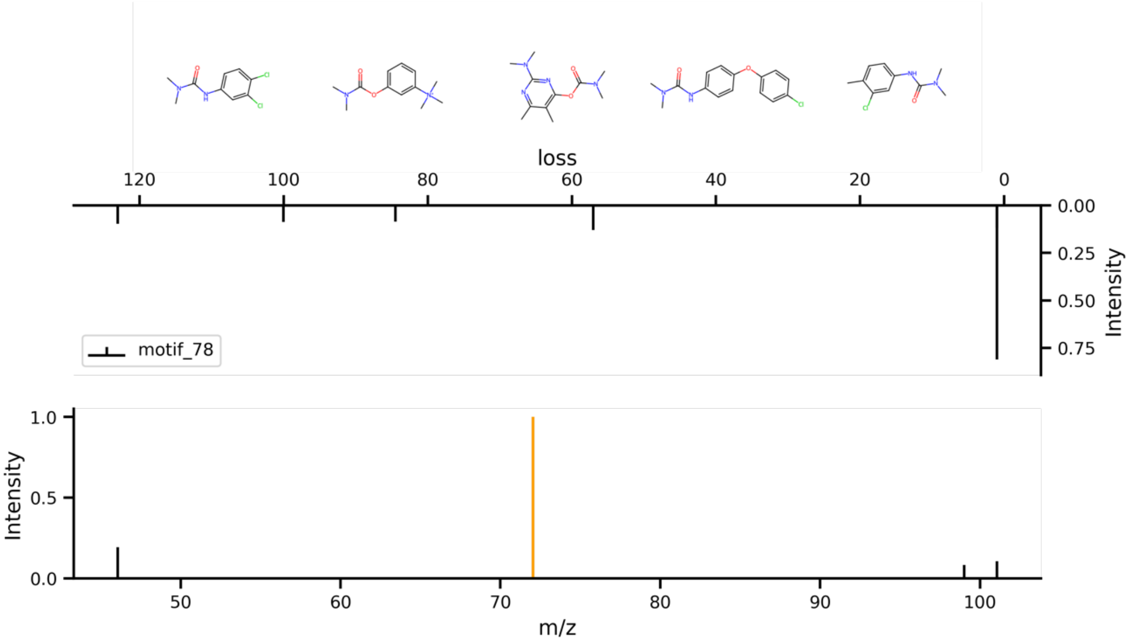

**Fig. S9.** *Mass2Motif 78 is associated with dimethyl-carbamate substructures. The top panel shows structure recommendations from MAG, all featuring dimethyl-carbamate moieties. The bottom panel displays the fragments associated with this Mass2Motif as a standard MS/MS spectrum. The orange-highlighted fragment at m/z 72.04 is consistently observed across all recommended structures and represents the characteristic fragmentation of the dimethyl-carbamate group. Mirrored above the fragments are the corresponding neutral losses, arranged from right to left in increasing mass difference from the precursor ion.*

Another prevalent pesticide group are the triazole pesticides. A specific triazole pesticide class is characterized by a triazole ring and a chlorinated benzene, which are captured by Mass2Motif 98 through the fragments at *m/z* 70.040 (triazole) and *m/z* 125.015 (chlorinated benzene). We note that, even though these substructures are not directly connected, MS2LDA 2.0 is able to recognize the co-occurrence pattern, demonstrating its capacity to associate distinct but recurring fragments (Fig. S10).

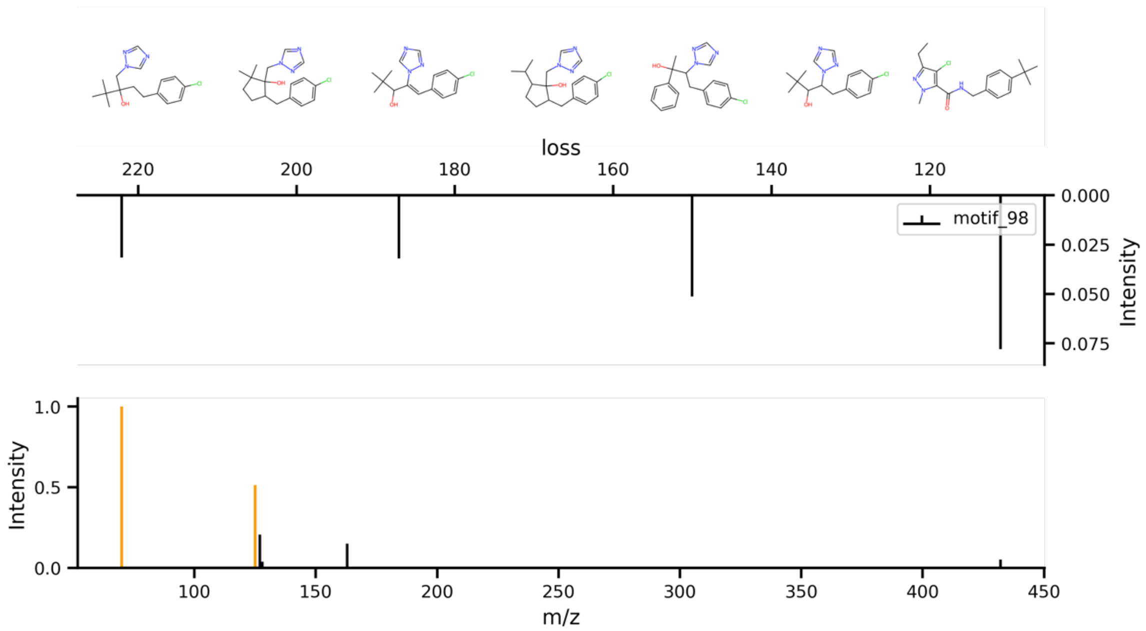

**Fig. S10.** *Mass2Motif 98 represents a common pesticide related pattern of two substructures. At the top the structure recommendations by MAG are shown. Below are the intensities of the losses related to the Mass2Motif and it is read from the precursor from right to left. At the bottom the fragments related to the Mass2Motif are shown as a normal spectrum plot. The orange highlighted fragments of 70.04 and 125.02 is present in all the spectra from structure recommendations from the top.*

#### 2.2 LDA probabilities for reported pesticides classes and their Mass2Motifs

For every spectrum in the input file for the MS2LDA 2.0 modelling, a probability score is calculated, which explains how much a spectrum is associated with a Mass2Motif. The scores are between 0 and 1, where 1 means a spectrum is only strongly associated with a Mass2Motif and 0 shows no association. In Table S1, the probability scores between three different Mass2Motifs representing organophosphates, benzoylureas, and sulfonylureas and their related spiked pesticides are shown.

The association between the compounds and Mass2Motifs highlighted by the probabilities reaches from 0.05 to 1.00 and was discussed in the case study on pesticides in the main text.

**Table S1.** *Probabilities between Mass2Motif and spiked pesticides for three different compound classes.*

| Compound Name | Mass2Motif | θ Probability |
|---|---|---|
| **Organophosphates** | | |
| Omethoate | 6 | 1.00 |
| Dimethoate | 6 | 0.99 |
| Mevinphos | 6 | 0.81 |
| Cythioate | 6 | 0.74 |
| Oxydemeton-methyl | 6 | 0.40 |
| Tolclofos-methyl | 6 | 0.32 |
| Pirimiphos-methyl | 6 | 0.02 |
| Malathion | 6 | 0.05 |
| **Benzoylureas** | | |
| Diflubenzuron | 90 | 0.99 |
| Flufenoxuron | 90 | 0.46 |
| **Sulfonylureas** | | |
| Nicosulfuron | 236 | 1.00 |
| Foramsulfuron | 236 | 0.98 |
| Mesosulfuron-methyl | 236 | 0.97 |
| Flupyrsulfuron-methyl | 236 | 0.62 |
| Rimsulfuron | 236 | 0.57 |
| Amidosulfuron | 236 | 0.10 |

#### 2.3 Spiked compounds

As part of the sample preparation (see Methods), a blank tomato sample was spiked with 211 pesticides. In Table S2, a complete list of all the pesticides can be found.

**Table S2**. *List of all 211 spiked pesticides.*

| Compound Name | Compound Name | Compound Name |
|---|---|---|
| 2,4-D | Abamectin | Acephate |
| Acetamiprid | Aldicarb | Ametoctradin |
| Amidosulfuron | Asulam | Azamethiphos |
| Azoxystrobin | Bendiocarb | Bentazone |
| Benthiavalicarb-isopropyl | Benzovindiflupyr | Bifenazate |
| Bifenthrin | Bitertanol | Bixafen |
| Boscalid | Brodifacoum | Bromadiolone |
| Bromoxynil | Bromuconazole | Bupirimate |
| Buprofezin | Carbaryl | Carbendazim |
| Carbetamide | Carbofuran | Carfentrazone-ethyl |
| Chlorantraniliprole | Chlorbromuron | Chloridazon |
| Chlorpyrifos | Clodinafop-propargyl | Clofentezine |
| Clomazone | Clothianidin | Cyantraniliprole |
| Cyazofamid | Cybutryne | Cyflufenamid |
| Cyflumetofen | Cymoxanil | Cyproconazole |
| Cyprodinil | Cythioate | Cyfluthrin |
| Cyhalothrin | Cypermethrin | Deltamethrin |
| Desmedipham | Dichlofluanid | Difenoconazole |
| Diflubenzuron | Dimethenamid | Dimethoate |
| Dimethomorph | Dinoterb | Diuron |
| DNOC | Dodemorph | Dodine |
| Emamectin B1a | Epoxiconazole | Ethirimol |
| Ethoprophos | Etoxazole | Famoxadone |
| Fenamidone | Fenamiphos | Fenhexamid |
| Fenoxaprop-p-ethyl | Fenoxycarb | Fenpropidin |
| Fenpropimorph | Fenpyrazamine | Fipronil |
| Flonicamid | Florasulam | Fluazifop |
| Fluazinam | Flucycloxuron | Fludioxonil |
| Flufenacet | Flufenoxuron | Flumioxazin |
| Fluopicolide | Fluopyram | Fluoxastrobin |
| Flupyradifurone | Flupyrsulfuron-methyl | Fluroxypyr |
| Fluroxypyr-meptylester | Flutolanil | Fluxapyroxad |
| Foramsulfuron | Fosthiazate | Haloxyfop |
| Haloxyfop-P-methyl | Imazalil | Imazamox |
| Imidacloprid | Indoxacarb | Iodosulfuron-methyl |
| Ioxynil | Iprovalicarb | Isoproturon |
| Isopyrazam | Isoxaben | Isoxadifen-ethyl |
| Isoxaflutole | Kresoxim-methyl | Lenacil |
| Linuron | Malathion | Mandipropamid |
| MCPA | MCPP | Mepanipyrim |
| Mesosulfuron-methyl | Mesotrione | Metalaxyl |
| Metamitron | Metazachlor | Metconazole |
| Methabenzthiazuron | Methamidophos | Methiocarb |
| Methomyl | Methoxyfenozide | Metobromuron |
| Metolachlor | Metoxuron | Metrafenone |

| | | |
|---|---|---|
| Metribuzin | Metsulfuron-methyl | Mevinphos |
| Myclobutanil | Napropamide | Nicosulfuron |
| Omethoate | Oxadixyl | Oxamyl |
| Oxathiapiprolin | Oxydemeton-methyl | Paclobutrazol |
| Penconazole | Pencycuron | Pendimethalin |
| Penflufen | Penthiopyrad | Phenmedipham |
| Picoxystrobin | Pinoxaden | Pirimicarb |
| Pirimiphos-methyl | Prochloraz | Profenofos |
| Propamocarb | Propiconazole | Propisochlor |
| Propyzamide | Prosulfocarb | Prosulfuron |
| Prothioconazole, desthio | Pymetrozine | Pyraclostrobin |
| Pyridaben | Pyridate | Pyrimethanil |
| Pyriproxyfen | Pyroquilon | Pyroxsulam |
| Quinmerac | Quinoclamine | Quinoxyfen |
| Quizalofop-ethyl | Rimsulfuron | Silthiofam |
| Simazine | Spinosyn A | Spinosyn D |
| Spirodiclofen | Spiromesifen | Spirotetramat |
| Spiroxamine | Sulcotrione | Tebuconazole |
| Tebufenpyrad | Tembotrione | Tepraloxydim |
| Terbuthylazine | Terbutryn | Tetraconazole |
| Thiabendazole | Thiacloprid | Thiamethoxam |
| Thiencarbazone-methyl | Thifensulfuron-methyl | Thiophanate-methyl |
| Tolclofos-methyl | Tolylfluanid | Topramezone |
| Triazophos | Tribenuron-methyl | Triclopyr |
| Trifloxystrobin | Triflumizole | Triflumuron |
| Triflusulfuron-methyl | Triforine | Trinexapac-ethyl |
| Tritosulfuron | Zoxamide | |

### 3. Case Study Natural Products

MS2LDA 2.0 identified common substructures from the molecular families described by Khatib et. al.[4]. We followed a strategy to dereplicate the Mass2Motifs present for each family: 1) we mapped the Mass2Motifs on the molecular families previously reported, 2) we explored the annotation of these motifs by MAG, 3) if a hit was retrieved, we queried the top three peaks present per Mass2Motif against MotifDB to explore its potential manual annotation using MassQL[5], and 4) we explore the results from the Mass2Motif-matching functionality against three MotifSets: GNPS, MassBank, and LDB MotifDB POS.

#### 3.1 Hericenone family findings

For the hericenone family we identified three main Mass2Motifs in its molecular family: Mass2Motif 163, Mass2Motif 186, and Mass2Motif 90, all nodes represented by at least one of them (Fig. S11). From the family reported by Khalib et. al.[4] the node with a precursor mass of 595.39 m/z, manually annotated as hericenone H, is represented by Mass2Motif 163 (Figure S13) and 90 (Figure S15). Hericenone G, with a precursor mass of 599.43 m/z, shared the same motifs. Finally, the manually annotated node with a precursor mass of 571.3986 *m/z*, is represented by Mass2Motif 186 (Fig. S13) and 90. All manually annotated nodes shared Mass2Motif 90, which was annotated as a 2-(2-methylbut-2-enamido)acetic moiety by its first recommendation as a 4-fluoro-2-(4-hydroxyoxan-3-yl)-5-methyl-6-[(4-pyrazol-1-ylphenyl)methyl]-3H-isoindol-1-one by the second structural recommendation (Fig. S11, b). By inspecting these results, we can verify that the first recommendation 2-(2-methylbut-2-enamido) acetic, aligns more with the carbon chain side shown in all the hericenone molecules. On the other hand, both Mass2Motif 163 and Mass2Motifs 186 represent the same substructure, a benzodihydropyran, recommended by MAG, both substructures aligning well with the manually annotated molecules (Fig. S12 and S13). These two motifs shared most of the peaks, which is a likely reason they retrieved the same structure.

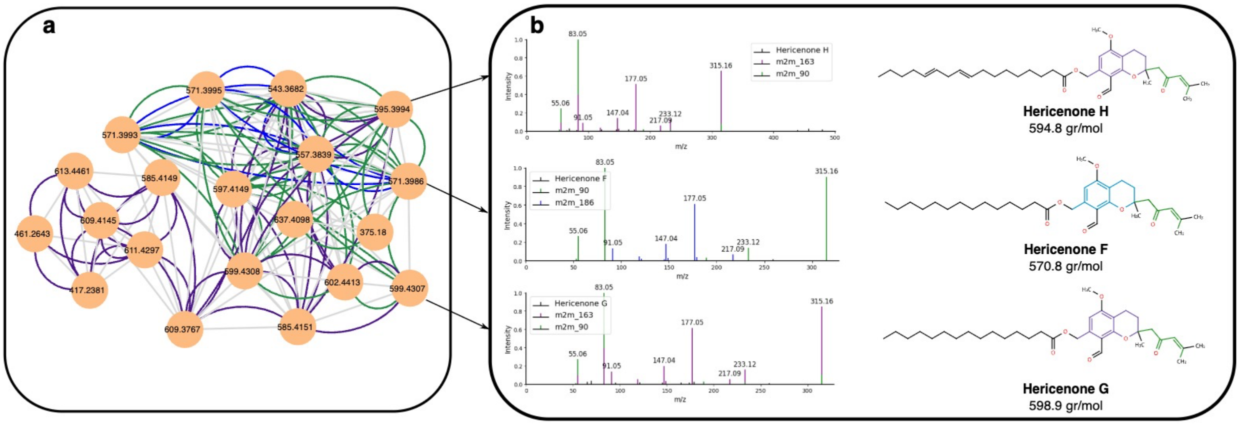

**Fig. S11.** *MS2LDA 2.0 results from the fungal natural product dataset for the hericenone family. a) Molecular family from hericenones originated reported by Kathib et al [4]. b) Spectra of the molecules manually annotated as hericenone H, F and G respectively with the Mass2Motifs related fragments colored on top. Structures of the complete erinacerin molecules with the inferred Mass2Motifs by MAG colored.*

All motifs were queried against all MotifDB entries. For Mass2Motif 163, we did not obtain any hits, even though their MAG recommendation is very similar to the Mass2Motif 186 recommendation (Fig. S12 and S13), i.e. both share a two coumarin-like related structures. However, the top three peaks (by intensity) differ between the two Mass2Motifs, which could influence the retrieved hits obtained through MassQL. For Mass2Motif 186, MotifDB correctly retrieved terpenoid-like Mass2Motifs for its annotation (Table S3).

Finally, for Mass2Motif 90, the MassQL results are not conclusive, since it retrieves a wide variety of structures such as sterone related, phenols, deterpenes, etc. (Table S4). In this case, the discrepancy may be due to the presence of only one top peak (in red), which is the only peak present in the structure retrieved by MAG (Fig. S14).

The above findings demonstrate how the increased annotation capabilities through various routes in MS2LDA provide the user with additional sources for structure annotation guidance that return useful and relevant structural information on the experimentally discovered Mass2Motifs.

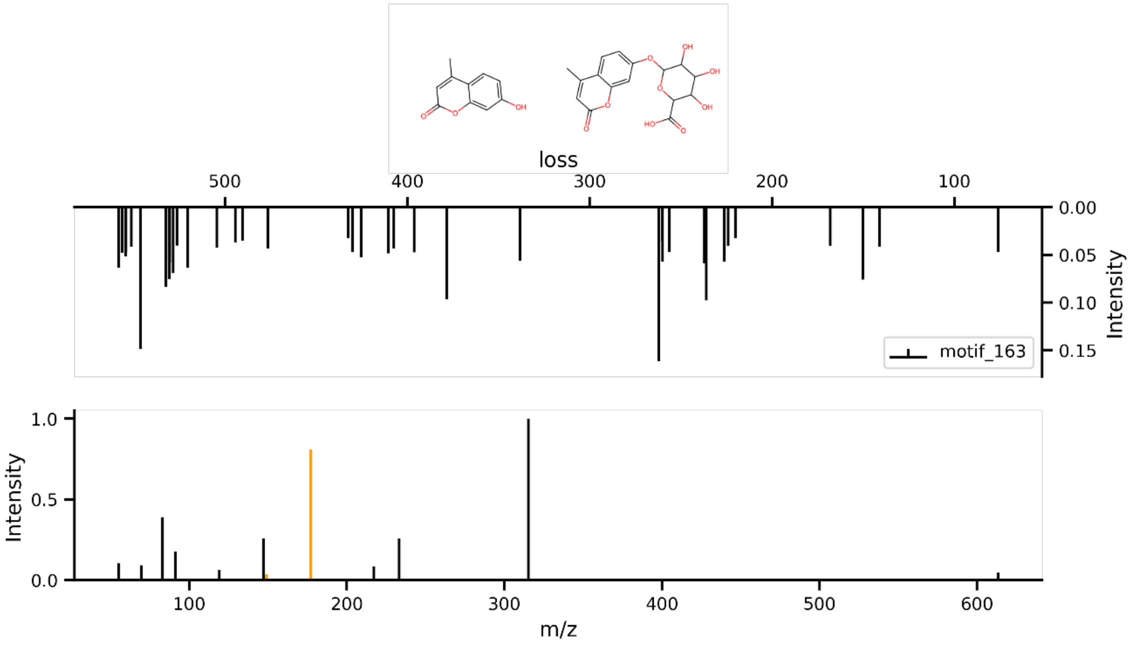

**Fig. S12**. *Mass2Motif 163 is associated with benzodihydropyran substructures. The top panel shows structure recommendations from MAG, all featuring benzodihydropyran moieties. The bottom panel displays the fragments associated with this Mass2Motif as a standard MS/MS spectrum. The orange-highlighted fragment at m/z 177.05 is consistently observed across all recommended structures and represents the characteristic fragmentation of the benzodihydropyran substructure. Mirrored above the fragments are the corresponding neutral losses, arranged from right to left in increasing mass difference from the precursor ion.*

**Table S3.** *Results from the query Mass2Motif 186 against MotifDB.*

| **motifset** | **motif_id** | **short_annotation** | **annotation** |
|---|---|---|---|
| **LDB MotifDB POS** | motif_92 | No short annotation available | 15 spectra, 6 molecules, 3 classes: 66.7% Paraconic acids, 16.7% Acids, 16.7% Pulvinic Acid Derivatives |
| **LDB MotifDB POS** | motif_23 | No short annotation available | 6 spectra, 4 molecules, 3 classes: 50.0% Chromanes and Chromones, 25.0% Terpenoids : Triterpenes, 25.0% Depsides (Didepsides) |
| **Euphorbia Plant Mass2Motifs** | motif_333 | Diterp 309 307 291 283 279 | Diterp 309 307 291 283 279 |
| **LDB_NEG_MotifDB_02** | motif_4 | No short annotation available | No annotation available |
| **Euphorbia Plant Mass2Motifs** | motif_3 | Loss of hexadienoic acid and its fragment / difference of 18 corresponds to the loss of-NH3 adduct | Loss of hexadienoic acid |
| **Euphorbia Plant Mass2Motifs** | motif_92 | Diterp 297 279 269 | Diterp 297 279 269 |
| **Euphorbia Plant Mass2Motifs** | motif_14 | Diterp 611 333 277 275 259 | Diterp 611 333 277 275 259 |

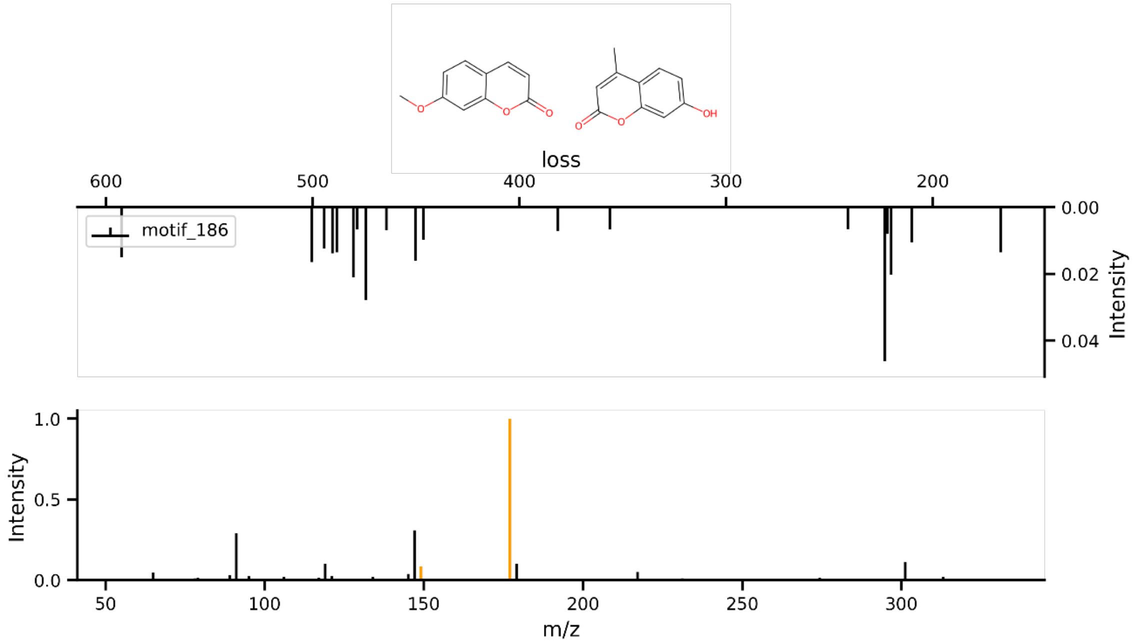

**Fig. S13**. *Mass2Motif 186 is associated with benzodihydropyran substructures. The top panel shows structure recommendations from MAG, all featuring benzodihydropyran moieties. The bottom panel displays the fragments associated with this Mass2Motif as a standard MS/MS spectrum. The orange-highlighted fragments at m/z 177.05 and 149.06 are consistently observed across all recommended structures and represents the characteristic fragmentation of the benzodihydropyran substructure. Mirrored above the fragments are the corresponding neutral losses, arranged from right to left in increasing mass difference from the precursor ion.*

**Table S4.** *Results from the query Mass2Motif 90 against MotifDB.*

| motifset | motif_id | short_annotation | annotation |
|---|---|---|---|
| **Urine derived Mass2Motifs** | motif_29 | Major loss is related to loss of sulphate group from phenolic (aromatic) structure (SO3 loss) | Major loss is related to loss of sulphate group from phenolic (aromatic) structure (SO3 loss) |
| **GNPS library derived Mass2Motifs** | motif_115 | Sterone steroid related | Sterone steroid related Mass2Motif |
| **Euphorbia Plant Mass2Motifs** | motif_333 | Diterp 309 307 291 283 279 | Diterp 309 307 291 283 279 |
| **Urine derived Mass2Motifs** | motif_22 | Mixed motif of Quinidine (urine 97) and different cores in other urines | Mixed motif of Quinidine (urine 97) and different cores in other urines |
| **LDB_NEG_MotifDB_02** | motif_37 | No short annotation available | No annotation available |
| **Urine derived Mass2Motifs** | motif_68 | Alkyl aromatic substructure â€“ indicative for aromatic ring with 2-carbon alkyl chain attached i.e. phenylethene fragment | Alkyl aromatic substructure â€“ indicative for aromatic ring with 2-carbon alkyl chain attached i.e. phenylethene fragment |
| **Urine derived Mass2Motifs** | motif_106 | unsat diC-FA | Unsaturated di-carboxilic fatty acids related Mass2Motif |
| **Euphorbia Plant Mass2Motifs** | motif_14 | Diterp 611 333 277 275 259 | Diterp 611 333 277 275 259 |

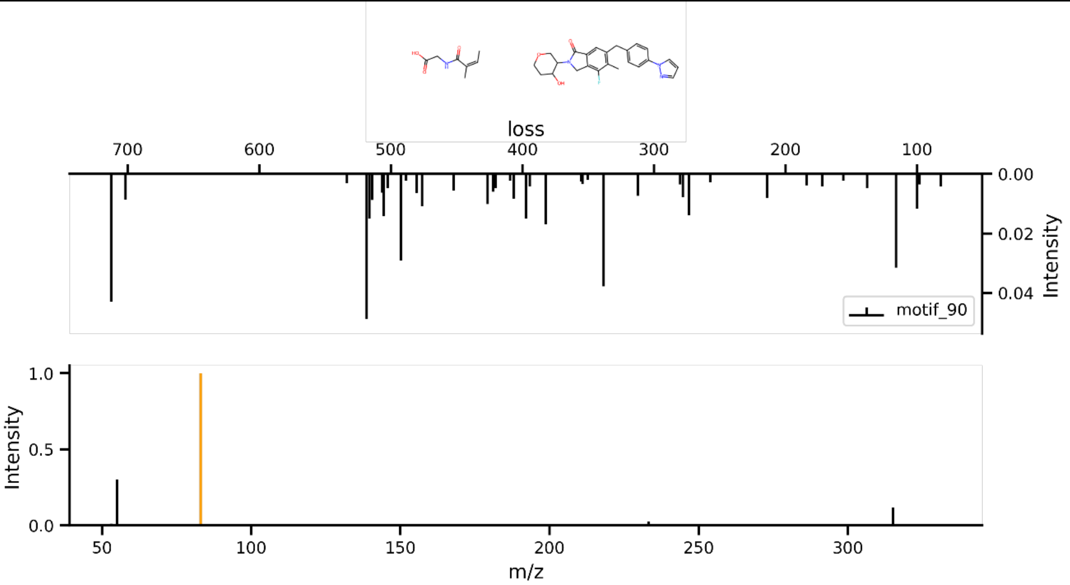

**Fig. S14**. *Mass2Motif 90 is associated with 2-(2-methylbut-2-enamido)acetic and 4-fluoro-2-(4-hydroxyoxan-3-yl)-5-methyl-6-[(4-pyrazol-1-ylphenyl)methyl]-3H-isoindol-1-one structures. The top panel shows structure recommendations from MAG. The bottom panel displays the fragments associated with this Mass2Motif as a standard MS/MS spectrum. The orange-highlighted fragment at m/z 83.05 is consistently observed across both recommended structures. Mirrored above the fragments are the corresponding neutral losses, arranged from right to left in increasing mass difference from the precursor ion.*

#### 3.2 Ergostane family findings

We identified the top abundant Mass2Motifs present in the reported ergostane molecular family: Mass2Motif 173, Mass2Motif 0, Mass2Motif 21, and Mass2Motif 193. All nodes were represented by at least one of these motifs (Fig. S15). Mass2Motif 173 is present in two out of three manually annotated nodes, one corresponding to citreoanthrasteroide B, with a precursor mass of 393.315 *m/z* and ergosta-5,7,22,24(28)-tetraen-3beta-ol with a precursor mass of 395.33 *m/z,* its MAG recommendation corresponds highly with the original structures, with an steroid scaffold and a carbon side chain, as [17-(5,6-dimethylhept-3-en-2-yl)-4,4,10,13-tetramethyl-1,2,3,9,11,12,14,15,16,17-decahydrocyclopenta[a]phenanthren-3-yl] acetate.

Ergosta-5,7,22,24(28)-tetraen-3beta-ol is also represented by Mass2Motif 21 (Fig. S17), which MAG recommendation points out a 2,15-dihydroxy-7-methyl-6-oxabicyclo[11.3.0]hexadeca-3,11-dien-5-one, which is not closely similar to the original structure proposed but they share the five-fused ring with the steroid backbone. We did not retrieve any results from MassQL-quering for this Mass2Motif.

For the last manually annotated compound, ergost-3,5,7,9(11),22-pentaen, it also contains Mass2Motif 21, and Mass2Motifs 0 and 193. For Mass2Motif 0 (Fig. S18), the MAG recommendation closely resembles the chain side carbon structure of this family reported previously (Fig. S18, b), the MassQL results retrieved a diverse class of motif-related structures such as depsidones, quinones and paraconic acids (Table S5). Mass2Motif 193 (Fig. 19) MAG recommendation features a sterol core structure without any side chain, a 5',7,9,13-tetramethylspiro[5-oxapentacyclo[10.8.0.02,9.04,8.013,18]icos-18-ene-6,2'-oxane]-16-ol. There were no retrieved Mass2Motifs from MassQL for this Mass2Motif.

These results show how MAG is a powerful assistant for Mass2Motif annotation, different Mass2Motifs may represent different part of a compound such as Mass2Motif 21 and Mass2Motif 0 represent ergost-3,5,7,9(11),22-pentaen. Thus, analyzing the Mass2Motifs related to some compounds can help to reveal their structure.

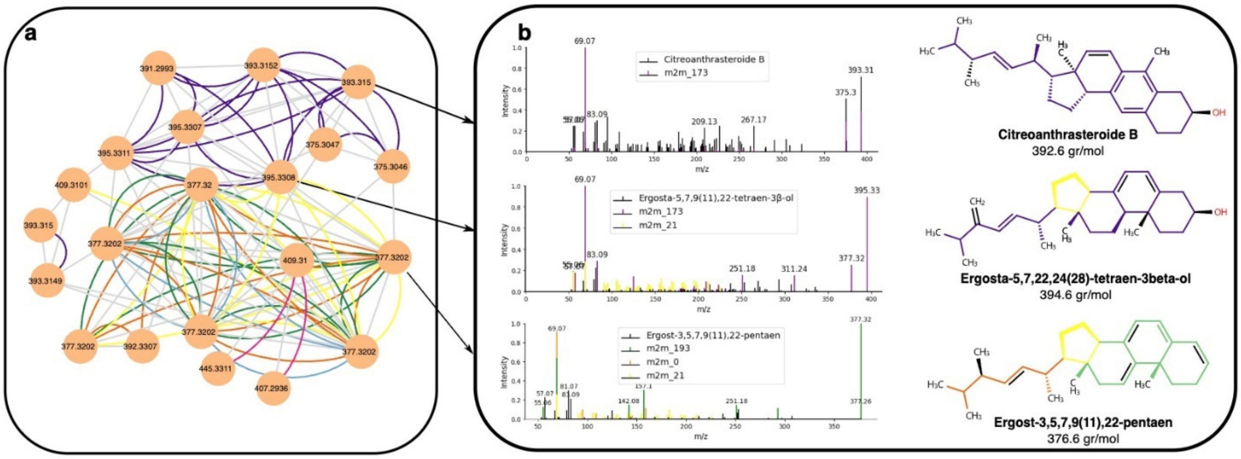

**Fig. S15.** *MS2LDA 2.0 results from the fungal natural product dataset for the ergostane family. a) Molecular family from ergostane originated reported by Khatib et al.[4], b) Spectra of the molecules manually annotated as citreoanthrasteroide B, Ergosta-5,7,22,24(28)-tetraen-3beta-ol and ergost-3,5,7,9(11),22-pentaen with their Mass2Motifs related fragments colored on top. Structures of the complete erinacerin molecules with the inferred Mass2Motifs by MAG colored.*

**Table S5.** *Results from the query Mass2Motif 173 against MotifDB.*

| motifset | motif_id | short_annotation | annotation |
|---|---|---|---|
| **Urine derived Mass2Motifs** | motif_236 | Mixed motif of modified Diltiazem related (urines 51 and 85) and smaller (methoxylated) phenolic substructure related Mass2Motif | Mixed motif of modified Diltiazem related (urines 51 and 85) and smaller (methoxylated) phenolic substructure related Mass2Motif |
| **Rhamnaceae Plant Mass2Motifs** | motif_3 | -related motif | Myricetin-related motif |

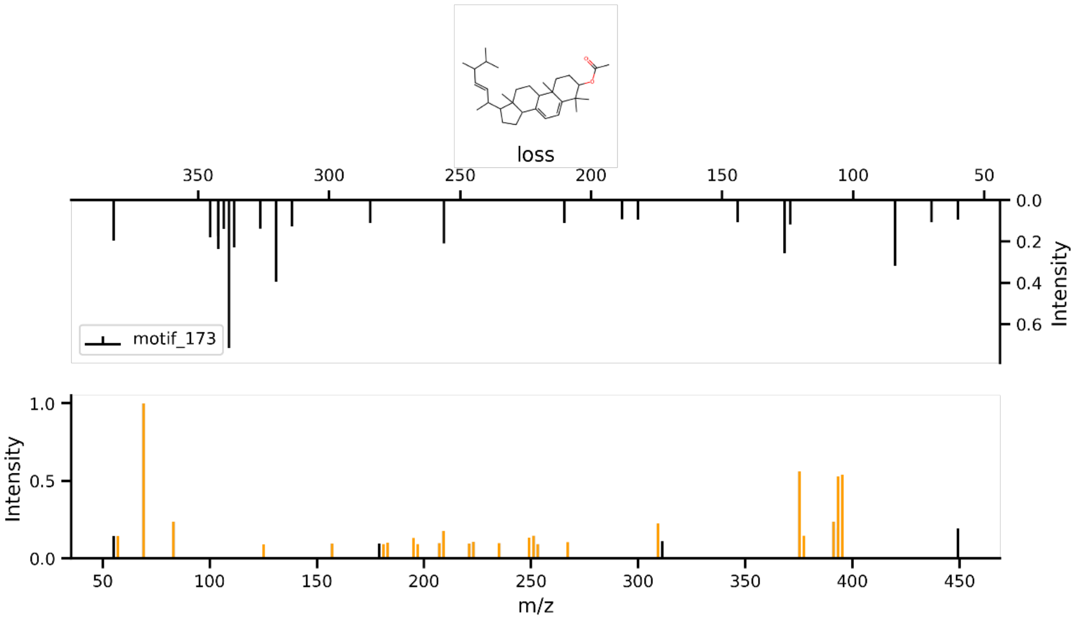

**Fig. S16**. *Mass2Motif 173 is associated with a steroid scaffold and a carbon side chain. The top panel shows structure recommendations from MAG. The bottom panel displays the fragments associated with this Mass2Motif as a standard MS/MS spectrum. The orange-multiple highlighted fragments are consistently observed across both recommended structures. Mirrored above the fragments are the corresponding neutral losses, arranged from right to left in increasing mass difference from the precursor ion.*

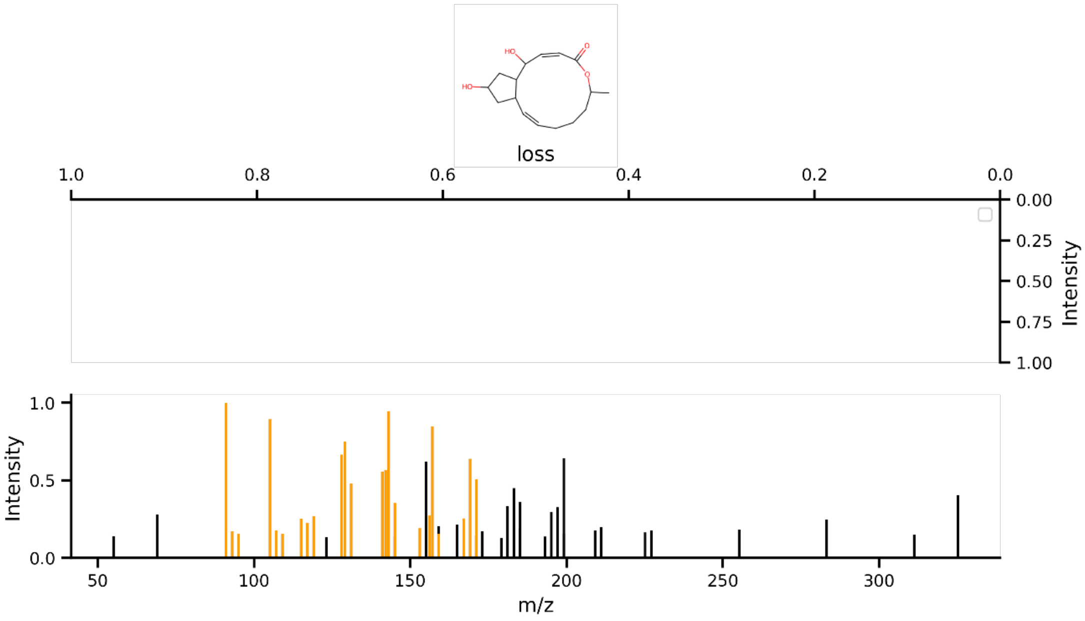

**Fig. S17**. *Mass2Motif 21 is associated with 2,15-dihydroxy-7-methyl-6-oxabicyclo[11.3.0]hexadeca-3,11-dien-5-one. The top panel shows the only structure from MAG. The bottom panel displays the fragments associated with this Mass2Motif as a standard MS/MS spectrum. The orange-multiple highlighted fragments are consistently observed in the recommended structure. Mirrored above the fragments are the corresponding neutral losses (empty), arranged from right to left in increasing mass difference from the precursor ion.*

**Table S6**. *Results from the query Mass2Motif 0 against MotifDB.*

| motifset | motif_id | short_annotation | annotation |
|---|---|---|---|
| **LDB MotifDB POS** | motif_16 | No short annotation available | 16 spectra, 9 molecules, 4 classes: 55.6% Depsidones, 22.2% Quinones, 11.1% Acids, 11.1% Paraconic acids |
| **LDB MotifDB POS** | motif_0 | No short annotation available | 4 spectra, 4 molecules, 3 classes: 50.0% Paraconic acids, 25.0% Xanthones and bis-Xanthones, 25.0% Depsidones |
| **Urine derived Mass2Motifs** | motif_291 | Check - nitrogen containing ring core structure - could be from alkaloid? | Check - nitrogen containing ring core structure - could be from alkaloid? |

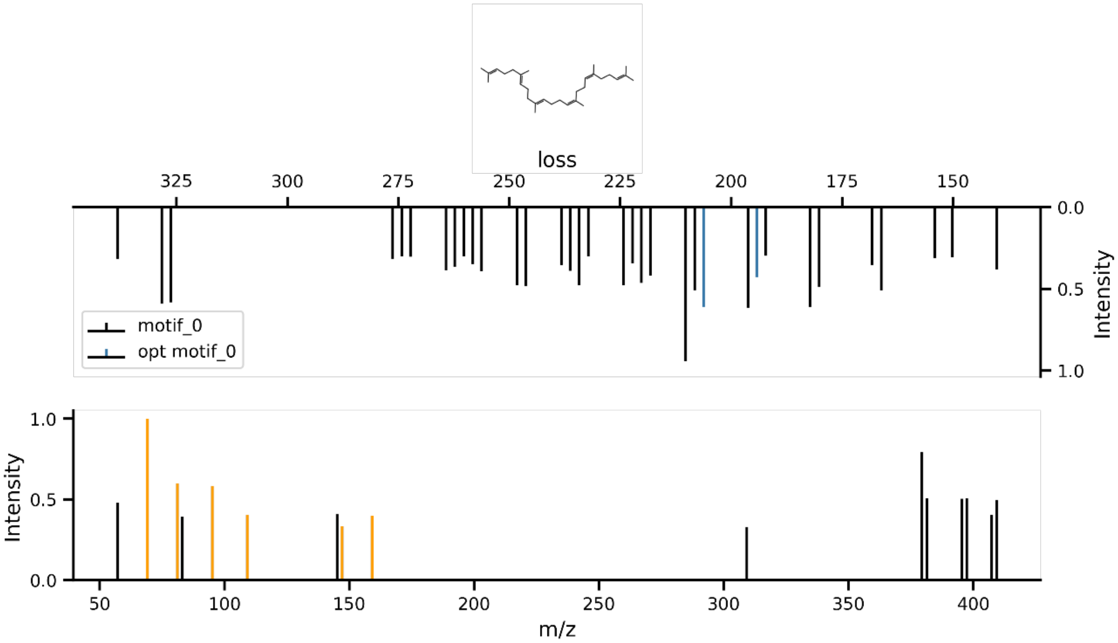

**Fig. S18**. *Mass2Motif 0 is associated with 2,15-dihydroxy-7-methyl-6-oxabicyclo[11.3.0]hexadeca-3,11-dien-5-one. The top panel shows the only structure recommended by MAG. The bottom panel displays the fragments associated with this Mass2Motif as a standard MS/MS spectrum. The orange-multiple highlighted fragments are observed in the recommended structure. Mirrored above the losses of 206.2 and 194.2 m/z are also part of the recommended structure. Losses are arranged from right to left in increasing mass difference from the precursor ion.*

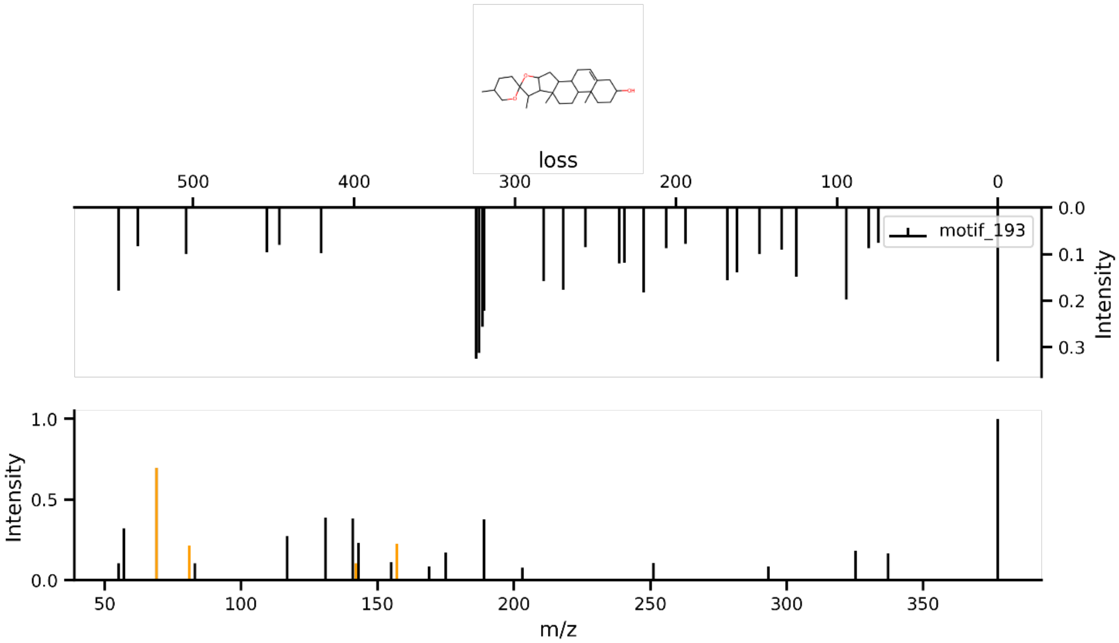

**Fig. S19**. *Mass2Motif 193 is associated with a steroid backbone. The top panel shows the only structure from MAG. The bottom panel displays the fragments associated with this Mass2Motif as a standard MS/MS spectrum. The orange-multiple highlighted fragments are consistently observed in the recommended structure. Mirrored above the fragments are the corresponding neutral losses, arranged from right to left in increasing mass difference from the precursor ion.*

#### 3.3 Erinacerin family findings

Finally, for the erinacerin family, mainly described in the main text, the manually annotated by Khatib et. al.[4], as erinacerin N with a precursor mass of 381.20 *m/z*, contains Mass2Motif 137 and Mass2Motif 22 (Fig. S21 and S22 respectively), the MAG recommendation for Mass2Motif 137 identified as 4-[4-(2-chlorophenyl)piperazin-1-yl]-4-oxobut-2-enoic acid by MAG, being the pyridine substructure also part of erinacerin N. Some of the hits for this motif from MassQL results matched part of the structure (Table S5), such as motif_59, annotated as verapamil related, this has also a six-carbon ring as part of the molecule (3S)-2-[(1R)-1-[5-acetyl-2-(hydroxymethyl)-4-oxopyridin-1-yl]-2-methylbutyl]imino-3-hydroxy-4-methylpentanoic acid. For Mass2Motif 22, the MAG recommendation, strongly supports the same substructure reported by the original paper, although no hits were retrieved by MassQL.

For the other compound manually annotated, erinacerin F, contains only Mass2Motif 108, we obtained five hits from MassQL (Table S6), most of which featured an indole-related structure, which aligns highly with the recommendation from MAG for this Mass2Motif.

For this family, both MAG and MassQL results aligned well with both compounds (erinacerin N and F) reported in the original paper. This is not always the case for every family reported in the paper. For example, the hericenone family contained redundant Mass2Motifs, where it shows the importance in hyper tuning parameters such as number of topics, alpha and beta. For this family, MassQL does not align very well with the MAG recommendation either. This could be due to differences in spectra quality. These results show how MS2LDA 2.0 framework offers multiple and valuable strategies to complement structural annotation in metabolomic studies.

**Table S7**. *Results from the query Mass2Motif 137 against MotifDB.*

| motifset | motif_id | short_annotation | annotation |
|---|---|---|---|
| **LDB MotifDB POS** | motif_92 | No short annotation available | 15 spectra, 6 molecules, 3 classes: 66.7% Paraconic acids, 16.7% Acids, 16.7% Pulvinic Acid Derivatives |
| **Euphorbia Plant Mass2Motifs** | motif_333 | Diterp 309 307 291 283 279 | Diterp 309 307 291 283 279 |
| **Euphorbia Plant Mass2Motifs** | motif_14 | Diterp 611 333 277 275 259 | Diterp 611 333 277 275 259 |
| **Urine derived Mass2Motifs** | motif_59 | Verapamil | Verapamil related (anithypertensive) |

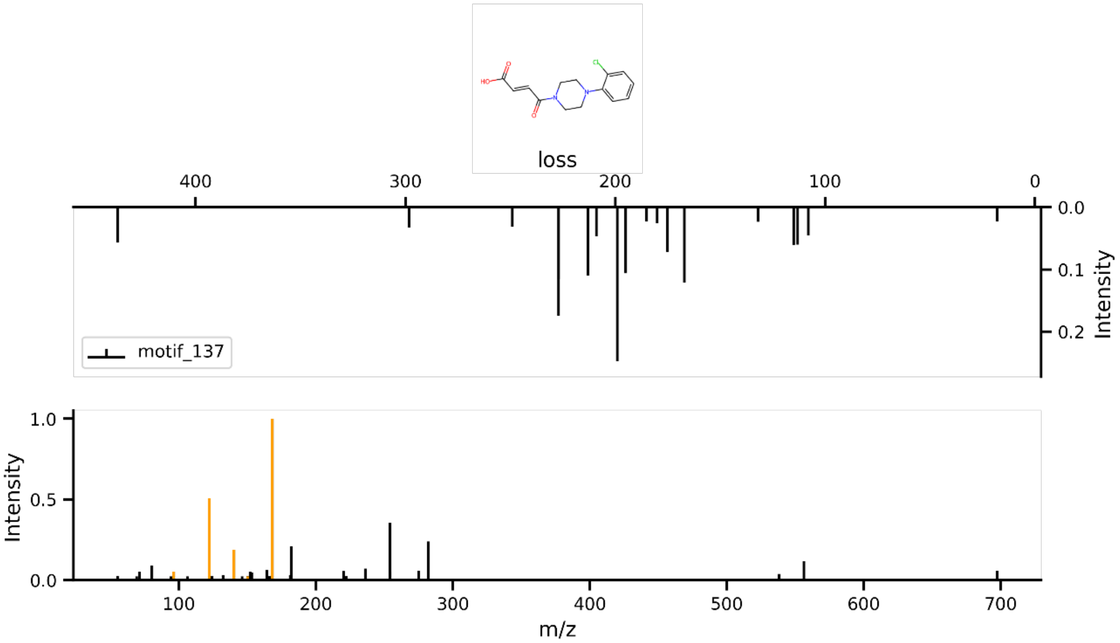

**Fig. S20**. *Mass2Motif 137 is associated with a pyridine backbone. The top panel shows the only structure from MAG. The bottom panel displays the fragments associated with this Mass2Motif as a standard MS/MS spectrum. The orange-multiple highlighted fragments are consistently observed in the recommended structure. Mirrored above the fragments are the corresponding neutral losses, arranged from right to left in increasing mass difference from the precursor ion.*

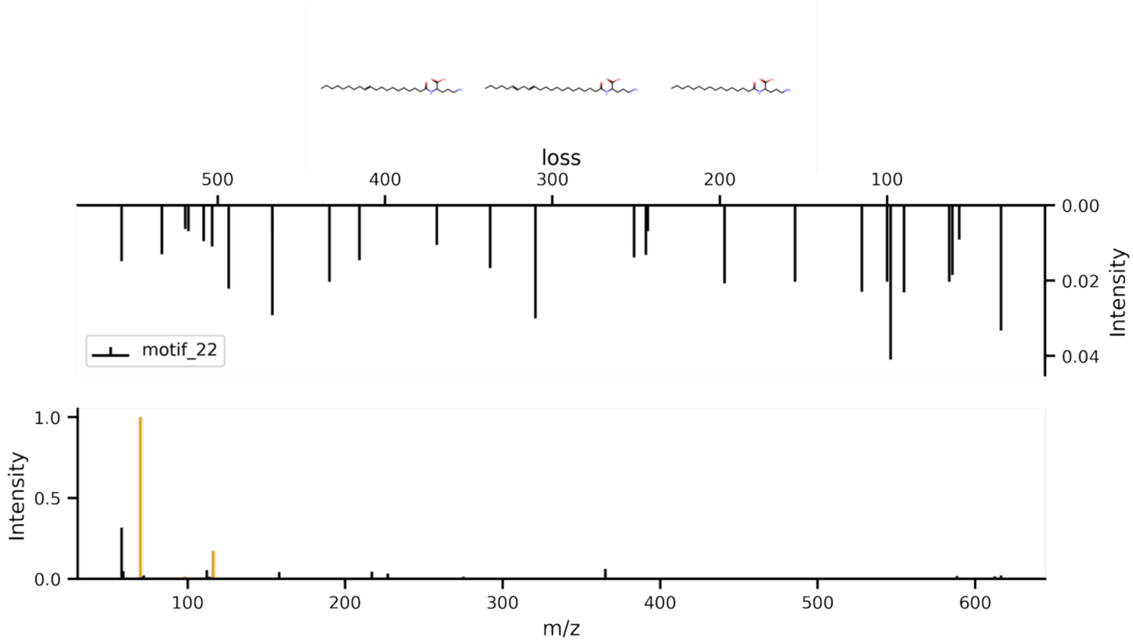

**Fig. S21**. *Mass2Motif 22 is associated with a 2 (propylidenamino-) acetic acid moiety. The top panel shows the only structure from MAG. The bottom panel displays the fragments associated with this Mass2Motif as a standard MS/MS spectrum. The orange-peaks highlighted 70.07 and 116.11 m/z are consistently observed in the recommended structures. Mirrored above the fragments are the corresponding neutral losses, arranged from right to left in increasing mass difference from the precursor ion.*

**Table S8.** *Results from the query Mass2Motif 108 against MotifDB*

| motifset | motif_id | short_annotation | annotation |
|---|---|---|---|
| **MIADB_pos_100** | motif_19 | Indole_in_mostly_corynanthean_series | Indole_in_mostly_corynanthean_series |
| **MIADB_pos_100** | motif_42 | Vobasane-containing-bisindole alkaloids | Vobasane-containing-bisindole alkaloids |
| **Urine derived Mass2Motifs** | motif_122 | Paracetamol mercapturate related (i.e. metabolites containing N-acetylcysteine adduct with paracetamol) | Paracetamol mercapturate related (i.e. metabolites containing N-acetylcysteine adduct with paracetamol) |
| **MIADB_pos_100** | motif_51 | hydroxylated or methoxylated Indole_Type1__MIAskeleton | hydroxylated or methoxylated Indole_Type1__MIAskeleton |
| **Urine derived Mass2Motifs** | motif_177 | Drug or xenobiotic - present in three urines and both sulphation and glucuronidation of core structure | Drug or xenobiotic - present in three urines and both sulphation and glucuronidation of core structure |

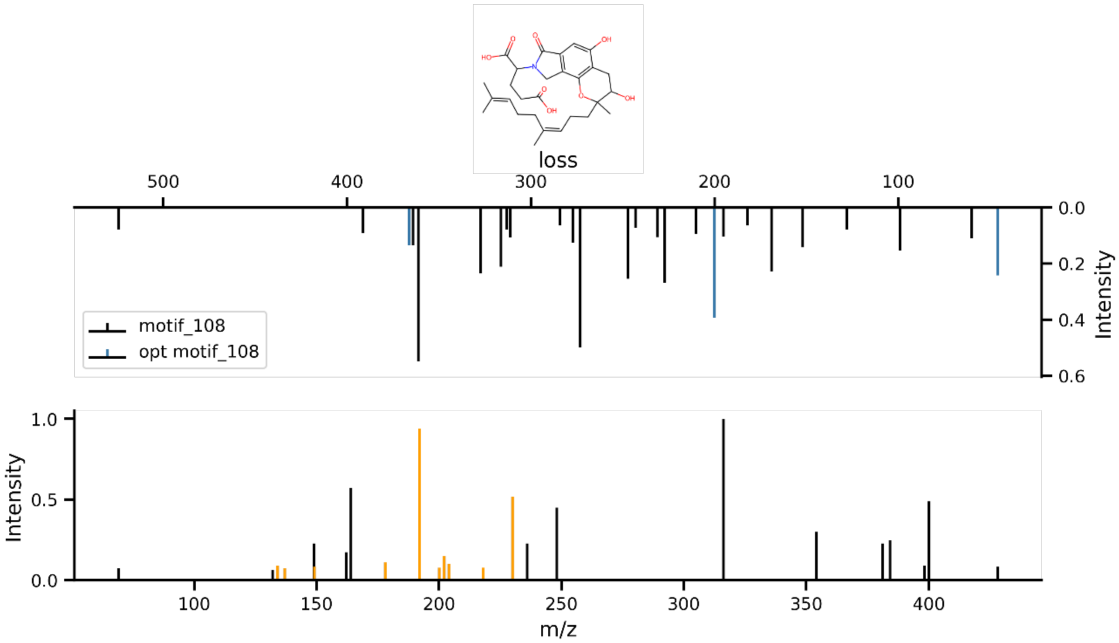

**Fig. S22**. *Mass2Motif 108 is associated with an indole substructure. The top panel shows the only structure from MAG. The bottom panel displays the fragments associated with this Mass2Motif as a standard MS/MS spectrum. The orange-multiple highlighted fragments are consistently observed in the recommended structure. Mirrored above the fragments are the corresponding neutral losses, arranged from right to left in increasing mass difference from the precursor ion.*

#### 3.4 Additional Mass2Motif annotations via motif-motif matching

In addition to the family compounds described in the original paper by Khatib *et al.*[4], we were able to annotate additional compounds present in the dataset. Using the MS2LDAViz 2.0 app, we computed the similarity between the inferred Mass2Motifs and MotifDB-MotifSets derived from GNPS, MassBank and LDB (Lichen DataBase - positive ionization mode mass spectra). These are the full descriptions of the MotifSets used:

**Table S9.** *Description of MotifSets used for motif-matching for the medicinal mushroom dataset results*

| Motifset Name | Feature Set | Description | No. motifs |
|---|---|---|---|
| GNPS library | 0.005 Da | MS/MS spectra obtained from reference compounds and isolated molecules from diverse sources with a focus on bacterial and plant related molecules - positive ionisation mode mass spectrometry on various instruments. | 78 |
| MassBank | 0.005 Da | MS/MS spectra obtained from reference compounds and isolated molecules from diverse sources with a slight focus on plant related molecules - positive ionisation mode mass spectrometry on various instruments | 46 |
| LDB MotifDB POS | 0.01 Da | MotifDB produced by the positive mode spectra of the LDB (250+ compounds with 745 spectra including different adducts and acquisition from three LC-MS instruments: Agilent 6530, Thermo Q-Exactive Focus, Waters Xevo G2-XS). Spectra were annotated with SMILES for MAGMa and substructure annotation. The chemical classes are from Huneck&Yoshimura (1996), and were used to annotate each motif. Parameters: 100 free motifs - binning of 0.01 Da | 100 |

The Mass2Motifs with a cosine similarity above 0.7, are displayed in Table S7. Most of the MAG-recommended structures align closely with the manual annotations from MotifDB. For example, Mass2Motif 10 matched Mass2Motif 13 and 11 with scores of 0.77 from Massbank and GNPS MotifSets respectively, both annotated as “Alkyl aromatic substructure indicative for aromatic ring”, which correlates with the MAG results shown in Fig. S24. Another good example was Mass2Motif 126 (Fig. S25), where the MAG recommendation correlates with the manual annotation of phenylalanine-like, with a high score of 0.757 against Mass2Motif 37 (from MassBank) and Mass2Motif 59 (from GNPS). Mass2Motif 61 is also noteworthy, as it is associated with a “fragment ion indicative of the presence of a phosphate group.” The MAG recommended structure clearly includes the phosphate group, as illustrated in Fig. S26. Not all the results completely align with SAG; for example, Mass2Motif 156 (Fig. S27). Although MAG recommendation does not exactly match with the manual annotation; further inspection reveals that one of the losses (161.1 m/z) could indicate the presence of a glycosylation (Fig. S28). This highlights the continued need for expert knowledge and curated annotation for some Mass2Motifs. It is important to notice that all the Mass2Motif found using a threshold over 0.7 show the same annotation between p MotifSet GNPS and MassBank, many motifs between these two sets overlap, since some of compounds and compound classes overlap between the spectral libraries.

**Table S10**. *Screening results from the App MS2LDA, of the mushroom dataset against GNPS, MassBank and LDB-positive MotifSets. The threshold was set as similarity above 0.7.*

| user_motif_id | user_auto_annotation | ref_motif_id | ref_short_annotation | ref_motifset | score |
|---|---|---|---|---|---|
| **motif_10** | COc1ccc(C(=O)c2ccccc2-c2ccc3c(c2)S(=O)(=O)Cc2cc(C(=O)NCc4ccc(C)cc4)sc2-3)cc1, O=C(O)CC(=O)O, CC(NC(=O)C(C#N)=Cc1cccc(Br)n1)c1ccccc1 | motif_13 | Alkyl aromatic substructure, indicative for aromatic ring with alkyl chain | Massbank library derived Mass2Motifs | 0.7742 |
| **motif_10** | COc1ccc(C(=O)c2ccccc2-c2ccc3c(c2)S(=O)(=O)Cc2cc(C(=O)NCc4ccc(C)cc4)sc2-3)cc1, O=C(O)CC(=O)O, CC(NC(=O)C(C#N)=Cc1cccc(Br)n1)c1ccccc1 | motif_11 | Alkyl aromatic substructure, indicative for aromatic ring with alkyl chain | GNPS library derived Mass2Motifs | 0.7742 |
| **motif_156** | CCC(=O)OC(CC(=O)[O-])C[N+](C)(C)C, CCC(=O)OC(CC(=O)O)C[N+](C)(C)C | motif_45 | Fragments indicative of a glycosylation, i.e. indicative of a glucose or galactose conjugation | GNPS library derived Mass2Motifs | 0.7671 |
| **motif_156** | CCC(=O)OC(CC(=O)[O-])C[N+](C)(C)C, CCC(=O)OC(CC(=O)O)C[N+](C)(C)C | motif_24 | Fragments indicative of a glycosylation, i.e. indicative of a glucose or galactose conjugation | Massbank library derived Mass2Motifs | 0.7671 |
| **motif_126** | NC(Cc1ccccc1)C(=O)O | motif_59 | Fragments indicative for [phenylalanine-CHOOH] based substructure | GNPS library derived Mass2Motifs | 0.7574000000000001 |
| **motif_126** | NC(Cc1ccccc1)C(=O)O | motif_37 | Fragments indicative for [phenylalanine-CHOOH] based substructure | Massbank library derived Mass2Motifs | 0.7574000000000001 |
| **motif_61** | CCCCCCCCCCCCOP(=O)(O)O, O=P(O)(O)OCCC(F)(F)C(F)(F)C(F)(F)C(F)(F)C(F)(F)C(F)(F)C(F)(F)C(F)(F)F, O=P(OC(CCl)CCl)(OC(CCl)CCl)OC(CCl)CCl | motif_48 | Fragment ion indicative for the presence of a phosphate group | GNPS library derived Mass2Motifs | 0.7079000000000001 |
| **motif_61** | CCCCCCCCCCCCOP(=O)(O)O, O=P(O)(O)OCCC(F)(F)C(F)(F)C(F)(F)C(F)(F)C(F)(F)C(F)(F)C(F)(F)C(F)(F)F, | motif_26 | Fragment ion indicative for the presence of a phosphate group | Massbank library derived Mass2Motifs | 0.7079000000000001 |

| | | | | | |
|---|---|---|---|---|---|
| | O=P(OC(CCl)CCl)(OC(CCl)CCl)OC(CCl)CCl | | | | |
| **motif_148** | CN(C)CCCCC(N)C(=O)O, NCCCCC(N)C(=O)O | motif_57 | Fragment ions indicative for alkylamine substructure C5H10N | GNPS library derived Mass2Motifs | 0.7067 |
| **motif_148** | CN(C)CCCCC(N)C(=O)O, NCCCCC(N)C(=O)O | motif_35 | Fragment ions indicative for alkylamine substructure C5H10N | Massbank library derived Mass2Motifs | 0.7067 |

##### Motif Details: motif_10

###### Spec2Vec Matching Results

The Spec2Vec matching results displayed here suggest chemical structures (SMILES strings) that closely match the selected motif. Spec2Vec calculates similarities by comparing motif pseudo-spectra against reference spectra from a known database. Matches shown here can help identify possible chemical identities or provide clues about structural characteristics represented by this motif.

###### Spec2Vec Matching Results

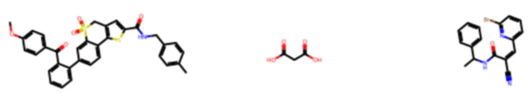

Match 1 Match 2 Match 3

Auto Annotations: ['COc1ccc(C(=O)c2ccccc2-c2ccc3c(c2)S(=O)(=O)Cc2cc(C(=O)NCc4ccc(C)cc4)sc2-3)cc1', 'O=C(O)CC(=O)O', 'CC(NC(=O)C(C#N)=Cc1cccc(Br)n1)c1ccccc1']

###### Screening Annotations

| Ref Motif ID | Ref ShortAnno | Ref MotifSet | Score |
|---|---|---|---|
| motif_13 | Alkyl aromatic substructure â€“ indicative for aromatic ring... | Massbank library derived Mass2Motifs | 0.7742 |
| motif_11 | Alkyl aromatic substructure â€“ indicative for aromatic ring... | GNPS library derived Mass2Motifs | 0.7742 |
| motif_23 | No short annotation available | LDB MotifDB POS | 0.1723 |
| motif_86 | No short annotation available | LDB MotifDB POS | 0.1473 |
| motif_50 | Steroid core related (C18H21 and smaller fragments thereof -... | GNPS library derived Mass2Motifs | 0.1460 |

**Fig. S23**. *Mass2Motif 10 details shown using the app. The matching results show the structure recommendations from MAG. The bottom panel displays the Mass2Motif-matching results, with their Spec2Vec respective scores. This inspection shows that it is highly probable that Mass2Motif 10 contains an aromatic ring as MAG recommendation and Mass2Motif-matching align.*

#### Motif Details: motif_126

##### Spec2Vec Matching Results

The Spec2Vec matching results displayed here suggest chemical structures (SMILES strings) that closely match the selected motif. Spec2Vec calculates similarities by comparing motif pseudo-spectra against reference spectra from a known database. Matches shown here can help identify possible chemical identities or provide clues about structural characteristics represented by this motif.

##### Spec2Vec Matching Results

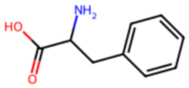

Match 1

Auto Annotations: ['NC(Cc1ccccc1)C(=O)O']

###### Screening Annotations

| Ref Motif ID | Ref ShortAnno | Ref MotifSet | Score |
|---|---|---|---|
| motif_59 | Fragments indicative for [phenylalanine-CHOOH] based substru... | GNPS library derived Mass2Motifs | 0.7574 |
| motif_37 | Fragments indicative for [phenylalanine-CHOOH] based substru... | Massbank library derived Mass2Motifs | 0.7574 |
| motif_91 | No short annotation available | LDB MotifDB POS | 0.6007 |
| motif_97 | No short annotation available | LDB MotifDB POS | 0.1407 |
| motif_85 | No short annotation available | LDB MotifDB POS | 0.1319 |

**Fig. S24**. *Mass2Motif 126 details shown using the app. The matching results show the only structure recommendation from MAG. The bottom panel displays the Mass2Motif-matching results, with their Spec2Vec respective scores. This inspection shows the MAG and Mass2Motif recommendation align, assigning Mass2Motif 126 phenylalanine containing-substructure.*

#### Motif Details: motif_61

##### Spec2Vec Matching Results

The Spec2Vec matching results displayed here suggest chemical structures (SMILES strings) that closely match the selected motif. Spec2Vec calculates similarities by comparing motif pseudo-spectra against reference spectra from a known database. Matches shown here can help identify possible chemical identities or provide clues about structural characteristics represented by this motif.

##### Spec2Vec Matching Results

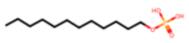

Match 1

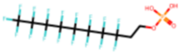

Match 2

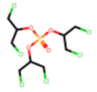

Match 3

Auto Annotations: ['CCCCCCCCCCCCOP(=O)(O)O', 'O=P(O)(O)OCCC(F)(F)C(F)(F)C(F)(F)C(F)(F)C(F)(F)C(F)(F)C(F)(F)C(F)(F)F', 'O=P(OC(CCl)CCl)(OC(CCl)CCl)OC(CCl)CCl']

###### Screening Annotations

| Ref Motif ID | Ref ShortAnno | Ref MotifSet | Score |
|---|---|---|---|
| motif_48 | Fragment ion indicative for conjugation of a phosphate group... | GNPS library derived Mass2Motifs | 0.7079 |
| motif_26 | Fragment ion indicative for the presence of a phosphate grou... | Massbank library derived Mass2Motifs | 0.7079 |
| motif_1 | Fragment of C5H10NO2 and loss of C5H9NO2 (in beer likely to ... | Massbank library derived Mass2Motifs | 0.1527 |
| motif_4 | Fragment and loss of [proline-H2O] - indicative for conjugat... | GNPS library derived Mass2Motifs | 0.1527 |
| motif_90 | No short annotation available | LDB MotifDB POS | 0.1249 |

**Fig. S25**. *Mass2Motif 61 details shown using the app. The matching results show the structure recommendations from MAG. All of them contain a phosphate group. The bottom panel displays the Mass2Motif-matching results, with their Spec2Vec respective scores. This inspection shows the MAG and Mass2Motif positively align, assigning Mass2Motif 126 phosphate containing-substructure.*

#### Motif Details: motif_156

##### Spec2Vec Matching Results

The Spec2Vec matching results displayed here suggest chemical structures (SMILES strings) that closely match the selected motif. Spec2Vec calculates similarities by comparing motif pseudo-spectra against reference spectra from a known database. Matches shown here can help identify possible chemical identities or provide clues about structural characteristics represented by this motif.

##### Spec2Vec Matching Results

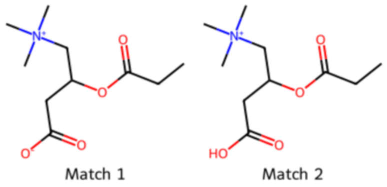

Auto Annotations: ['CCC(=O)OC(CC(=O)[O-])C[N+](C)(C)C', 'CCC(=O)OC(CC(=O)O)C[N+](C)(C)C']

##### Screening Annotations

| Ref Motif ID | Ref ShortAnno | Ref MotifSet | Score |
|---|---|---|---|
| motif_45 | Fragments indicative of a glycosylation â€“ i.e. indicative ... | GNPS library derived Mass2Motifs | 0.7671 |
| motif_24 | Fragments indicative of a glycosylation â€“ i.e. indicative ... | Massbank library derived Mass2Motifs | 0.7671 |
| motif_23 | No short annotation available | LDB MotifDB POS | 0.2647 |
| motif_42 | Fragments indicative for kaempferol/glycosylated kaempferol ... | Massbank library derived Mass2Motifs | 0.2430 |
| motif_39 | No short annotation available | LDB MotifDB POS | 0.1678 |

**Fig. S26**. *Mass2Motif 156 details shown using the app. The matching results show the structure recommendations from MAG. The structures shown are identical, containing an methylammonium group and a carboxylic group. The bottom panel displays the Mass2Motif-matching results, with their Spec2Vec respective scores. This inspection shows the MAG and Mass2Motif do not align, as the manual annotation assign a glycosylation for this Mass2Motif.*

##### Motif Features Table and Summary Plots

The table below lists the motif features (fragments and losses) that pass the probability filter, including their probabilities within the motif. Below it, a bar plot shows how frequently each feature appears **within the filtered set of documents** for this motif (i.e., documents whose doc-topic probability and overlap score both pass the current threshold ranges).

##### Motif Features Table

| Feature | Probability |
|---|---|
| frag@85.03 | 0.4927 |
| loss@27.96 | 0.0404 |
| loss@133.11 | 0.0398 |
| loss@161.1 | 0.0340 |
| frag@57.03 | 0.0269 |
| frag@145.05 | 0.0247 |
| frag@60.08 | 0.0238 |
| loss@366.24 | 0.0196 |
| loss@234.07 | 0.0196 |
| frag@323.26 | 0.0196 |

**Fig. S27**. *Mass2Motif 156 inspection suing the app. The table shows the fragments and losses representing this Mass2Motif. This Mass2Motig is represented by a loss of 161.1 m/z, that could be connected to a glycosylation.*

### 4. Suspect list

In the case study on the suspect list, four examples are presented where at least one Mass2Motif was linked to an identified suspect, or more precisely to a substructure of the identified suspect. The information provided in 4.1-4.4 includes the topic-document probabilities, CFM-ID explanations, and MassBank reference spectra to confirm the findings and links between Mass2Motifs, the MAG, and manual identified suspects.

#### 4.1 Peptide example

Fig. S28 shows two peptide-related Mass2Motifs with probabilities of 0.12 and 0.29 being linked to a suspect peptide. The two Mass2Motifs shown had the highest probability of being associated with the suspect structure. Table S11 shows the fragments and losses of each motif and its CFM-ID[6] annotation as a SMILES to confirm the MAG predictions.

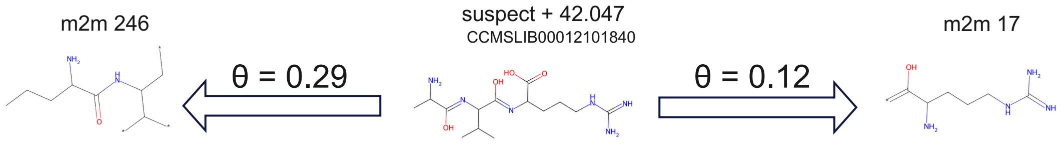

**Fig. S28.** *Topic-document probability between Mass2Motifs 246 and 17 (left and right) and suspect (center). This shows how the Mass2Motifs represent different substructures of the complete suspect molecule.*

**Table S11.** *CFM-ID 4.0 calculation for the peptide suspect with id = spec_20879. Used InChI for CFM-ID calculations: i= 'InChI=1S/C14H28N6O4/c1-7(2)10(20-11(21)8(3)15)12(22)19-9(13(23)24)5-4-6-18-14(16)17/h7-10H,4-6,15H2,1-3H3,(H,19,22)(H,20,21)(H,23,24)(H4,16,17,18) - 42.047 Da'*

| Frag or Loss | CFM-ID SMILES | Mass2Motif 17 | Mass2Motif 246 |
|---|---|---|---|
| Frag@60.06 | NC(N)=[NH2+] | NO | YES |
| Frag@70.07 | CC(C)C#[NH+] | YES | YES |
| Frag@72.08 | CC(C)C=[NH2+] | YES | YES |
| Frag@143.12 | C=C([NH3+])C(=O)NC C(C)C | YES | NO |
| Frag@175.12 | NC(N)NCC=CC(=[NH2 +])C(O)O | NO | YES |
| Loss@17.03 | NH3 | NO | YES |

#### 4.2 Steroid example

Fig. S29 shows to Mass2Motifs being related with probabilities of 0.13 and 0.23 with the steroid suspect, which represent the two most associated Mass2Motifs with the suspect structure. Table S12 explains the fragments of the Mass2Motifs by CFM-ID annotations, which can confirm the MAG substructure prediction.

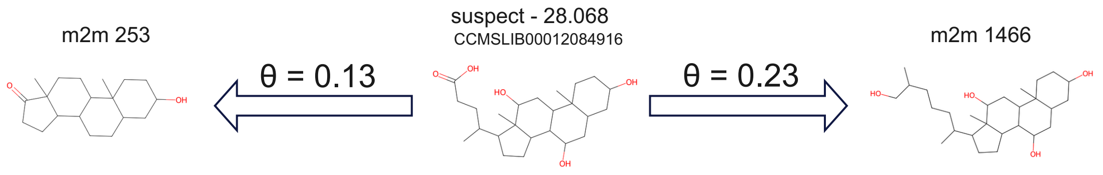

**Fig. S29.** *Topic-document probability between Mass2Motifs 253 and 1466 (left and right) and suspect (center). This shows how the Mass2Motif 1466, which covers most of the suspect molecule, shows higher probability than Mass2Motif 253, which also contains a steroid substructure.*

**Table S12.** *CFM-ID 4.0 calculation for the steroid suspect. Used InChI for CFM-ID calculations: InChI=1S/C24H40O5/c1-13(4-7-21(28)29)16-5-6-17-22-18(12-20(27)24(16,17)3)23(2)9-8-15(25)10-14(23)11-19(22)26/h13-20,22,25-27H,4-12H2,1-3H3,(H,28,29)/t13-,14+,15-,16-,17+,18+,19+,20+,22+,23+,24-/m1/s1*

| Frag or Loss | CFM-ID SMILES | Mass2Motif 1466 | Mass2Motif 253 |
|---|---|---|---|
| Frag@133.1 | CCC(C)CCC(O)[OH2+] | YES | NO |
| Frag@145.1 | C=CCC(C)CCC(O)[OH2+] | YES | NO |
| Frag@147.12 | CCCC(C)CCC(O)[OH2+] | YES | NO |
| Frag@157.1 | CC1CCC(O)CC1CC=[OH+] | YES | NO |
| Frag@159.12 | CC1CCC(O)CC1CC[OH2+] | YES | NO |
| Frag@161.13 | CC12C=CC=CC1=CC(=[OH+])CC2 | YES | NO |
| Frag@185.13 | CC12CCC(O)CC1CC([OH2+])CC2 | YES | NO |
| Frag@187.15 | CC1CCCC1C(C)CCC(O)[OH2+] | YES | NO |
| Frag@199.15 | CC1CC([OH2+])CC2CC(O)CCC12C | YES | NO |
| Frag@213.16 | CC(CCC(O)[OH2+])C1=CC=CC1(C)CO | YES | NO |
| Frag@215.16 | CC(CCC(O)[OH2+])C1CC=CC1(C)CO | YES | NO |
| Frag@227.18 | CC12CCC(O)CC1CC(O)CC2CC=[OH+] | YES | NO |
| Frag@365.32 | CCC(C)C1CCC2C3C(O)CC4CC(O)CCC4(C)C3CC([OH2+])C12C | NO | YES |

#### 4.3 Sulfonamide example

Fig. S30 show one Mass2Motif representing a core substructure of the sulfonamide suspect. The probability of the Mass2Motif being associated with the suspect by the LDA model is 0.22. Table S13 shows annotations of fragments that are present in the Mass2Motif. A fragment present in the Mass2Motif couldn't be explained by CFM-ID but could be linked to the sulfonamide substructure by the molecular formula from a structure analog in MassBank (Fig. S31).

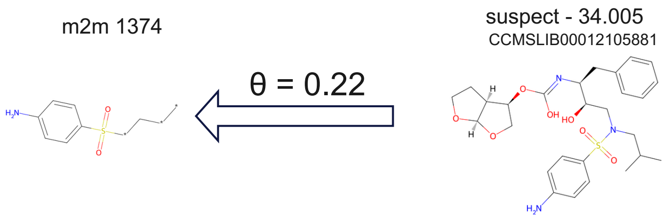

**Fig. S30.** *Topic-document probability between Mass2Motifs 1374 (left) and suspect (right). This shows how the Mass2Motif 1374 represents part of the suspect structure with an aromatic ring attached to a sulfone functional group.*

**Table S13.** *CFM-ID 4.0 calculation for the sulfonamide suspect. Used InChI for CFM-ID calculations: InChI=1S/C27H37N3O7S/c1-18(2)15-30(38(33,34)21-10-8-20(28)9-11-21)16-24(31)23(14-19-6-4-3-5-7-19)29-27(32)37-25-17-36-26-22(25)12-13-35-26/h3-11,18,22-26,31H,12-17,28H2,1-2H3,(H,29,32)/t22-,23-,24+,25-,26+/m0/s1*

| Frag or Loss | CFM-ID SMILES | Mass2Motif 1374 |
|---|---|---|
| Frag@92.05 | C#CCC#CC=[NH2+] | YES |
| Frag@108.04 | / | YES |
| Frag@156.01 | [NH2+]=C1C=CC(=S(=O)=O)C=C1 | YES |

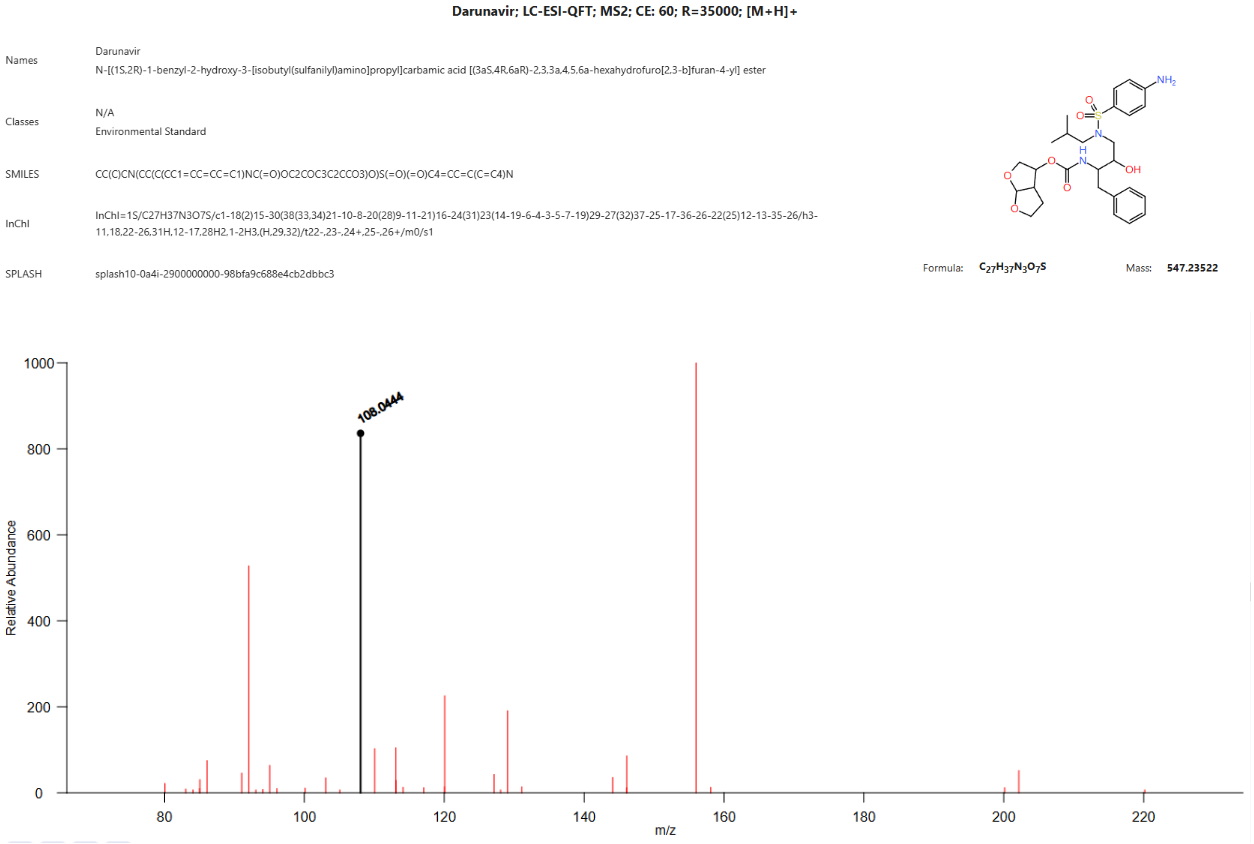

**Fig. S31.** *Massbank spectra with fragment m/z 108.04 with a calculated molecular formula of C6H6NO+. The compound contains a sulfonamide moiety, which is based on the molecular formula responsible for the fragment 108.04.*

#### 4.4 Glycoside example

Fig. S32 show a glucose substructure related Mass2Motif being related with a probability of 0.77 to a glycoside suspect. The overlapping loss of 162.05 and the m/z difference of 18.01 are known losses corresponding to glucose and water, respectively[7].

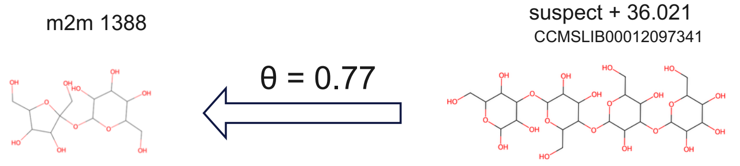

**Fig. S32.** *Topic-document probability between Mass2Motifs 1388 (left) and suspect (right). This shows how the Mass2Motif 1388 a sugar moiety from the complete suspect molecule.*

### 5. Integration and handling large datasets

#### 5.1 Handling large datasets within reasonable timeframes

One of the main improvements in MS2LDA 2.0 is its runtime performance. This enhancement is partly due to the use of multiple cores during model execution, a feature enabled by the Tomotopy[10] package. We confirmed this by comparing the running performance between the original version and MS2LDA 2.0. In the original version only one single core can be used to process data, so firstly we compared MS2DA original version and MS2LDA 2.0 using one single core. Fig. S33 shows MS2LDA 2.0 outperforming the original version for small size libraries (Fig. S33, green dot and squares) with only 1,267, whereas for larger libraries the difference between the two models is reduced, because both implementations become memory-bandwidth bound on a single core, making the optimized C++ routines in Tomotopy less advantageous over NumPy's compiled operations in the original implementation. For instance, we observed for the two largest spectral libraries, the Suspect-Library and BERKELEY-LAB-Library, how the runtime was nearly identical between MS2LDA 2.0 and its predecessor (Fig. S33, blue dots and squares).

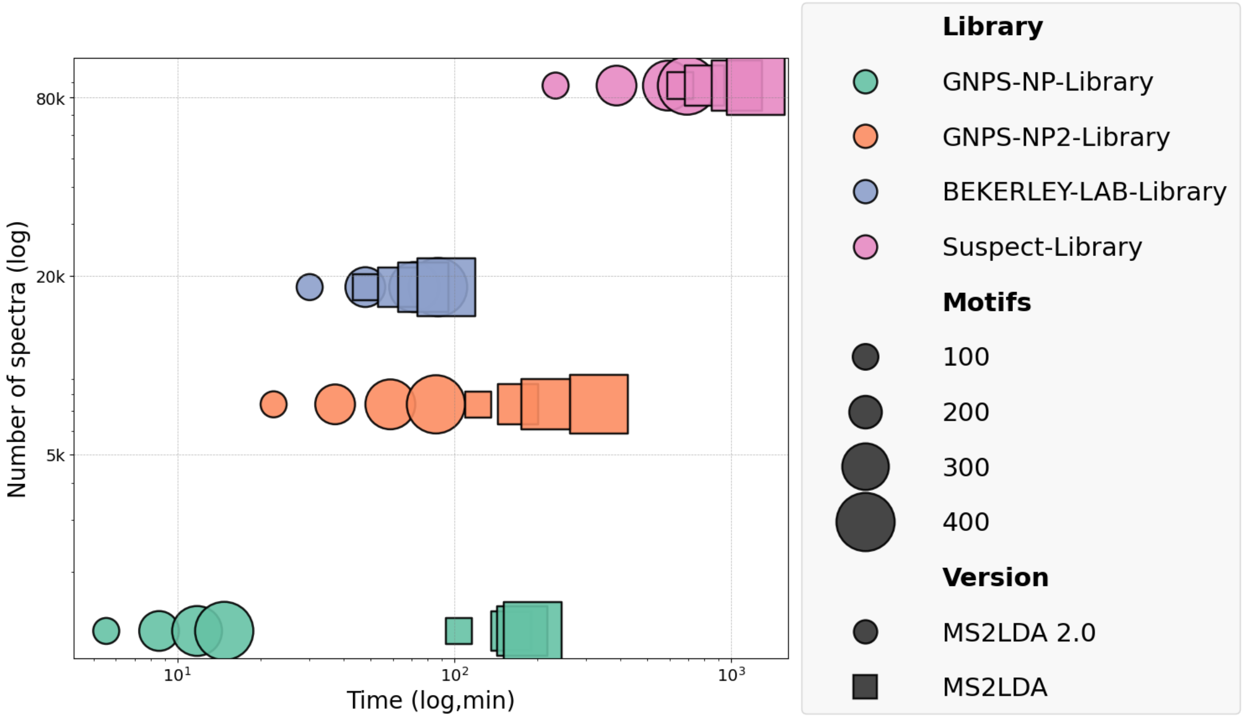

**Fig. S33.** *Comparison of speed performance between MS2LDA and MS2LDA 2.0 across four different libraries: GNPS-NP (1,267 spectra), GNPS-NP2 (7,377 spectra), BERKELEY-LAB (18,352 spectra) and Suspect-List (87,899 spectra), using single core by AMD EPYC 7532 32-Core Processor. Running time is plotted on the x-axis and number of spectra on the y-axis. Software versions are distinguished by shape; number of motifs are represented by size and libraries are indicated by colour.*

However, when using MS2LDA 2.0 with multiple cores, its implementation outperforms the original version in each of the datasets (Fig S34). Moreover, with four cores (that most laptops have available nowadays), datasets containing fewer than 20,000 spectra can be processed in under an hour, and processing time can be further reduced with more cores or additional server resources.

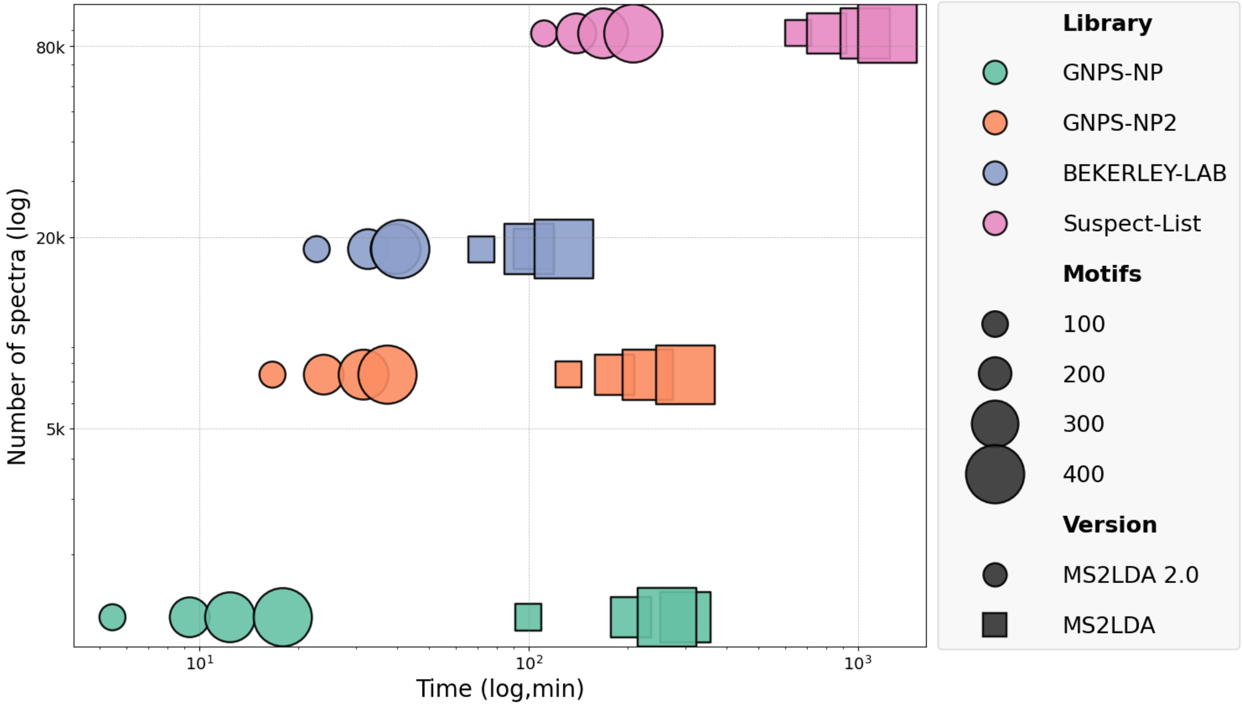

**Fig. S34.** *Comparison of speed performance between MS2LDA and MS2LDA 2.0 across four different libraries: GNPS-NP (1,267 spectra), GNPS-NP2 (7,377 spectra), BERKELEY-LAB (18,352 spectra) and Suspect-List (87,899 spectra), using 4-cores by AMD EPYC 7532 32-Core Processor. Running time is plotted on the x-axis and number of spectra on the y-axis. Software versions are distinguished by shape, number of motifs are represented by size and libraries are indicated by colour.*

#### 5.2 Mass2Motif Object

To be able to store all information needed for Mass2Motifs, we created a Mass2Motif object. The Mass2Motif object is similar to the matchms spectrum object, but it has been changed in a few functions to be able to work with Mass2Motifs instead of mass spectra. First, Mass2Motifs can consist of only losses and therefore the mandatory fragments needed for matchms spectrum objects had to be replaced. The second difference is that Mass2Motifs do not have a precursor mass and therefore associated losses with a Mass2Motif need to be stored since they cannot be calculated. All metadata as well as fragments and losses are retrieved in the same way as it is done in the matchms framework. This allows users to use most tools (for similarity search or other aims) that were developed with the matchms framework also for Mass2Motifs.

**Table S14.** *Comparison between Mass2Motif objects and Matchms Spectrum objects. The main difference between Mass2Motif objects and matchms spectrum objects is that Mass2Motifs don't need fragments and losses can be stored within the object.*

| | **MS2LDA 2.0 Mass2Motif object** | **Matchms Spectrum object** |
|---|---|---|
| **Contains fragments** | Optional | Mandatory |
| **Contains losses** | Optional | No (calculated based on precursor mass) |
| **Access fragments** | .peaks.mz | .peaks.mz |
| **Access losses** | .losses.mz (retrieved) | .lossses.mz (calculated based on precursor mass) |
| **Access metadata** | .get("metadata") | .get("metadata") |

### 5.3 MotifDB

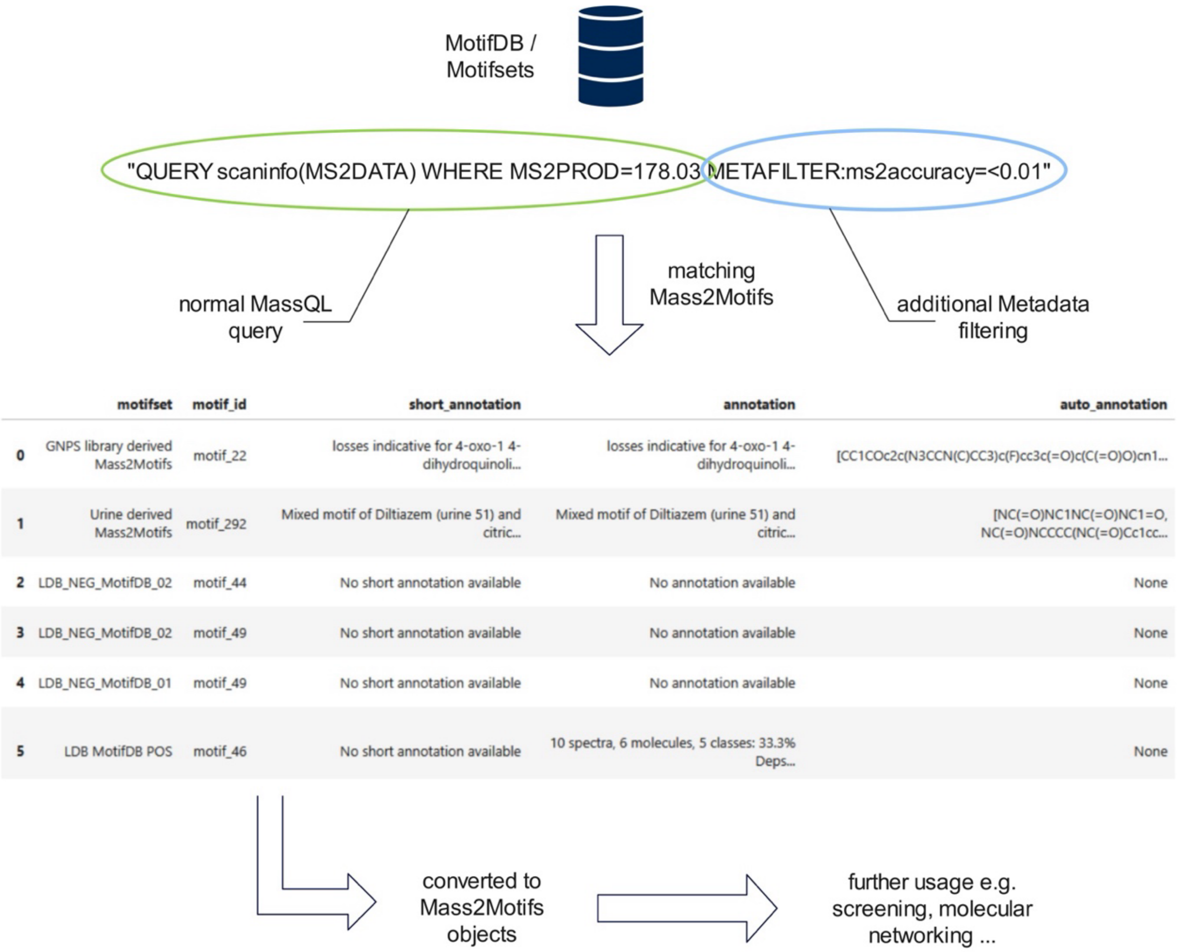

**Fig. S35.** *Graphical MotifDB workflow starting by querying MotifDB via MassQL, then converting it to a Mass2Motif object that can be used for molecular networking or spectra screening or Mass2Motif matching tasks.*

To query MotifDB, we developed MassQueryLanguage4Mass2Motifs (https://github.com/j-a-dietrich/MassQueryLanguage4Mass2Motifs), a fork of MassQL[8]. MassQL provides a framework to query spectra but does not meet the requirements for querying Mass2Motifs. The major difference between spectra and Mass2Motifs is that Mass2Motifs do not possess a precursor ion and therefore no losses can be calculated. Since MassQL calculates losses in-time based on precursor ions (precursor ion - fragments), it cannot be used for Mass2Motifs. MassQL4Mass2Motifs stores losses permanently without the need for in-time calculations. Furthermore, Mass2Motifs are often associated with metadata such as annotations (manual or automated), instrument type, MassiveID, and more. We integrated the possibility to query not only for fragments and losses but also for the metadata by using the key word METAFILTER:columnname=value. Metafilters can be combined and should be listed at the end of the query. Finally, we not only want to find Mass2Motifs of interest but also to use them further for screening purposes or molecular networking by using Mass2Motifs as nodes or edges between spectra. Therefore, we integrated a function to convert queried Mass2Motifs back to Mass2Motif objects. These Mass2Motif objects can then be easily integrated into common metabolomics workflows, for example to be used in MolNetEnhancer.

**Table S15:** *Comparison between the MassQL format and the modified version for MotifDB. The main difference is the storage of losses and metadata, which both can be queried as well.*

| | **MotifDB** | **MassQL** |
|---|---|---|
| **Contains fragments** | Optional | Mandatory |
| **Contains losses** | Optional | No (calculated based on precursor mass) |
| **Query MS1 data** | No | Yes, massql syntax |
| **Query MS2 data** | Yes, massql syntax | Yes, massql syntax |
| **Query metadata** | Yes, metafiler:column=value | No |

### 6. MS2LDAViz 2.0: Visualization Interface

A new and revamped MS2LDAViz 2.0 interface has been developed as part of MS2LDA 2.0, providing users with an intuitive web-based platform for exploring and interpreting Mass2Motifs. Built using the Dash framework[9], this visualization tool enables interactive browsing of Mass2Motifs, examination of automated annotations provided by Motif Annotation Guidance (MAG), and targeted searches of Mass2Motifs using MassQL-based queries. Compared to its predecessor, the new MS2LDAViz is easier to deploy and maintain, offers improved scalability, and allows users to run analyses locally on their computers or remotely on servers. This interface aims to streamline the annotation workflow, facilitate rapid exploration of complex MS/MS data, and promote community-driven annotation efforts through seamless integration with MotifDB.
We have made available an online version of the app running at ms2lda.org/viz that is able to load ms2lda 2.0 models to inspect and visualize them – it includes buttons to upload the pesticide and natural products case study models for interactive exploration of the in this study presented results. We note that if installed locally, the app also contains a “Run Analysis” tab (see section 6.1) that guides users in setting up an MS2LDA run.

The following sections describe different sections of the visualization app.

#### 6.1 Run Analysis

Fig S.36 shows the start screen of the visualization app. This screen allows users to initiate a new MS2LDA 2.0 analysis directly within the visualization application, provided that the application is installed locally with the necessary computational performance. Users can upload their MS/MS data file (supported formats include .mgf, .mzML and .msp) using the provided drag-and-drop interface. Basic MS2LDA 2.0 parameters such as the number of motifs, acquisition type (DDA or DIA), polarity, and iteration count can be adjusted directly. Additionally, Spec2Vec parameters are configurable from this screen; users can select and manage pre-trained Spec2Vec model and embedding files necessary for automatic motif annotation. Advanced settings (accessible by expanding the corresponding section) allow fine-grained control over preprocessing details, convergence criteria, and LDA model hyperparameters.

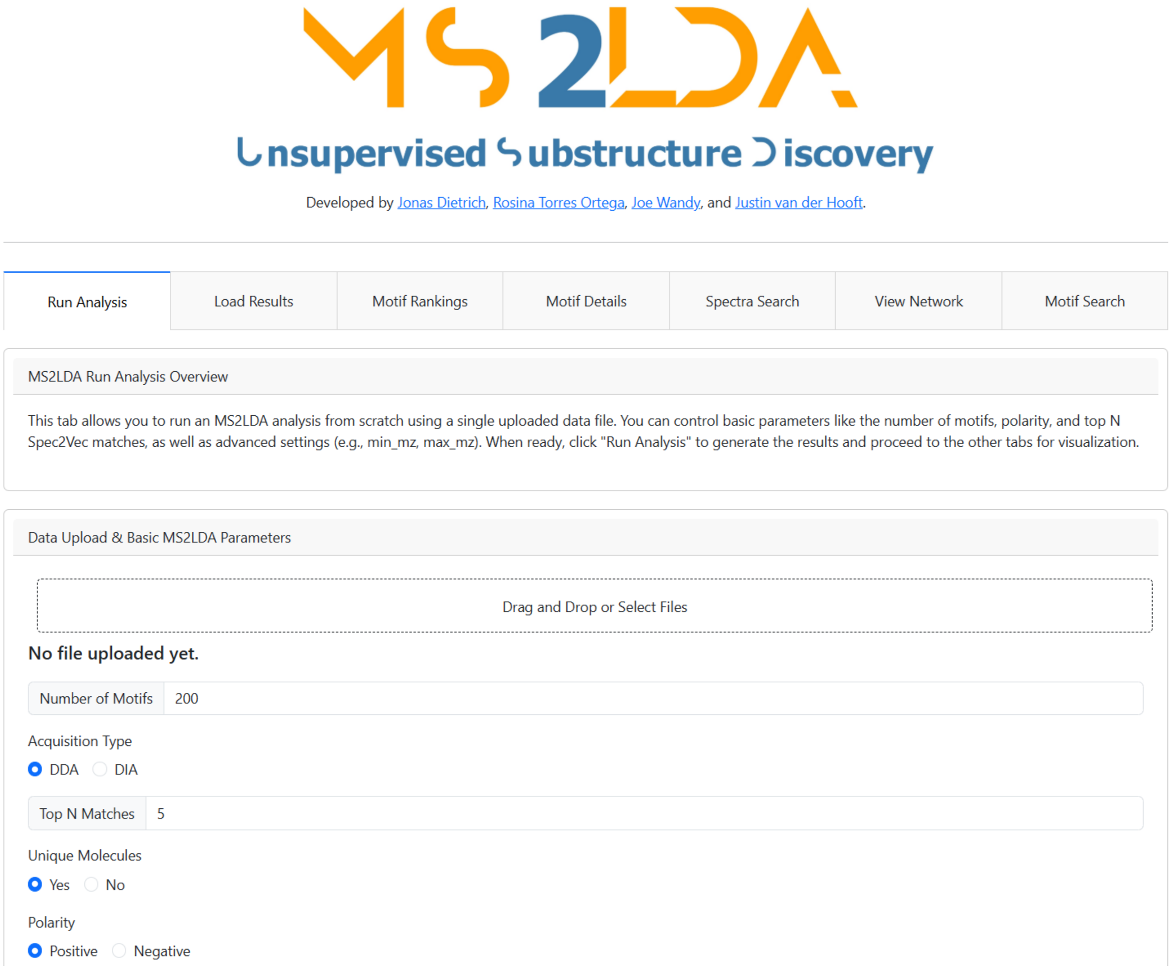

**Fig. S36.** *Example of the landing page from MS2LDA 2.0 app. It starts with the tab "Run Analysis", then after running, the results can be submitted at the "Load Results" section, "Motif Rankins" serves to generally exploring all the Mass2Motifs inferred in an experiments, once you select a motif, you can further inspect it at the "Motif Details" tab, on the "Spectra Search" you can explore spectra containing certain motifs, "View Network" serves to explore the connections between our Mass2Motifs, if they share some fragments and "Motif Search" .*

Once parameters are set, clicking "Run Analysis" initiates the modeling process. A folder containing all the results, including visualization results will be generated. By default, this folder is called "ms2lda_results" and the corresponding viz data file is ms2lda_viz.json.gz in that folder. This data file can be loaded for further analysis into the "Load Results" tab. This ensures that the running of MS2LDA 2.0 and the visualization and exploration of the results are separated.

#### 6.2 Motif Ranking and Motif Details

The “Motif Ranking” screen provides a ranked list of inferred Mass2Motifs based on their significance across the analyzed dataset. Each motif is characterized by several key metrics:

- Degree: The number of spectra (documents) that are associated with the Mass2Motif.
- Average Doc-Topic Probability: The average probability of this motif appearing in associated spectra.
- Average Overlap Score: An averaged measure indicating how distinctly this motif explains the associated spectra.
- Annotation: Automatic annotations provided by Motif Annotation Guidance (MAG), summarizing potential chemical structures matching the Mass2Motif.

Interactive sliders at the top allow users to set probability and overlap thresholds, dynamically filtering motifs to quickly identify the most relevant and specific ones. Clicking a motif identifier (e.g., motif_148) leads users directly to a detailed view, where more information such as specific spectra associations, MAG results, and detailed annotations can be further explored.

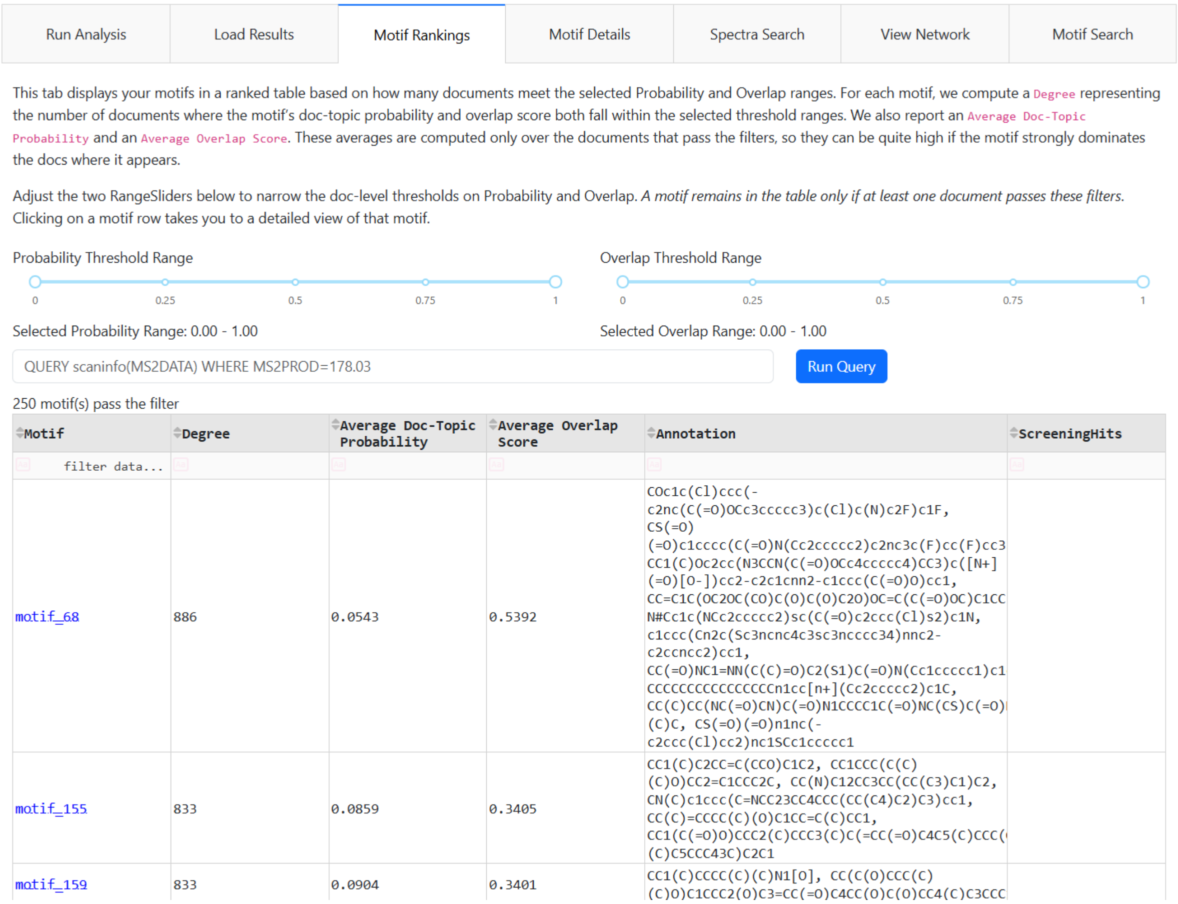

**Fig. S37.** *Mass2Motif exploration interface in the "Motif Rankins" tab of the application. All Mass2Motifs are presented in an interactive table for exploration, accompanied by summary statistics to facilitate interpretation. The Degree indicates the number of MS/MS spectra in which given Mass2Motif occurs. The Average Doc-Topic Probability reflects how strongly a Mass2Motif is associated with each MS/MS spectrum on average, higher value meaning that a motif is commonly present in the data. The Average Doc-Topic Probabilities shows how much overlap there is among the fragment/neutral loss in the Mass2Motifs, with lower values indicating more distinct and interpretable motifs.*

Once a motif is selected from the ranking table, its details are shown in the "Motif Details" screen. This screen provides comprehensive insights into a selected Mass2Motif, including automated structural annotations, spectral composition, and associated features. At the top, users see chemical structures suggested by Motif Annotation Guidance (MAG) based on Spec2Vec matching. Below this, a dual pseudo-spectrum plot compares optimized motif features with the raw fragments from associated spectra, facilitating interpretation and validation. The interface also lists individual fragment and loss features, highlighting their frequency and probability within the motif. Finally, users can interactively browse individual spectra linked to the motif, visualizing how each spectrum's fragments map onto the motif's characteristic features. This detailed breakdown allows for rapid, informed interpretation of discovered Mass2Motifs.

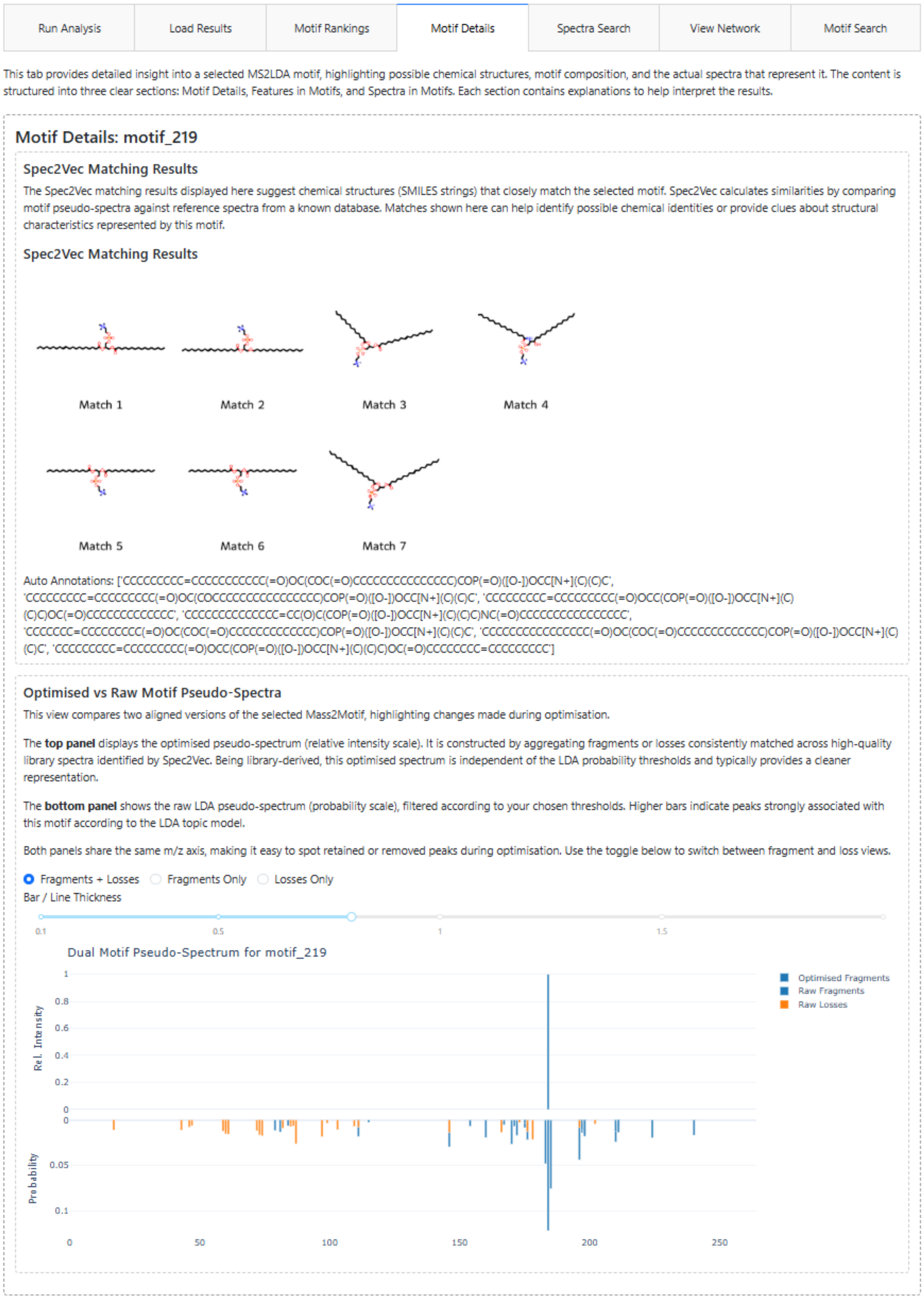

**Fig. S38**. *Mass2Motif exploration interface in the "Motif Details" tab of the application. This section shows in detail the Mass2Motif selected. On top the recommended structures by MAG are shown. Below, a mirror spectrum shows on the bottom the Mass2Motif as a standard MS/MS with the fragments and losses and their corresponding probabilities on the x axis. On top it shows the optimized-spectra Mass2Motif, where fragments or losses shown are present in all the structural MAG recommendations given above with their relative intensity.*

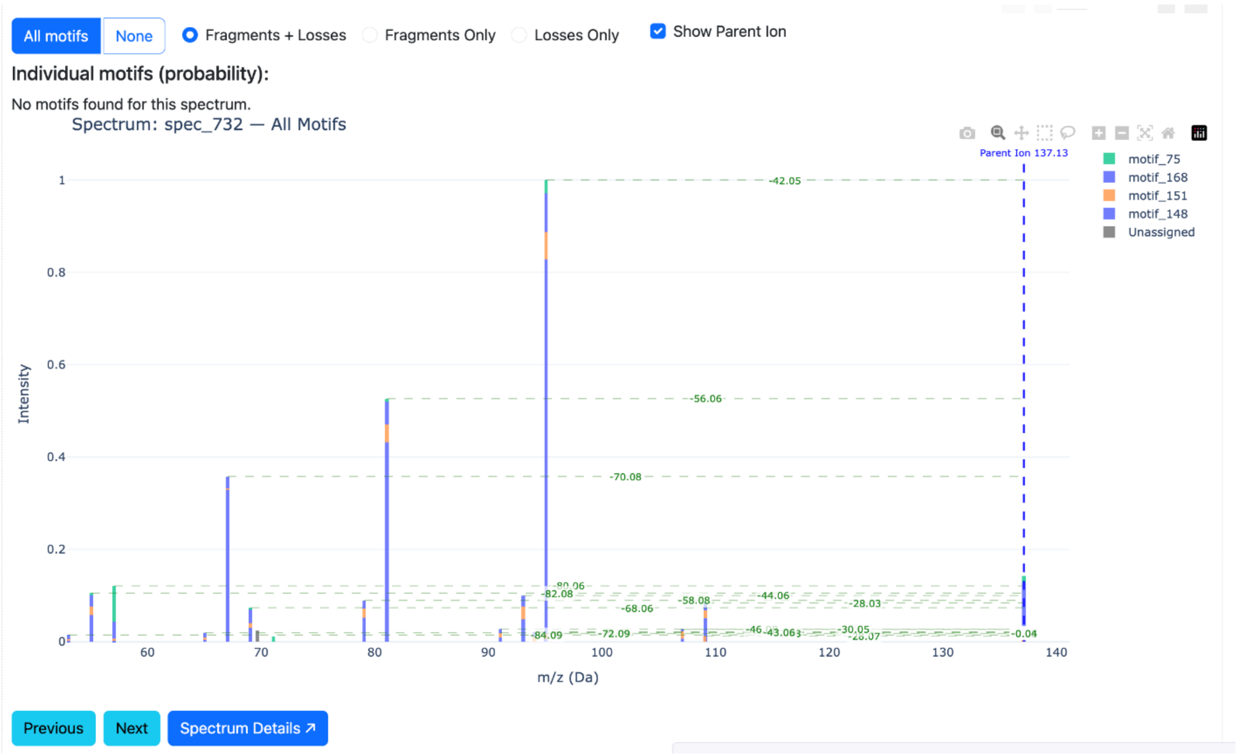

**Fig. S39.** *Mass2Motif exploration interface in the "Motif Details" tab of the application. Below in this tab, MS/MS spectra can be explored with their assigned Mass2Motifs.*

#### 6.3 Spectra Search

The "Spectra Search" screen enables rapid retrieval and detailed exploration of spectra based on specified fragments, losses, or parent masses. Users can input fragment or loss patterns (e.g., frag@150, loss@40) or select a mass range for the precursor mass via the interactive slider (Fig. S40). Matched spectra are displayed in a searchable table, providing direct links to individual spectra. Selecting a specific spectrum reveals an interactive plot underneath the table, highlighting the fragment and loss peaks and their assignments to Mass2Motifs (like the spectrum plot in the Motif Details screen). Peaks color-coded by associated motifs clearly demonstrate how a single spectrum can be explained by multiple co-occurring Mass2Motifs, a distinctive feature of the MS2LDA framework.

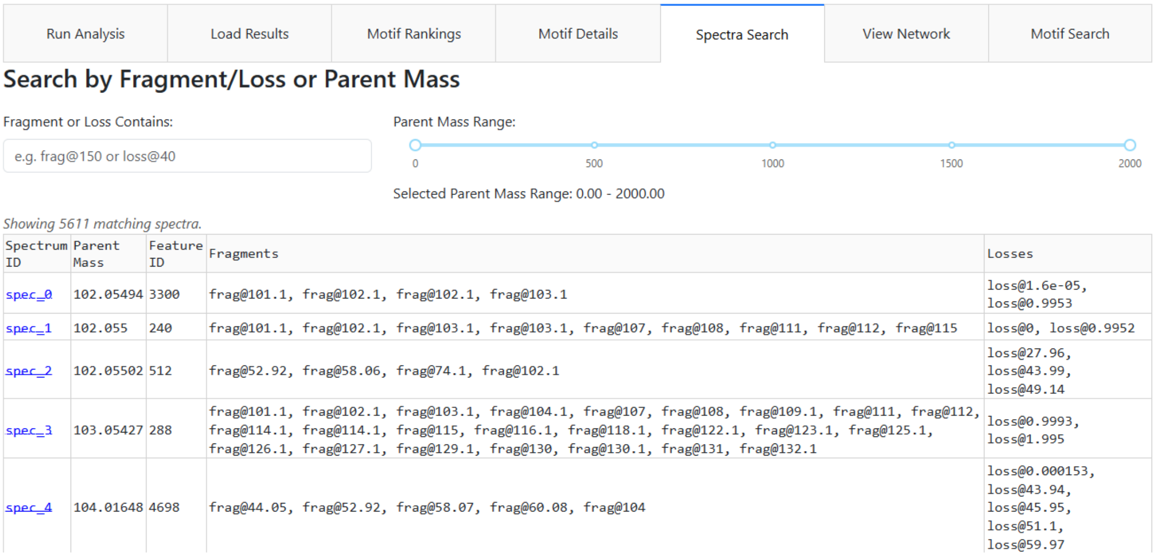

**Fig. S40.** *"Spectra Search" section shown from the application. In this section, is possible to filtering spectra by parent mass range, by fragment and loss contained.*

#### 6.4 View Network

A "View Network" screen shows an interactive network of optimized motifs (Supplementary Notes 1.2) . Each Mass2Motif is displayed as a node (blue), and its fragments (green) or losses (yellow) appear as connected nodes (Fig. S41). Only edges above the selected intensity threshold will be shown. This can be customized using the slider. By default, the extra edges from each loss node to its corresponding fragment node are hidden for less clutter, but the user can re-enable them using the checkbox. By selecting any Mass2Motif, the associated MAG recommendations on the right side are shown. The user can select among multiple layouts, such as "Force-Directed", for clustering the single nodes aside, "CoSE" for a similar layout as used in molecular networking, as "Circle" for checking all the connections between nodes and "Concentric", for grouping the single nodes with no connections as an outer circle, while the nodes with connections are in a small circle inside.

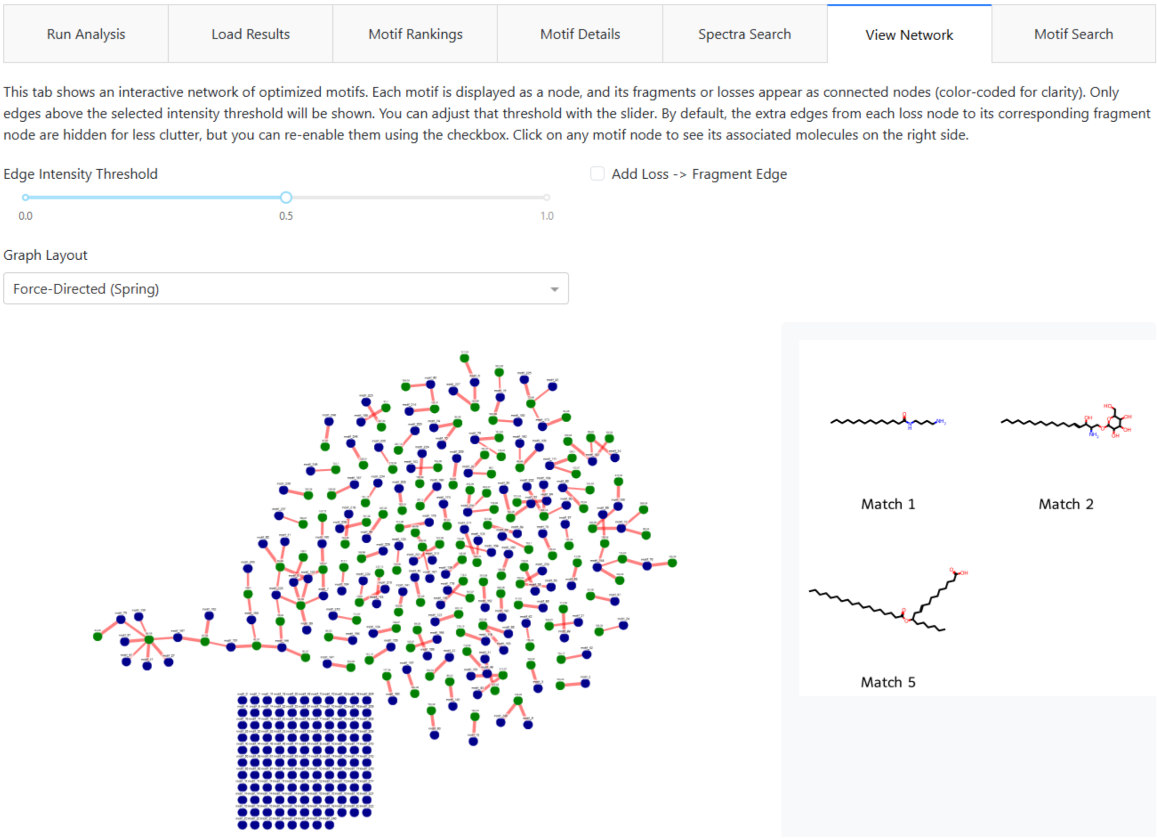

**Fig. S41.** *"View Network" section shown from the application. This section allows to visualize the Mass2Motifs inferred using different layouts (Graph Layout), for example "Force-Directed". When a node is selected, the MAG recommendations are shown. The blue nodes represent the Mass2Motifs, green nodes represent fragments and yellow nodes represent losses. The network can be customized by changing the edge intensity threshold.*

#### 6.5 Motif Search

The "Motif Search" screen facilitates automated annotation by enabling comparison of user-generated Mass2Motifs against selected reference motif sets from MotifDB. Unlike the Motif Annotation Guidance (MAG), which retrieves candidate annotations directly from spectral embeddings of general compound libraries, this approach specifically compares newly inferred motifs against previously annotated motif sets in MotifDB. Users select one or multiple reference MotifSets, run similarity calculations using their Spec2Vec embedding, and visualize matching results. Matches are displayed in a sortable table, showing user motifs, corresponding annotated motifs from reference databases, and similarity scores (Fig. S42). An interactive slider allows dynamic filtering based on similarity thresholds, efficiently narrowing down relevant annotations. This targeted comparison accelerates structural interpretation by linking motifs directly to community-curated annotated substructures in MotifDB MotifSets.

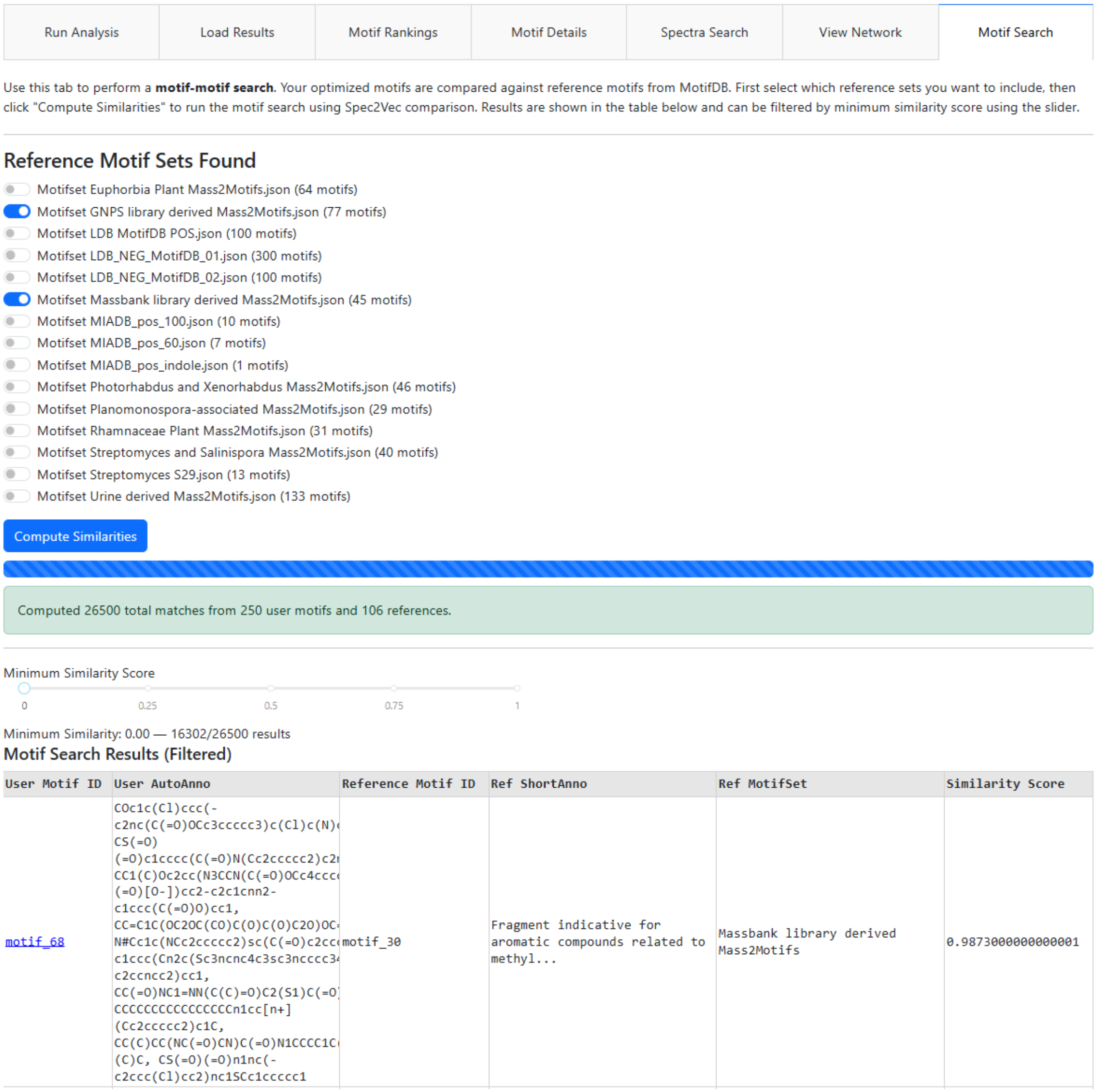

**Fig. S42**. *“Motif Search” section shown from the application. This section enables to compute the Mass2Motif-matching, by computing Spec2Vec similarities between the inferred Mass2Motifs and MotifSets present in MotifDB. On top a selection of different MotifSets is given. After computing similarities, a table of results is shown. The results can be filtered using the Minimum Similarity Score sliding bar. In the results table, information about the Mass2Motif from the dataset and the Mass2Motif belonging to the MotifSet is shown plus their similarity score.*

### 7.MS2LDA 2.0 convergence curves

To measure how well the LDA models converged, MS2LDA 2.0 outputs for every run a convergence curve showing the log-likelihood, perplexity, topic entropy, and document entropy score. In the Fig. S43-S45 the convergence for all case studies are shown. For all case studies the perplexity score was used for early stopping as described in the Method section "Convergence Criteria".

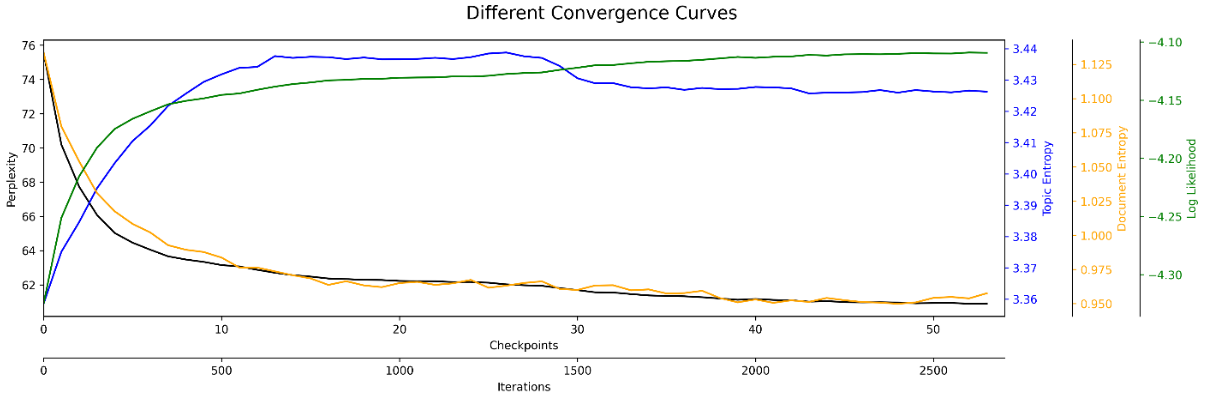

**Fig. S43**. *Convergence curves that were obtained for the case study on pesticides. On the y-axis are plotted: Perplexity, topic entropy, document entropy and log likelihood. On the x-axis checkpoints and iterations (1 checkpoint = 50 iterations). This shows show the evolution of model performance metrics across iterations. Perplexity and document entropy decrease and stabilize after approximately 1800 iterations, indicating improved model fit and reduced uncertainty in document-topic assignments. In contrast, topic entropy and log-likelihood increase initially and likewise reach a plateau beyond 1800 iterations, reflecting improved topic diversity and overall model optimization. These trends collectively indicate stable convergence of the model.*

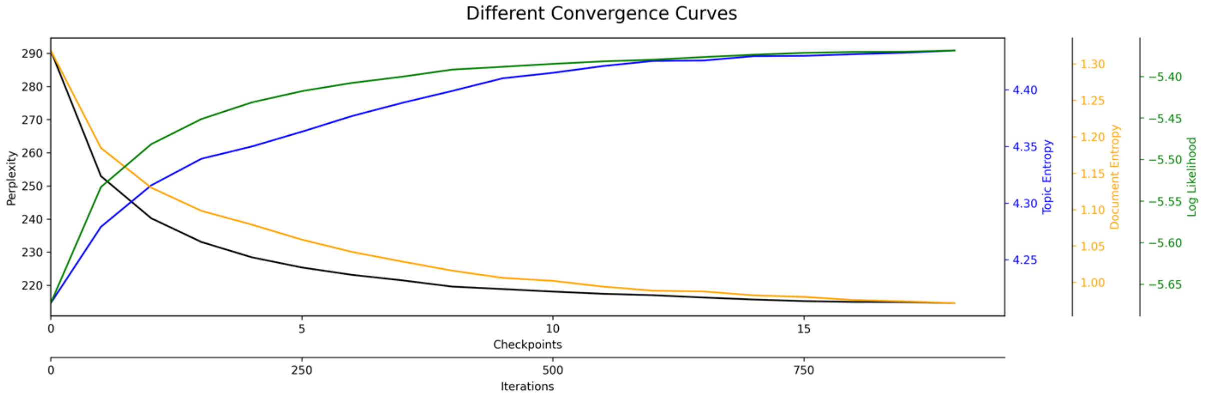

**Fig. S44.** *Convergence curves that were obtained for the case study on pesticides. On the y-axis are plotted: Perplexity, topic entropy, document entropy and log likelihood. On the x-axis checkpoints and iterations (1 checkpoint = 50 iterations). This shows show the evolution of model performance metrics across iterations. Perplexity and document entropy decrease and stabilize after approximately 750 iterations, indicating improved model fit and reduced uncertainty in document-topic assignments. In contrast, topic entropy and log-likelihood increase initially and likewise reach a plateau beyond 750 iterations, reflecting improved topic diversity and overall model optimization. These trends collectively indicate stable convergence of the model.*

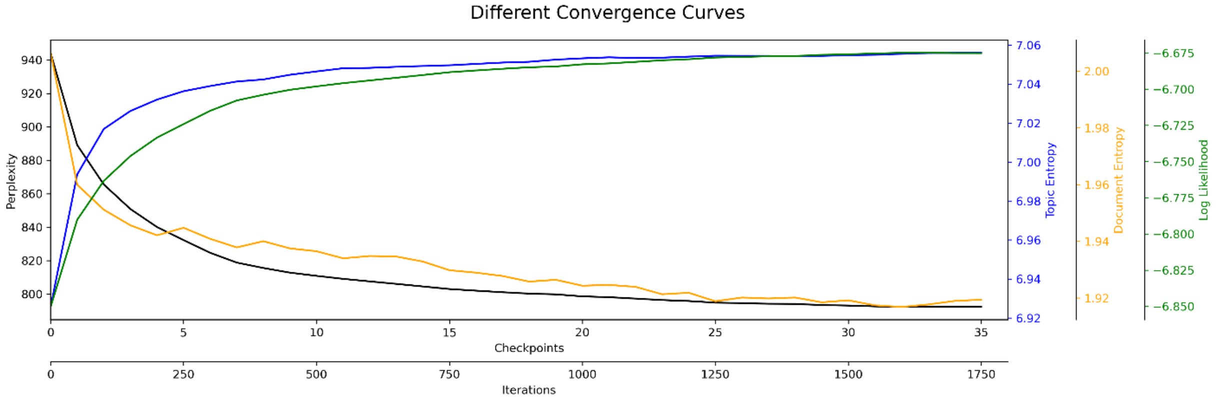

**Fig. S45.** *Convergence curves that were obtained for the case study on pesticides. On the y-axis are plotted: Perplexity, topic entropy, document entropy and log likelihood. On the x-axis checkpoints and iterations (1 checkpoint = 50 iterations). This shows show the evolution of model performance metrics across iterations. Perplexity and document entropy decrease and stabilize after approximately 1500 iterations, indicating improved model fit and reduced uncertainty in document-topic assignments. In contrast, topic entropy and log-likelihood increase initially and likewise reach a plateau beyond 1500 iterations, reflecting improved topic diversity and overall model optimization. These trends collectively indicate stable convergence of the model.*

### 8. MS2LDA 2.0 parameter selection

To guarantee the best modelling outcome and reproducible results when using MS2LDA 2.0 the selected parameter for preprocessing, annotation, and the LDA modelling play a key role. The parameters for all case studies and the MAG benchmarking are given below. Since we used the multi-core option for the LDA modelling (no seed can be set), the results generated for the manuscript can change when run on a different device.

**Parameter selection for the MAG benchmarking.**

```
annotation_parameters = {
    "criterium": "best",
    "cosine_similarity": 0.90,
    "n_mols_retrieved": 5,
    "s2v_model_path": "...",
    "s2v_library_embeddings": “...",
    "s2v_library_db": "...",
}
```

**Parameter selection for Case Study Pesticides.**

```
preprocessing_parameters = {
    "min_mz": 0,
    "max_mz": 1000,
    "max_frags": 1000,
    "min_frags": 4,
    "min_intensity": 0.01,
    "max_intensity": 1
}
convergence parameters = {
    "step_size": 50,
    "window_size": 10,
    "threshold": 0.001,
    "type": "perplexity_history"
}
annotation_parameters = {
    "criterium": "biggest",
    "cosine_similarity": 0.75,
    "n_mols_retrieved": 10,
    "s2v_model_path": "...",
    "s2v_library_embeddings": "...”,
    "s2v_library_db": "...",
}
n_motifs = 250
n_iterations = 5000
model_parameters = {
```

```
    "rm_top": 0,
    "min_cf": 0,
    "min_df": 3,
    "alpha": 0.4,
    "eta": 0.01,
    "seed": 42,
}
train_parameters = {
    "parallel": 3,
    "workers": 0,
}
dataset_parameters = {
    "acquisition_type": "DDA",
    "significant_digits": 3,
    "charge": 1,
    "name": "...",
    "output_folder": "...",
}
fingerprint_parameters = {
    "fp_type": "maccs",
    "threshold": 0.8,
}
motif_parameter = 10
```

**Parameter selection for Case Study Natural Products**

```
preprocessing_parameters = {
    "min_mz": 0,
    "max_mz": 2000,
    "max_frags": 1000,
    "min_frags": 5,
    "min_intensity": 0.01,
    "max_intensity": 1
}

convergence_parameters = {
    "step_size": 50,
    "window_size": 10,
    "threshold": 0.005,
    "type": "perplexity_history"
}

annotation_parameters = {
    "criterium": "biggest",
    "cosine_similarity": 0.7,
    "n_mols_retrieved": 10,
```

```
    "s2v_model_path": “...",
    "s2v_library_embeddings": “...",
    "s2v_library_db": “...",
}

n_motifs = 200
n_iterations = 10000

model_parameters = {
    "rm_top": 0,
    "min_cf": 0,
    "min_df": 3,
    "alpha": 0.6
    "eta": 0.01,
    "seed": 42,
}

train_parameters = {
    "parallel": 3,
    "workers": 0,
}

dataset_parameters = {
    "acquisition_type": "DDA",
    "charge": 1,
    "significant_digits": 2,
    "name": "CaseStudy_mushrooms",
    "output_folder": f"CaseStudy_Mushrooms_{n_motifs}",
}

fingerprint_parameters = {
    "fp_type": "maccs",
    "threshold": 0.8,
}

motif_parameter = 50
```

**Parameter selection for Case Study Suspect list.**

```
preprocessing_parameters = {
    "min_mz": 0,
    "max_mz": 1000,
    "max_frags": 1000,
    "min_frags": 3,
    "min_intensity": 0.01,
```

```
    "max_intensity": 1
}
convergence_parameters = {
    "step_size": 50,
    "window_size": 10,
    "threshold": 0.001,
    "type": "perplexity_history"
}
annotation_parameters = {
    "criterium": "best",
    "cosine_similarity": 0.70,
    "n_mols_retrieved": 10,
    "s2v_model_path": “...",
    "s2v_library_embeddings": "...",
    "s2v_library_db": "...",
}
n_motifs = 1500
n_iterations = 5000
model_parameters = {
    "rm_top": 0,
    "min_cf": 0,
    "min_df": 3,
    "alpha": 0.6,
    "eta": 0.1,
    "seed": 42,
}
train_parameters = {
    "parallel": 3,
    "workers": 0,
}
dataset_parameters = {
    "acquisition_type": "DDA",
    "significant_digits": 2,
    "charge": 1,
    "name": "DDA-Suspectlist",
    "output_folder": f"CaseStudy_Suspectlist_{n_motifs}motifs_output",
}
fingerprint_parameters = {
    "fp_type": "maccs",
    "threshold": 0.8,
}
motif_parameter = 15
```
